# TrueSpot: A robust automated tool for quantifying signal puncta in fluorescent imaging

**DOI:** 10.1101/2025.01.10.632467

**Authors:** Blythe G. Hospelhorn, Benjamin K. Kesler, Hossein Jashnsaz, Gregor Neuert

## Abstract

Characterizing the movement of biomolecules in single cells quantitatively is essential to understanding fundamental biological mechanisms. RNA fluorescent in situ hybridization (RNA-FISH) is a technique for visualizing RNA in fixed cells using fluorescent probes. Automated processing of the resulting images is essential for large datasets. Here we demonstrate that our RNA-FISH image processing tool, TrueSpot, is useful for automatically detecting the locations of RNA at single molecule resolution. TrueSpot also performs well on images with immunofluorescent (IF) and GFP tagged clustered protein targets. Additionally, we show that our 3D spot detection approach substantially outperforms current 2D spot detection algorithms.

## Background

Fundamental biological processes within single cells can result in the formation of localized clusters of macromolecules such as RNA at transcription sites and proteins in phase separated compartments or stress granules^1–5^. These clusters can be visualized with fluorescently labeled probes and imaged with a fluorescence microscope, resulting in large data sets that need to be quantified to extract meaningful biological insights.Manual or semi-automated quantification of such images is labor intensive, biased, and difficult to reproduce, often limiting the interpretable information that can be gleaned. To overcome these limitations, an automated, minimally biased, and robust image processing software tool is needed.

One common experimental approach focuses on the quantification of RNA molecules in fixed single cells. Single molecule RNA fluorescent in situ hybridization (smRNA-FISH) is a technique utilizing fluorescently tagged oligonucleotide probes to visualize target RNA molecules as defraction limited single spots in 3 dimensions^1,6–10^. Such experiments produce large volumes of image data, especially in cases where the target is tracked or quantified over many time points.

Several tools exist that are capable of detecting and quantifying RNA-FISH signal from raw image stacks using a variety of strategies and implementations. However, many prominent tools such as FISH-Quant v1^11,12^, Airlocalize^13^, Starfish^14^, dNEMO^15^, and RS-FISH^16^ are manually oriented or require at least one manual step. Others, such as Big-FISH (the core of FISH-Quant v2)^17^, and deepBlink^18^, claim to be fully automated. However, we have found that in practice they require manual parameter adjustment and human feedback to perform well on a wide variety of experimental images.

Additionally, current deep learning-based tools such as deepBlink, Piscis^19^, Spotiflow^20^, and Polaris^21^ only operate on 2D images instead of directly on 3D image stacks. This puts an additional requirement on the user to either generate a maximum intensity projection from a stack, which may not be entirely representative of the original 3D image, or process each slice individually and re-merge the resulting call sets. Both of these options introduce additional steps and opportunity for error.

A particular step that has traditionally been a challenge to automate is the selection of an intensity threshold that filters out local maxima below a certain intensity value, affecting which of the brighter voxels get called by the software as signal spots^7,17,18,22^. Thresholds that are set too low result in a high number of false positives, defined as points that are identified as signal but are not real signal. Whereas thresholds that are set too high filter out many true positives-points that are correctly identified as signal spots. Plotting the number of spots detected against the threshold for a variety of image types usually yields a curve that starts with a steep decline followed by a level-off (Figures 1A-I, 2). The point where the level-off begins is usually a good threshold choice^7,17,22^.

**Figure 1.**
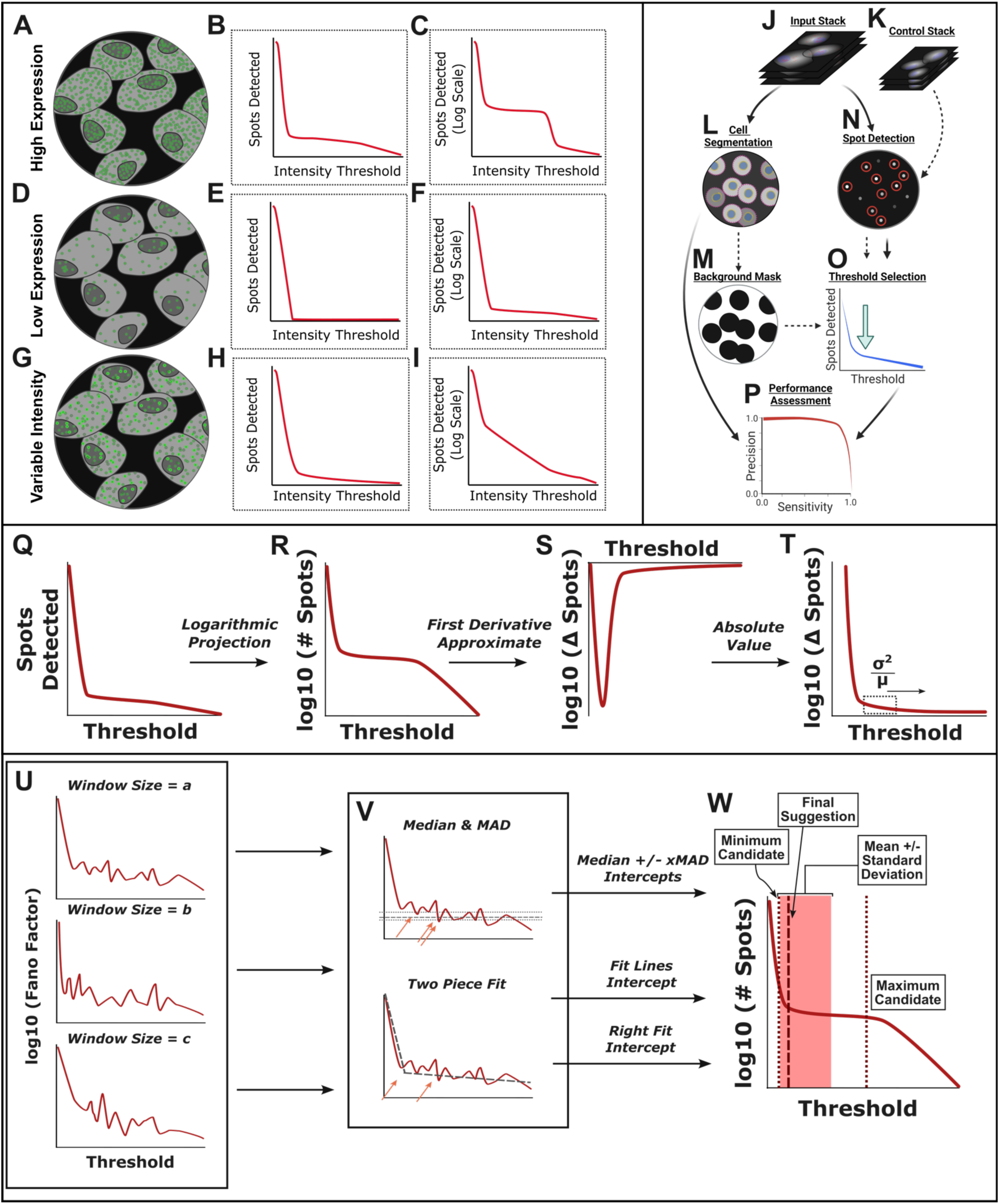
TrueSpot uses multiple approaches to generate a threshold range, allowing it to adapt to different curve shapes. In a High Expression image (A-C), there is a large number of signal puncta at around the same intensity (A) and the spot count curve has a distinct plateau well above the zero signal level(B,C). Low Expression images (D-F) have far fewer puncta, often around the same intensity level (D). Variable Intensity images (G-I) are characterized by the presence of distinct signal puncta that have a wide range of intensities (G). TrueSpot’s maxima detection pipeline is depicted in (J-P). An input image stack and optional control image (J-K) are run through initial spot detection using a Laplacian of Gaussian approach and local maxima are filtered by sweeping across a range of intensity threshold values (N). Automatic threshold selection (O) is then performed by searching for a relative flat region in the spot count curve as detailed in (Q-W). Additionally providing a control image stack or cell segmentation mask (L) to extract the image background (M) can assist threshold selection by setting a signal floor. Tool performance can be assessed against reference call sets using measures of recall (Equation 1A) and precision (Equation 1B). (W) A pool of threshold suggestions are then derived from the steps described in (Q-V), and a single suggestion is made using a tunable weighted average of the pooled values. The dashed vertical line denotes the returned threshold value, shaded rectangle the mean ± standard deviation, and dotted lines the minimum and maximum extremes of threshold suggestion pool.

However, not all curves are shaped quite the same, and a tool that is to consistently select a threshold that maximizes call accuracy must have an algorithm that can flexibly handle this variability so that it can be used on many different kinds of images with different targets, probes, and levels of expression. The assumption typically made when using spot count curves to find noise-signal boundaries is that a curve will have a clear plateau region between two drops, but this generally only occurs when the target expression is high and uniform (Figures 1A-C, 2A-E). When expression is low, the curve will usually be ‘L’ shaped with no sharp second drop, and often the right arm of the ‘L’ will not be completely flat (Figures 1D-F, 2F-J). Another common occurrence in experimental images is heterogeneity in intensity within the signal region, or an overlap between signal and noise. These “variable intensity” images tend to produce curves with very gradual bends instead of sharp ‘L’ shapes. These bends can sometimes be so gradual as to appear diagonal on a logarithmic projection (Figures 1G-I, 2K-O).

We have developed TrueSpot, a software tool that automates spot detection, threshold selection, and quantification (Figure 1J-P). We tested TrueSpot against three other tools using three different approaches to RNA-FISH signal quantification: Big-FISH^17^, RS-FISH^16^, and deepBlink^18^. We chose these tools to test against because they either had the ability to bypass manual thresholding (Big-FISH, deepBlink) or utilized a different approach to spot localization (deepBlink, RS-FISH).

We found that TrueSpot overall outperformed the other tools while requiring less manual effort and behaving more consistently with its threshold selections (Figures 3-6). Overall, we believe TrueSpot will be a useful addition to the computational toolbox of laboratories that use RNA-FISH and similar imaging techniques to quantify localized signal, addressing serious deficiencies and overcoming limitations of existing tools, while increasing performance consistency across large image batches and reducing human-induced subjectivity.

## Results

### TrueSpot uses an automated threshold selection algorithm that utilizes multiple approaches

TrueSpot accepts either a 2D image or 3D image stack (which may or may not include multiplexed channels) as input along with an optional control image (Figure 1J-K). The sample and control are then passed through the initial spot detection module (Figure 1N). Here, a Laplacian of Gaussian (LoG) filter^6,7^ is applied to the image and coordinates of local intensity maxima in the filtered image are recorded. A range of threshold values is scanned to determine which identified maxima are dropped at each tested threshold. The number of spots/maxima retained at each threshold in the sample image is then plotted against the threshold value as the first step in automatic threshold selection (Figure 1O). If a control image is not provided, a background mask can be derived from a cell segmentation mask (our cell segmentation module is discussed in a previous publication^23^) and the masked image can be used as a control instead (Figure 1L-M). The use of controls is entirely optional - their purpose is to assist the threshold selector by finding a noise ceiling. This functionality is however not required to elicit high performance from the threshold selection algorithm, as it was not utilized in any of our simulated image tests, nor in many of the experimental image groups (Figure 3).

Performance is assessed using recall (also called “sensitivity”) (Equation 1A) and precision (Equation 1B) as benchmarking measures across all thresholds, at the selected threshold, and in single cells (Figure 1P). Area under the precision-recall curve (PR-AUC) was used to evaluate overall detection performance across all tested thresholds. The maximum recall (highest value for recall reached for a tool on a given image across all tested thresholds) was used to distinguish between relative precision and recall contributions to PR-AUC. We used the F-Score (Equation 1C), a quantification of the trade-off between precision and recall, at auto-selected threshold values to assess tools’ ability to automatically select appropriate thresholds.

#### Equation 1. Performance Metrics

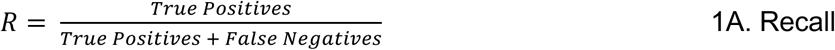

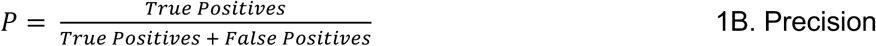

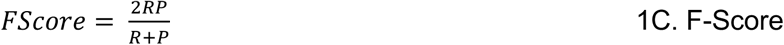

Our threshold selection module operates by combining two strategies and sweeping over a range of parameter values to produce a pool of potential threshold values, then combining these threshold suggestions into a single suggestion which can then be applied to the set of initial spot calls (Figure 1Q-W). Importantly, while both the range suggestion and single threshold selection procedures are tunable, the default parameters work well for a variety of image types (Figure 3) and tuning is generally not necessary.

TrueSpot ‘s threshold selection module performs a transformation on input spot count curves (Figure 1Q) by default as a normalization measure instead of operating directly on the original curves. (Curves in Figure 1R-W are shown as base ten logarithmic projections to better emphasize the region of interest (Figure 1C,F,I). However, the software operates on the original curve in the steps illustrated in Figure 1R-T).

First, the first derivative approximate (difference between adjacent data points) of the spot count curve is taken (Figure 1S). Since spot count only decreases or remains the same as threshold increases, the first derivative approximate is expected to always be negative or zero. Thus for simplicity, its absolute value is used instead. Additionally, threshold values below that correlating to the peak magnitude in spot count difference are trimmed out to ensure that the transformed curve still begins at its highest point (Figure 1T).

Finally, the Fano Factor (Equation 2) of the values within a sliding window^7,22^ across this projection is calculated (Figure 1T-U). This provides a measure of the local curve topology better normalized to the magnitude of spots counted. The purpose of these transformation steps is to improve threshold selection consistency across images with vastly different properties.

#### Equation 2. Fano Factor

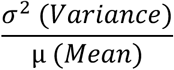

The size of the window across which the Fano Factor is calculated is adjustable, and the default settings accumulate threshold suggestions across a range of window sizes (Figure 1U). Two operations are performed on the log10 projection of a transformed curve to generate a pool of threshold selections (Figure 1V). First, the median and median absolute deviation (MAD) of the y values (ie. transformed spot counts) are determined. The lowest x (threshold) value with y values at or below the median added to a constant multiplied by the MAD is recorded as a threshold suggestion. If multiple values are provided for the constant MAD factor, all will be tried and their threshold suggestions recorded (Figure 1V). This provides an initial threshold range which can then be used to aid the second operation, a simple two-piece linear fit of the ‘L’ shaped transformed curve. The x coordinate of the piece intercept and the x coordinate of the leftmost (lowest x) intersection of the right-hand fit line and the original curve are also recorded as threshold suggestions (Figure 1V). This pool of median and fit based threshold suggestions is then combined using a weighted average to output a single suggestion (Figure 1W).

### TrueSpot and three other spot detection tools were benchmarked on simulated and experimental images

To evaluate the performance of TrueSpot and compare it to similar puncta identification and quantification tools on a variety of images with different parameters, we performed benchmarking against Big-FISH^17^, the python core of FISH-Quant v2 which uses a similar Laplacian of Gaussian (LoG) approach as TrueSpot and also provides an interface for automated threshold selection; RS-FISH^16^, a Java-based spot caller that utilizes a radial symmetry approach and provides an interactive graphical user interface (GUI) as a plugin for the image viewer Fiji; and deepBlink^18^, a deep learning based tool that detects image features based on the provided model.

For the purpose of having genuine ground truths to benchmark against, we generated 1555 simulated images using the Sim-FISH tool^17^. In addition, we pulled the set of 42 publicly available and externally generated simulated images used previously to test RS-FISH^16^.

As it is difficult to truly replicate the realistic noisiness of experimental imaging with simulated images (supplementary results, Figures S1-S5), we also tested TrueSpot, Big-FISH, RS-FISH, and deepBlink on the 674 images from our previously published RNA-FISH time course data on salt stimulated *Saccharomyces cerevisiae*^24^ and 195 miscellaneous experimental images from our lab^25^, (Unpublished data), collaborators^26^, and the RS-FISH benchmarking set^16^.

### Experimental spot count curves are highly variable with many falling into three archetypes

Experimental images produce spot count versus threshold curves with properties sufficiently variable to make designing an automated threshold selection algorithm challenging. We observed three common archetypes: low expression, high expression, and variable intensity (Figure 1A-I). Experimental examples of each can be seen in Figure 2.

**Figure 2.**
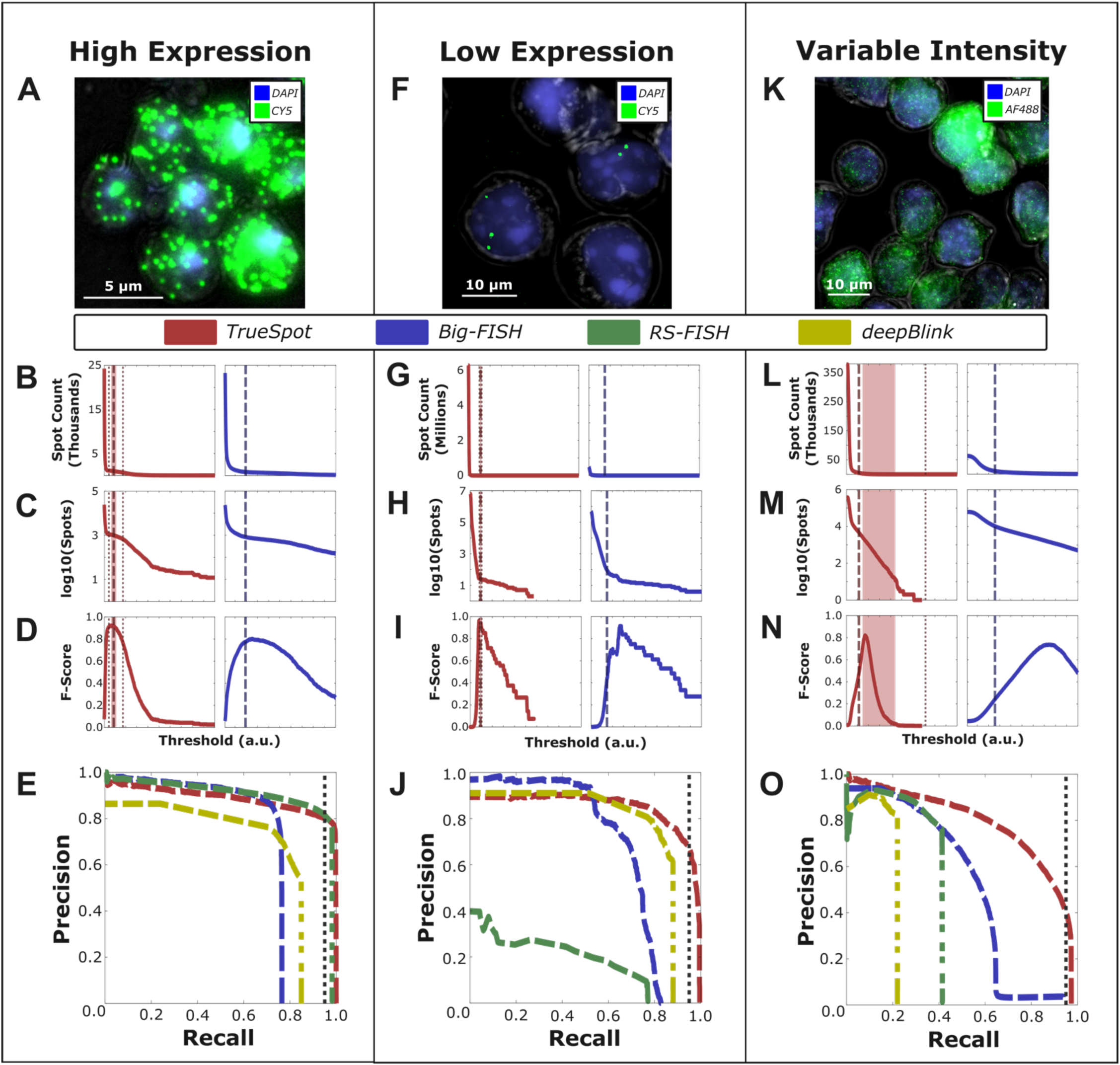
Experimental images produce a variety of signal curves. Example image samples with broad spectrum of expression. (A,F,K) White light (grey), DAPI stained nucleus (blue), and target channels (green, yellow) in representative sample images, (B,G,L) spot count (linear and (C,H,M) logarithmic scale), (D,I,N) F-Score curves for tools with automated threshold selection (TrueSpot in red, Big-FISH in blue -RS-FISH and deepBlink omitted as they do not have automated threshold selection), and (E,J,O) composite precision-recall curves (all four tools – TrueSpot (red), Big-FISH (blue), RS-FISH (green), and deepBlink using stock smFISHmodel (yellow)) for the example groups (CTT1-CY5, XistExonic, and H3K36me3-AF488 respectively) used to illustrate each of the three curve archetypes: (A-E) high expression, (F-J) low expression, and (K-O) variable intensity. Dashed lines in (B-D,G-I,L-O) denote automatically selected threshold. Dotted lines denote threshold pool minimum and maximums. Shaded region denotes mean of threshold pool ± standard deviation. Vertical dotted line in (E,J,O) marks 95% recall.

We found it difficult to evaluate linear plots of spot count versus threshold due to the extreme order of magnitude variability between the counts at lowest thresholds and the signal range. As a result, we will focus on the logarithmic projections for the purpose of our analyses.

The first archetype is represented in Figure 2A-E. The image in Figure 2A shows highly expressed CTT1 mRNA tagged with CY5 labeled RNA-FISH probes in *S. cerevisiae*^24,27^. There are many distinct puncta in the image all around the same narrow range of intensity. Images like these produce the classic spot count curve type, characterized by a sharp bend followed by a signal plateau and second dropoff (Figure 2B-C). Both TrueSpot and Big-FISH tended to have the least amount of trouble finding thresholds at peak or near-peak F-Score (Figure 2D), and all four tools tended to display high general performance for images of this type (Figure 2E). Interestingly, Big-FISH had a maximum recall at just under 0.8 (Figure 2E). We noticed that this was a recurring issue with Big-FISH on experimental images, particularly those with densely clustered spots. This could either be the product of a genuine flaw in the tool’s algorithm or simply how Big-FISH handles closely neighboring calls prior to signal quantification. deepBlink similarly had an unexpectedly low maximum recall. This couldbe due to similar clustering behavior or merging during 2D to 3D conversion.Another possibility is that this is the productofan absence of permissive threshold options resulting in the lack of large call sets produced by oversensitivity.

Figure 2F-J show a representative example of the second archetype, characterized by few sparse, sometimes extremely sparse puncta as can be seen in the image in Figure 2F of CY5 labeled RNA-FISH probes targeting the long non-coding RNA (lncRNA) Xist^25^. This archetype’s spot count curve is characterized by a single sharp bend, and lack of a second falloff and preceding plateau (Figure 2G-H). These curves are usually ‘L’ shaped, though the lower arm may not be completely flat. TrueSpot was initially designed to handle this type of image, and in this example it indeed performed quite well, cleanly identifying a threshold at or near the peak F-Score (Figure 2I, left). In contrast, while it did not perform terribly, Big-FISH selected a threshold that was on the sensitive side of optimal (Figure 2I, right). General detection performance could be variable for images in this category on the part of all four tools, however in the case of this particular image, TrueSpot, Big-FISH, and deepBlink performed quite well while RS-FISH appeared to struggle immensely.

The final example in Figure 2K-O illustrates a case where signal intensity is highly variable and at times difficult to distinguish from background. Of note, the image in Figure 2K was produced from an immunofluorescence(IF) experiment probing histone mark H3K36me3 rather than an RNA-FISH experiment. Still, as it is not uncommon for similar attributes to be present in FISH images, the challenges that arise when attempting to extract quantitative information from images of this archetype are still applicable.

Figure 2K shows many small, densely clustered signal puncta, some of which are quite dim and some very bright. Images like these are difficult to threshold, even manually, as one individual may have a different opinion from another individual regarding at what level spots are too dim to consider signal. The resulting spot count curves usually have very gradual falls (Figure 2L), not uncommonly appearing outright diagonal on a logarithmic projection (Figure 2M). Both TrueSpot and Big-FISH picked threshold values that produced call sets in poor agreement with the manually generated reference set and tended to struggle with threshold selection in these variable intensity groups (Figure 2N). The PR curves in Figure 2O, generated from the composite spot counts from all of the curated H3K36me3 images, indicate a trend in all four tools of high maximum precision under restrictive parameters, but highly variable maximum recall and PR-AUC. TrueSpot performed the best overall. Big-FISH’s maximum recall is over 90%, but the threshold values yielding recall values above around 65% appear to have incredibly low precision. RS-FISH and deepBlink had maximum recalls of just above 40% and 20% respectively. These drop-offs are very sudden, and it is possible that higher recall values may be obtained by testing more permissive thresholds (or, in deepBlink’s case, lower call probabilities). However, such difference of Gaussian (DoG) threshold values or call probabilities well below the range of those we tested would be so low (≤ 0.0036 RS-FISH, < 1% deepBlink) as to be impractical for regular use.

The archetypes shown in Figure 2 are not necessarily members of discrete groups. Rather, it is more useful to think of these examples as extremes on a spectrum of curve shapes resulting from different target expression levels and signal intensity ranges.

### TrueSpot and Big-FISH show the strongest overall performance on simulated images

To assess the general spot detection performance of each tool across a wide range of threshold values, we used the PR-AUC as a performance metric. Distributions of PR-AUC values calculated for the set of simulated images for all four tools are shown in Figure 3A.

**Figure 3.**
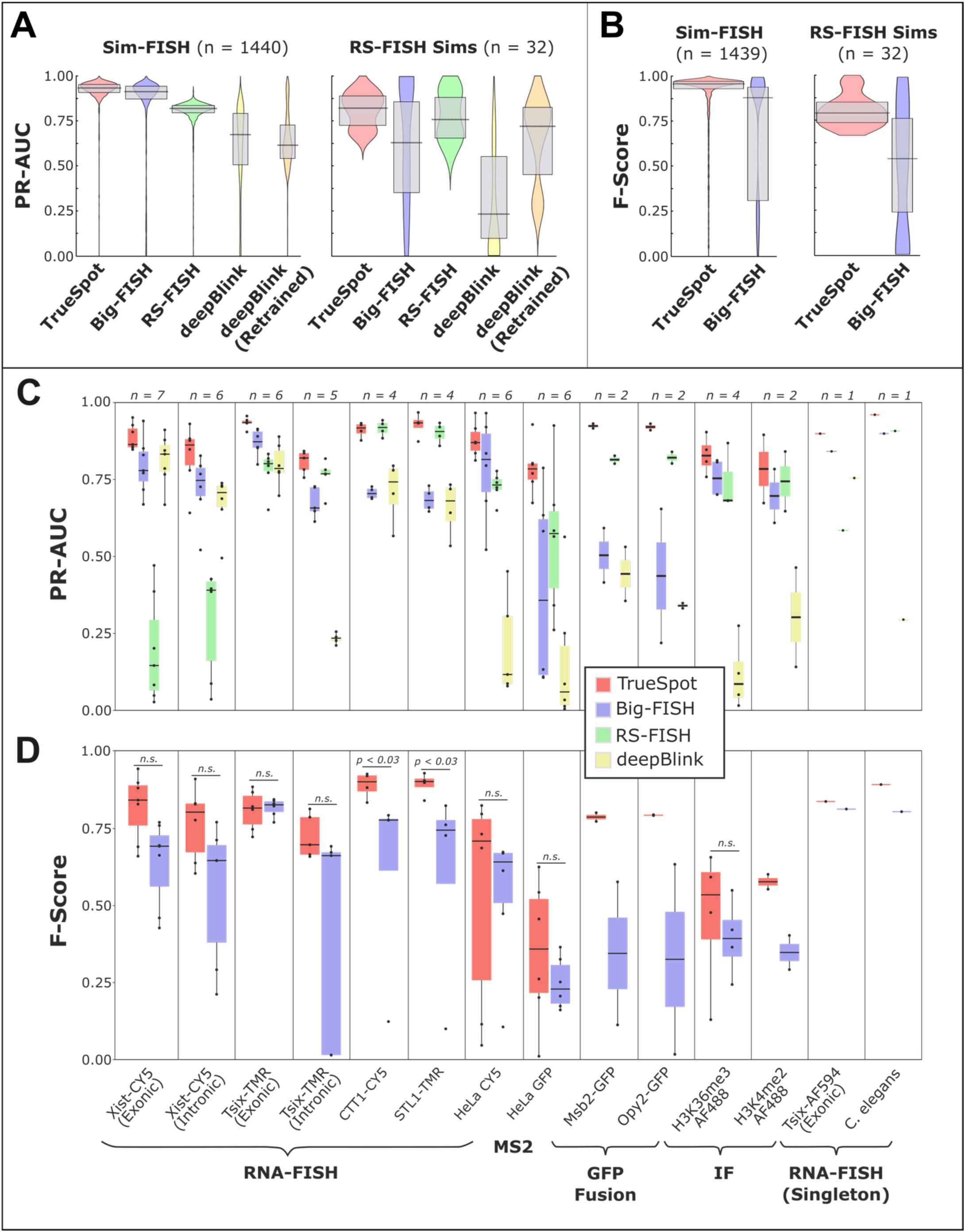
TrueSpot performs well on simulated and experimental images. (A) Distribution of PR-AUC values for both sets of simulated images (zero voxel proportion ≤ 0.7), all four tools and deepBlink with model retrained on simulated data subset (orange) (Sim-FISH n = 1337, RS-FISH set n = 32). PR-AUC is used as a metric for overall detection performance across all threshold values tested. 25^th^ and 75^th^ percentiles are marked by grey box, median is marked by horizontal black line. Statistical calculations for PR-AUC in simulated image sets can be found in Table S1B. (B) Distribution of F-Score values at automatically selected thresholds for both sets of simulated images (zero voxel proportion ≤ 0.7). Only tools with automatic threshold selection were included (TrueSpot in red, Big-FISH in blue). 25^th^ and 75^th^ percentiles are marked by grey box, median is marked by horizontal black line. Statistical calculations for F-Score in simulated image sets can be found in Table S1C. (C) Distribution of PR-AUC scores across 14 groups (9 batches, 3 pairs, 2 singletons) of experimental images for all four tested tools. Box boundaries represent 25^th^ and 75^th^ percentiles, midline represents median, and whiskers represent furthest values within 1.5 * interquartile range (IQR) from box edges. *n* values at the top are the number of image channels in each set. Statistical calculations for PR-AUC in experimental image sets can be found in Table S2B. (D) Distribution of F-Scores at automatically selected threshold across 14 groups of experimental images for tools with automated threshold selection (TrueSpot in red, Big-FISH in blue). Statistical calculations for F-Score in experimental image sets can be found in Table S2C.

TrueSpot and Big-FISH performed the strongest, with tight clusters of PR-AUC values at or above 0.85 across all Sim-FISH generated images (Figure 3A, Left) (TrueSpot: mean = 0.926, median = 0.939; Big-FISH: mean = 0.847, median = 0.916; Mann-Whitney p = 3.41 x 10^−39^). Both tools performed worse across the RS-FISH simulated image set (TrueSpot: mean = 0.812, median = 0.838; Big-FISH: mean = 0.683, median = 0.883; Mann-Whitney p = 0.898), though this effect was more pronounced with Big-FISH than with TrueSpot (Figure 3A, Right).

RS-FISH performed consistently well on the Sim-FISH images (mean = 0.804, median = 0.821), though not as well as TrueSpot or Big-FISH (Figure 3A, Left). The likeliest explanation for this is that the fixed batch parameters that RS-FISH requires as input such as the sigma (expected spot size) or z anisotropy factor were suboptimal for most images in the dataset, despite us attempting to tune them manually beforehand. Other possibilities are that there are regions in an image stack that RS-FISH’s detection algorithm does not look at that we were not aware of during testing, or that it is sensitive to certain artificialities present in the simulated images Sim-FISH produces. In contrast, RS-FISH performed on par with Big-FISH and TrueSpot on the RS-FISH simulated dataset (Figure 3A, Right)(mean = 0.781, median = 0.772; Mann-Whitney vs. TrueSpot p = 0.36, vs. Big-FISH p = 0.877).

deepBlink using the default smFISH model was the worst performing tool on both simulated image subsets with middling PR-AUC median values and large well-spread distributions (Sim-FISH set: mean = 0.612, median = 0.672; RS-FISH set: mean = 0.216, median = 0.046). This however can likely be attributed to model choice and/or 2D to 3D coordinate conversion (see methods). For consistency across all tests on experimental and simulated images, we used the stock smFISH model provided by the developers^18^, which was likely trained on experimental or more realistic datasets.

To test the impact of model choice, we trained a new model on a subset of the Sim-FISH simulated images and reran the deepBlink prediction model on the remaining simulated images (Sim-FISH set n = 1337, RS-FISH set n = 32). The results are shown alongside the other tools in Figure 3A. Interestingly, though the training set was comprised of our randomized Sim-FISH images, deepBlink did not perform any better on the remaining Sim-FISH set using the retrained model (mean = 0.646, median = 0.615; Mann-Whitney vs. deepBlink default p = 0.202), yet it performed much better on the RS-FISH set (mean = 0.66, median = 0.767; Mann-Whitney vs. deepBlink default p = 1.99 x 10^−6^).

With the exception ofdeepBlink, the cases where the tools performed particularly poorly (PR-AUC < 0.6) were in the vast minority (TrueSpot: 0.011, Big-FISH: 0.093, RS-FISH: 0.021, deepBlink: 0.37, deepBlink Retrained: 0.372; visual representation for maximum recall and PR-AUC can be seen in Figures S6A-B). Notably, in cases where one tool performed poorly, the other tools often performed poorly as well, indicating that the poor performance was more likely due to the properties of the simulated image itself than flaws in any of the individual tools.

Another interesting observation we made was that recall did not reach 100% for any of the four tools on the simulated dataset very often (TrueSpot: 0.018, Big-FISH: 0.016, RS-FISH: 0.015, deepBlink: 0.00, deepBlink Retrained: 0.0063; Figures S6A, S7A). This is an indication that in even the cleanest simulated images, there was a subset of spots that no tool was able to detect at even the lowest thresholds. There are several possible causes for this independent of any tool’s detection ability. Namely, the presence of particularly dim spots that cannot be distinguished from the background even by manual inspection, the presence of tightly clustered spots that may be called as a single signal nucleated by only the brightest in the cluster, or the presence of spots on the very edges of the stack that fall outside the tools’ detection boundaries or due to being partially or mostly cut off cannot be resolved as puncta.

#### TrueSpot performs auto-thresholding well on simulated images with high background noise

To benchmark the auto-thresholding performance of TrueSpot and Big-FISH, we compared F-Score values at the chosen thresholds across all simulated images (Figure 3B). There was only one image that both tools failed auto thresholding, which was omitted from these figures. Interestingly, there was far less overlap in the subset of individual images with poor F-Scores between the two tools than with PR-AUC or maximum recall (Figure S7).

In both subsets of simulated images, TrueSpot (Sim-FISH: mean = 0.902, median = 0.959; RS-FISH: mean = 0.802, median = 0.799) tended to produce threshold calls that yielded higher F-Scores (Mann-Whitney: Sim-FISH p = 5.28 x10^−72^; RS-FISH p = 1.52 x10^−4^) than Big-FISH (Sim-FISH: mean = 0.729, median = 0.937; RS-FISH: mean = 0.318, median = 0.391) (Figure 3B, statistics in Table S1C). The difference in PR-AUC between the two tools being far less pronounced, which suggests that Big-FISH struggles with automated threshold selection relative to TrueSpotmoreso than with initial spot detection.

To evaluate whether there were specific image attributes that correlated with poor threshold selection for either tool, we examined the distribution of F-Score values versus SNR (Figure S8B). The most striking trend was the increased density of low F-Score callsets from low SNR images by Big-FISH, relative to both TrueSpot and Big-FISH PR-AUC (Figure S8A). This indicates that while Big-FISH tends to struggle slightly with general spot detection for low SNR simulated inputs relative to TrueSpot and RS-FISH, it has even more difficulty with consistent selection of appropriate thresholds for these images.

In contrast, TrueSpot tended to consistently choose good thresholds for images regardless of SNR (Spearman: 0.039, p = 0.123; Examples shown in Figure S4). There is however a small subset of simulated images with low to middling F-Scores which can be seen as faintly lightened regions around the right center of Figure S8B (left). As the PR-AUC distribution for TrueSpot lacks a similar subset of poor performances (Figure S8A), this is indicative of a different, more mild, and seemingly SNR independent flaw in the auto-thresholding process. This is likely tied to the presence of spot count curves that lack a sharp initial dropoff that simulated images with insufficient noise can produce (Figures S2J-L, S3). This is supported by the observation that the prevalence of these low scoring cases drops when the maximum proportion of zero-value voxels drops, and that the proportion of low F-Score cases that have a zero voxel proportion at or above 0.65 (F-Score ≤ 0.7: n = 131; n = 81, 0.618; Figure S5) is higher than the proportion of such images in the full set (n = 438; 0.275).

#### Overall detection performance is variable across experimental images with different properties

To analyze the performance of the four tools on experimental data, we used the same PR-AUC and F-Score metrics we used to evaluate performance on simulated data. However unlike simulated images, experimental images do not have an objective ground truth set to compare against. Instead, we manually created spot call “reference sets” for each evaluated image, either by agnostically choosing individual spots or by using a manually thresholded call set from an outside software program that was not tested here. Thus, these results should be interpreted as a quantitative evaluation of agreement regarding what constitutes a “spot” between tool calls and the human curator.

Experimental images were assessed in groups, determined by the type of target and probe used (Figures 3C-D, S9). Grouping experimental images in this manner allowed us to better evaluate the behavior of the tools on certain types of image. Of our main groups, nine^16,24–26^ were the product of RNA-FISH experiments, three^26^ (Unpublished data) utilized GFP tags that are prone to producing duller and blurrier images, and the remaining two^25^ came from immunofluorescent (IF) tagging of histone markers. Two of the evaluated RNA-FISH classes are singletons, one of which is the *C. elegans* embryo image evaluated in the RS-FISH paper^16^.

In one of the seven RNA-FISH batch groups (TsixExonic), TrueSpot had the significantly highest PR-AUC (Mann-Whitney comparisons and distribution statistics can be found in Table S2B), and in the other six TrueSpot effectively tied for the highest with another tool, with the difference between the tools being nonsignificant. Additionally, TrueSpot produced the highest PR-AUC for both singleton images.

TrueSpot had significantly a higher PR-AUC than all three other tools in the HeLa-GFP group, and scored the best on all four protein-GFP fusion images (Msb2 and Opy2 groups). TrueSpot, Big-FISH, and RS-FISH tied statistically in the larger histone mark IF group (H3K36me3) with a similar pattern being present in the other histone mark group.

TrueSpot also scored the significant highest maximum recall in two of the nine batch groups, and tied for best in the other seven (Figure S9, Table S2A). It also trended the highest, often alongside RS-FISH, in each of the five pairs and singletons. Though PR-AUC tended to be slightly lower than maximum recall for most groups, there did not appear to be any group-specific large discrepancies between the metrics. This suggests that precision may contribute slightly more to limiting overall performance than recall in TrueSpot’s maxima identification process.

Big-FISH generally performed acceptably across most groups, with PR-AUC values tending to fall between 0.6 and 0.8, though there were no groups with manual reference sets where Big-FISH had consistently strong results that set it apart from the other tools. Maximum recall trended around the same regions for RNA-FISH groups, but notably much higher than PR-AUC for the protein groups (Figure S9). This contrast, particularly in the GFP groups, indicates that Big-FISH’s poorer overall performance on these images is also a matter of precision rather than recall.

RS-FISH performed quite well on experimental images. This is particularly fascinating when contrasted to its relatively mediocre performance on simulated images and may indicate a strength in processing more realistic images given well-tuned input parameters. Surprisingly, PR-AUC was consistently higher in the protein GFP fusion pairs than in some other groups that would seem to be better suited for RS-FISH’s design, such as either Xist-CY5 group or Tsix-AF594. However, precision appeared to be a very strong limiting factor for RS-FISH in certain cases. In the aforementioned three mouse lncRNA groups, RS-FISH scored very well in maximum recall, but poorly in PR-AUC. This is odd because these images have large, distinct puncta and the lncRNA groups RS-FISH struggled the most with also tended to have a narrow signal range. This behavior could be the result of low total signal such as the case illustrated in Figure 2A-E, or possibly a hard to detect property of these specific images causing the manual parameters to poorer fit than expected from initial tuning.

Assessing deepBlink is more difficult than the other three tools due to its inability to process images in more than two dimensions. Thus, its apparent poor performance here should not be taken as a broad indication that the tool works poorly. Indeed, despite the potential score depression introduced by suboptimal model selection or flaws inherent in attempting to compare a 2D call set to a 3D truth set, deepBlink performed relatively well using the same developer provided smFISH model that was used for simulated data assessment. In five of the seven RNA-FISH groups, it performed on par with Big-FISH (statistics in Table S2B), and produced a high PR-AUC on the Tsix-AF594 test image (0.0757). Interestingly, when measuring maximum recall, deepBlink performs within the range of at least one other tool in all but one RNA-FISH group (Figure S9, Table S2A), implying that the weaker PR-AUC scores are due primarily to loss of precision at higher call probabilities. The tendency towards stronger performance on RNA-FISH images and very poor performance on images derived from non-FISH techniques is not unexpected given that it was a smFISH model that was used for prediction across all experimental groups. What was perhaps more unexpected was the poor PR-AUC and recall seen for HeLa CY5 and *C. elegans*, as the signal in these images are primarily clean puncta. The reduced performance might be the consequence of the wide signal amplitude range present in these groups.

### F-Scores at auto selected thresholds reveal strengths and weaknesses of tools’ algorithms

Similar to with our simulated dataset, we used F-Score (Equation 1C) as a metric for measuring automated threshold selection performance in TrueSpot and Big-FISH, the only tools of the four that offer automated threshold selection. The results are shown in Figure 3D. TrueSpot scored significantly higher in two of the seven RNA batch groups (statistical calculations in Table S2C) and performed slightly better on the Tsix-AF594 test image. The difference in performance between tools in the other five groups was not significant, though some general trends in relative consistency and performance can be observed.

TrueSpot scored considerably better than Big-FISH for both protein GFP fusion pairs, but there was no significant difference in the HeLa GFP group (n = 6, p = 0.39), with both tools tending to pick thresholds yielding call sets in poor agreement with the manually chosen set. TrueSpot also tended to score better for both histone group marks, though the difference in the H3K36me3 group was nonsignificant and the H3K4me2 group only contained two member images. TrueSpot also outperformed Big-FISH on the C. elegans singleton image.

Mirroring its overall detection performance, Big-FISH’s auto-thresholder tended to perform better on images in the RNA-FISH groups and worse on GFP and other non-FISH images. As F-Score range is limited by detection performance, this indicates that there was not a large discrepancy between threshold selection and general detection performance in these groups. The histone mark groups were the exception, where the F-Scores trended noticeably lower than the PR-AUC values, pointing to the auto-thresholder as the primary source of disagreement with the manual set. Despite this consistency in image type preference between the general detection and threshold selection, it is worth noting that Big-FISH’s F-Scores trend lower and broader ranged than its PR-AUC scores for most groups. This highlights a degree of inconsistency in threshold selection that sometimes leads to poor quality final call sets from images where Big-FISH’s detection algorithm otherwise performs well. No pattern was immediately clear from manual assessment of these cases that would easily pinpoint what properties of the image or spot count curve were derailing Big-FISH’s auto-thresholder (Figure S10).

In contrast, these data allowed us to clearly characterize and address a specific weakness in TrueSpot’s threshold selection process. Initially, the groups where TrueSpot produced considerably lower F-Scores than PR-AUC scores were sets of images that had large dynamic ranges or variability in signal intensity. Such images produce spot count curves with a gradual bend that in extreme cases can appear diagonal in a logarithmic projection (Figures 1H-I, 2L-M). At the default settings, TrueSpot’s threshold selection algorithm is very good at finding an approximate middle point of the curve bend (Figure S11). However, the very start or very end of the elbow may be more in line with what a human would choose for a given image, though this depends upon the image. Thus, it is difficult to formulate a one-size-fits-all approach for these images with high variability in signal intensity.

We developed additional thresholder parameter presets that favored various degrees of either precision or recall, functioning by adjusting the final call toward one or the other end of the detected range (Table S3). When we applied thresholder presets that better reflected the intensity profiles of these wide-range or signal-variable image batches, the agreement with manual reference sets increased noticeably (Figure S12).

### Tool agreement on per-cell spot counts is highly variable between experimental image groups

Practical applications of punctate signal quantification software often include the derivation of target molecule counts in single cells. For all images with sufficient imaging data to conduct cell segmentation, per-cell spot counts were derived using the thresholds automatically derived by TrueSpot and Big-FISH. Probability distributions and TrueSpot versus Big-FISH plots of per-cell counts for eight experimental groups are shown in Figure 4A-B.

**Figure 4.**
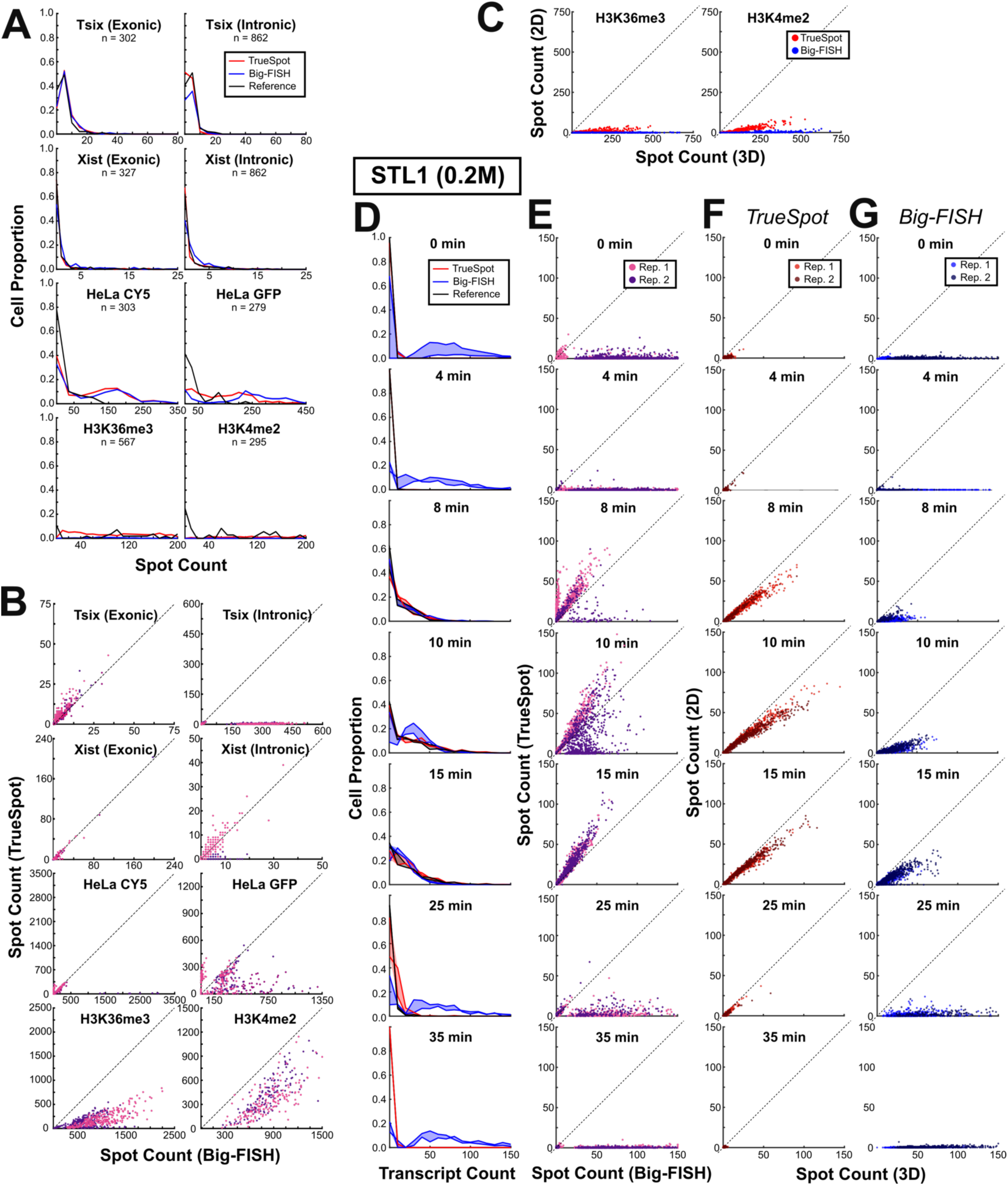
Comparing single cell spot counts across conditions and image groups provides insight into how image characteristics affect tool behavior. Per-cell transcript count probability distribution plots, TrueSpot versus Big-FISH count scatterplots, and 2D maximum intensity projection versus 3D stack count scatterplots for eight experimental images groups (A-C) and subsampling of STL1 measurement time course after 0.2M NaCl stimulation (D-G).(A) Solid lines represent single cell transcript count distributions for TrueSpot with auto-thresholding (red), Big-FISH with auto-thresholding (blue), and reference derived from manual thresholding (black). Supergroup total cell counts are displayed under group names. (Bin sizes: Tsix I/E: 5; Xist I/E: 1; HeLa-CY5: 35; HeLa-GFP: 25; H3K36me3,H3K4me2: 10) (B) Single cell spot counts made by TrueSpot versus Big-FISH for each supergroup. Each point represents a single cell, point fill color represents subgroup (ie. groups are further split by experiment, time point, etc.. Split method varies by group. See Table S4). *y = x* line shown for each scatterplot. (C) Single cell spot counts produced from full stack derived maximum intensity projections versus full stack in 3D for both histone groupswithTrueSpot (red) and Big-FISH (blue).(D) Per-cell probability distributions (bin size = 10) for STL1 (0.2M) time course. Solid lines represent each of two biological replicates, fill represents space between replicates. (E) Single cell spot counts made by TrueSpot versus Big-FISH for each STL1 (0.2M) time point. Each point represents a single cell, point fill represents biological replicate. *y = x* line shown for each scatterplot. (F-G) Single cell spot counts produced from full stack derived maximum intensity projections versus full stack in 3D for a subset of STL1 (0.2) time points with TrueSpot (F) and Big-FISH (G).

High agreement between tools was seen in both Xist groups, with both tools identifying most cells as having no to single-digit spot counts in line with the reference subset (Figure 4A), and linear regression fit slopes tending toward 1 during assessment of tool count correlation (Figure 4B, Table S4). The three highest Pearson estimates for correlation of all 22 subgroups came from exonicXist (Table S4). Poorer correlation in the remaining subgroups appeared to be due to slight relative overcalling on the part of Big-FISH or undercalling on the part of TrueSpot, particularly in one of the intronic Xist groups. Though the low number of positive spots perhaps makes correlation appear stronger than it would was there more signal in these images, it is encouraging that both tools were able to correctly identify threshold values that would yield appropriate spot counts in these cases of low expression.

In contrast, while there is high agreement between the two tools and reference subset for the exonicTsix group, count correlation is stunningly poor for the intronic group (Figure 4A-B). Though it would seem from the probability distribution in Figure 4A that there should not be a large discrepancy in counts between TrueSpot and Big-FISH, Figure 4B shows that when overcalling by Big-FISH does occur, it is to an extreme magnitude. Furthermore, such overcalling is not infrequent enough for these cases to be considered outliers (Table S4).

An interesting feature of the HeLa groups is that the two tools tend to agree with each other more than either with the reference, particularly for the CY5 channel (Figure 4A). It may be that the subset of images that reference sets were created for was not representative of the group as a whole. A likelier explanation is that the manual curator was more stringent in spot selection than either tool, resulting in both tools either outperforming or underperforming relative to the manual curator. There was a pattern of relative overcalling with TrueSpot tending toward a higher frequency and Big-FISH tending toward a higher magnitude present in these groups, though disagreement seemed to be more the exception than the rule (Figure 4B). In the HeLa GFP group, cases where TrueSpot relatively overcalled were usually cases where Big-FISH detected very few to no spots in a given cell, whereas the same did not hold true the other way around. As these GFP images are among the noisier and blurrier of the groups, this could be the result of a greater sensitivity to noise pushing Big-FISH to select a higher threshold.

TrueSpot and Big-FISH exhibit fairly high raw correlation (Pearson for all but one subgroup range from about 0.7 to 0.83, see Table S4) between per-cell spot counts in the two histone mark groups, but very poor agreement. Linear regression of the relationship between the tools’ counts (Figure 4B) produced lines with slopes of around 0.15 to 0.3 for H3K36me3 and 0.6 to 0.8 for H3K4me2 (Table S4). The three subgroups with high Pearson estimates also had surprisingly low projected y-intercepts of around −100 to −250.In other words, Big-FISH tended to call about 100 plus 3 to 7 times as many spots per cell as TrueSpot in the former group and about 150 to 250 plus 1.7 to 1.25 times as many in the latter. With such consistent behavior, it is possible that despite the inherent difficulty of thresholding images with such high signal range (Figure 2K-O), either tool could become capable of attaining excellent accuracy with these groups with only mild sensitivity tuning.

### Maximum intensity projections produce lower per-cell spot counts than their counterpart 3D stacks

Though many modern imaging setups are capable of producing 3D image stacks as output, it is common for labs to condense these stacks into 2D maximum intensity projections (MIPs) to simplify analysis and save storage space. Additionally, due to difficulties introduced when accounting for a third, usually anisotropic dimension in image processing, many image processing tools including deepBlink^18^ and other current deep learning approaches^19–21^ will only work with 2D inputs.

MIP derivation is an inherently lossy process, though to what degree this data loss affects the quantitative results and downstream biological interpretation is generally unclear and likely context dependent. Cases of particular concern might be when the target molecule is abundant and densely clustered, thus brighter signals would obscure other signals above and below leading to potentially drastic undercounting.

We compared the per-cell spot counts obtained from running MIPs derived from our set of histone mark IF images, which have very densely clustered signal puncta, with the full stack per-cell spot counts for both TrueSpot and Big-FISH (Figure 4C, Table S5). For this analysis, the threshold used across all cells in both 2D and 3D was set to a fixed value derived from the mean of thresholds at peak F-Score for each channel (see Table S6 for threshold values used). The MIPs were derived from full stacks without trimming any z-slices.

For both marks and both tools, the counts obtained from the 2D MIPs were consistently a small fraction of the counts obtained from the full 3D stacks (Linear regression slope: H3K36me3 TrueSpot: 0.050, Big-FISH: 8.68 x 10^−4^; H3K4me2 TrueSpot: 0.159, Big-FISH: 0.014). This pattern did not change when the stacks were trimmed to only the most in-focus slices, though the degree of the effect was marginally smaller (Linear regression slope: H3K36me3 TrueSpot: 0.191, Big-FISH: 0.017; H3K4me2 TrueSpot: 0.213, Big-FISH: 0.051, Figure S13, Table S5).

### Tool agreement on per-cell spot counts is poor for STL1 at low expression time points, but not CTT1

Per-cell spot count distributions were also derived for each time point across the *S. cerevisiae* osmotic stress time course image dataset^24^. Distributions for certain time points of STL1 at 0.2M are shown in Figure 4D-E (full results for STL1 0.2M and results for the other three experiments can be found in Figures S14-S17).

Both tools appear to show high agreement with each other and the manually thresholded set for CTT1 at 0.2M, and for most time points at 0.4M, though Big-FISH trends a little lower than TrueSpot at expected peak expression (Figures S14, S15). The discrepancy between counts made by TrueSpot and Big-FISH is much more striking for STL1 as can be clearly seen in Figure 4D-E, where Big-FISH grossly overcalls relative to TrueSpot at most time points. Interestingly, the handful of time points where Big-FISH’s spot counts appear closer to TrueSpot’s and more in the range of what is expected within single cells are the time points where expression is expected to be higher (typically peaking at around 10 to 15 minutes^28–30^). Thus, Big-FISH’s overcalling behavior appears to occur when STL1 expression is low, that is to say, whenthere is little signal. This phenomenon is also seen in all three biological replicates of the 0.4M experiment as well (Figures S16, S17), so it is not simply the result of a batch specific effect.

Maximum intensity projection (MIP) derived per-cell counts trended lower than 3D counts from the same cell, though to a less dramatic degree in the salt-shocked yeast cells than with the mESChistone marks (Figure 4F-G, full time courses in Figures S18, S19). Linear regression slopes fell between 0 and 1 for almost every comparison made across the time courses for both tools, with the only exceptions being at very low expression time points (Table S5). Furthermore, traces of a potentially non-linear rather than linear relationship are visible at the highest expression time points (Figure 4F) perhaps indicating that the discrepancy in spot count between 2D projections and 3D stacks increases as the number of spots increases. This would make sense, as it would be expected that as the amount of signal increases, the amount of signal that would be obscured by collapsing one dimension would also increase.

### Partially manually derived RNA counts could be replicated with automatically selected thresholds

The *S. cerevisiae* osmotic stress time course dataset (n = 337 image stacks, 0.2M: 2 biological replicates, 0.4M: 3 biological replicates) came from a previous set of experiments that our lab had published^24^. We compared RNA counts derived from the automatically thresholded call sets of TrueSpot and Big-FISH to the counts used for the published paper (Figures 5, S20, S21). The published RNA counts were derived using an earlier version of the TrueSpot spot detection module with manually selected thresholds being chosen for each of the two channels and applied to all images.

**Figure 5.**
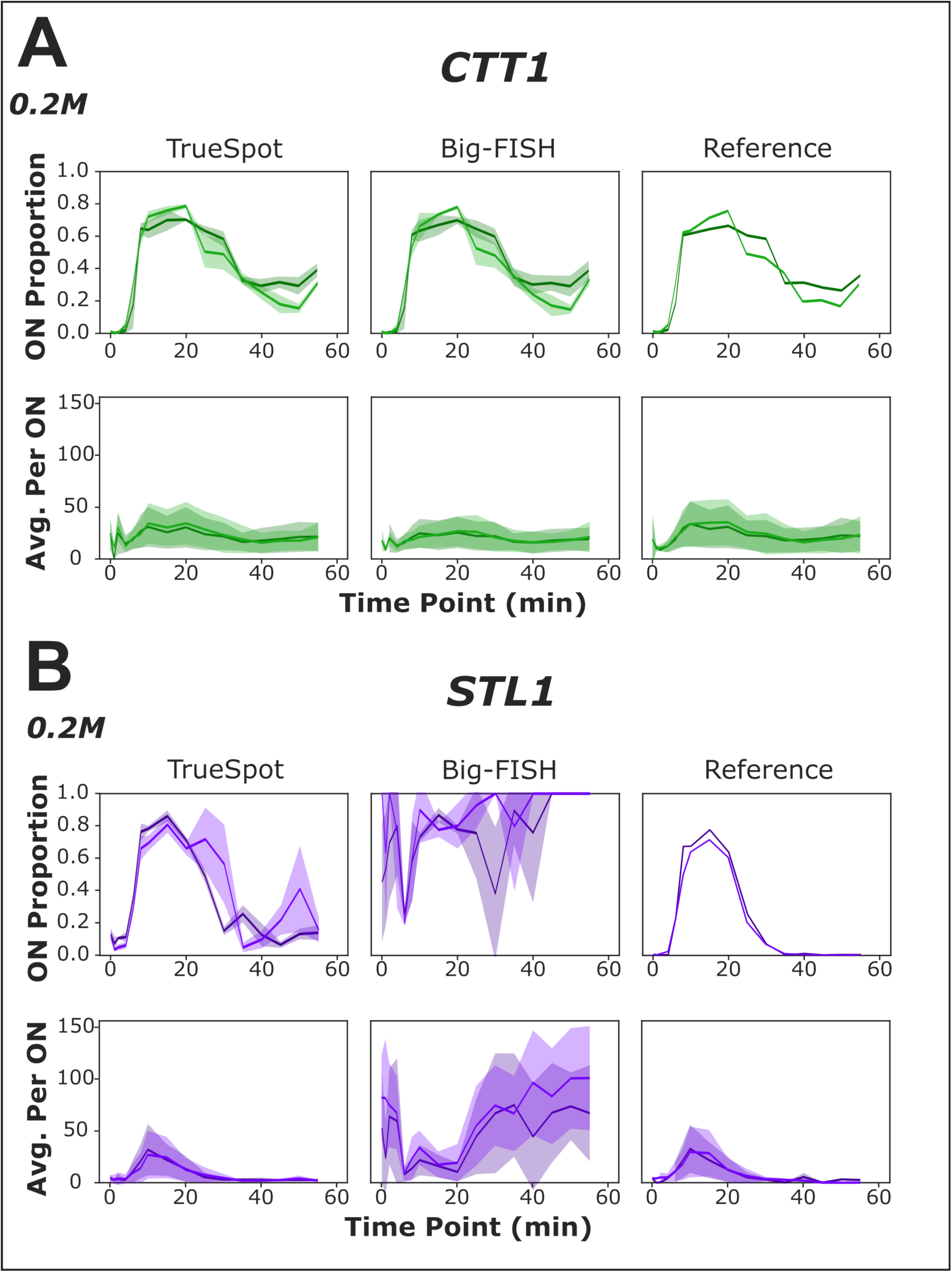
RNA quantification measurements derived from manual thresholding can be replicated using automated thresholding. Measurements of ON proportion of total cells (top) and average transcripts per ON cell (bottom) derived from analysis of experimental time course images tracking CTT1-CY5 (A) and STL1-TMR (B). Comparison of TrueSpot (left) and Big-FISH (middle) fully automated spot detection pipelines (using the same cell segmentation mask from TrueSpot) with measurements from previously published data using manual thresholding with a prototype of TrueSpot (“Reference”, right). Each line represents a biological replicate (0.2M n = 2), mean value across technical replicates (ON proportion) or all ON cells (Average per ON). Shaded areas are mean ± standard deviation at each time point. For 0.4M replicates, see Figures S20 and S21.

TrueSpot was able to replicate both the proportion of cells that were transcriptionally “on” (≥ 8 transcripts/spots for CTT1, ≥ 2 for STL1) and the approximate average number of transcripts per “on” cell across the assayed time points for CTT1 at both input concentrations (Figures 5A and S20, paired t-test outputs in Table S7– magnitude of estimate values reflect general distance between compared lines, signs reflect direction relative to reference). Big-FISH also replicated the CTT1/0.2M “on” proportion curve from the published dataset (Table S7). However, at 0.4M it produced extremely noisy results for two of the three biological replicates at 10-20 min (Figure S20B).

Both tools were able to replicate the average transcripts per “on” cell curves for CTT1 at both concentrations (Figures 5A, S20, paired t-test outputs in Table S7), though Big-FISH’s RNA counts trended slightly lower (consistent with previous observations regarding the data shown in Figures S14 and S15).

TrueSpot was able to decently replicate the manually thresholded results for STL1 at both concentrations (Figures 5B, S21, statistical calculations in Table S7), albeit not as cleanly as with CTT1. In stark contrast, Big-FISH had immense difficulty with this channel, producing such great fluctuations in threshold values and spot counts between replicates that the resulting curves are very noisy and in the case of per cell average counts, explicitly contradictory to the reference. As was noted in regards to Figure 4C-D, gross overestimations of spots per “on” cell by Big-FISH tend to occur primarily at time points where low expression is expected.

Manual evaluation of cases of extreme threshold selection (Figure S10) and the PR-AUC versus F-Score discrepancies in the yeast time course images with manual reference sets (Figure 3C-D) indicate that the inconsistency in performance from Big-FISH is likely due to auto thresholding, not initial detection of local intensity maxima.

While threshold selection inconsistency by itself does not necessarily imply inaccuracy, the extreme variation in spot counts made between technical replicate images taken using the same microscopy setup at the same time, the wide variability in threshold values, and the fact that this variability is not observed with manual inspection and thresholding, point to a limitation in Big-FISH’s automatic thresholding algorithm rather than a genuine biological phenomenon or heterogeneity in the source photographs.

We also assessed the impact that setting a fixed threshold for an image batch would have on RNA counts versus using the automatically determined per-image threshold values. The results are discussed in more detail in Supplemental Results. Briefly, we found that fixing a batch threshold to the mean of the automatically selected thresholds for all member images reduces outliers and thus noise in RNA counts (Figures S22-S25, Tables S6,S8,S9). However, this effect is mild, particularly on experimental data and does not necessarily increase accuracy (Figures S24-S25, Table S9).

Using maximum intensity projections (MIPs) instead of full stacks to derive per-cell counts and “on” proportions (Figure 4F-G) produces “on” proportion time course plots the same or similar to those produced by 3D stack counts when using a fixed threshold (Figures S26-S27). This is not unexpected, since a cell need only pass a low count threshold (8 for CTT1 and 2 for STL1) to be considered “on”. However, plots of average spots per “on” cell show frequent plateauing at high expression time points and trend overall lower for both tools when using MIPs as compared to stacks. This trend not only persists when automatic thresholding is used instead of fixed threshold values, but is often amplified. Additionally, use of automatic thresholding on MIPs introduces extra noise, likely as a result of the thresholding algorithms having more difficulty identifying a signal-noise boundary on the flatter spot count curves produced from 2D images(Figures S26-S27, Table S5).Overall, it is apparent that a notable loss in data resolution was incurred by collapsing these image stacks to their MIPs.

### TrueSpot tends to show more consistent threshold selection behavior than Big-FISH

Reasoning that image stacks from the same source batch and channel should have similar intensity profiles, we compared the variability of threshold selections by TrueSpot and Big-FISH between similar images (Figures 6, S28, S29).

**Figure 6.**
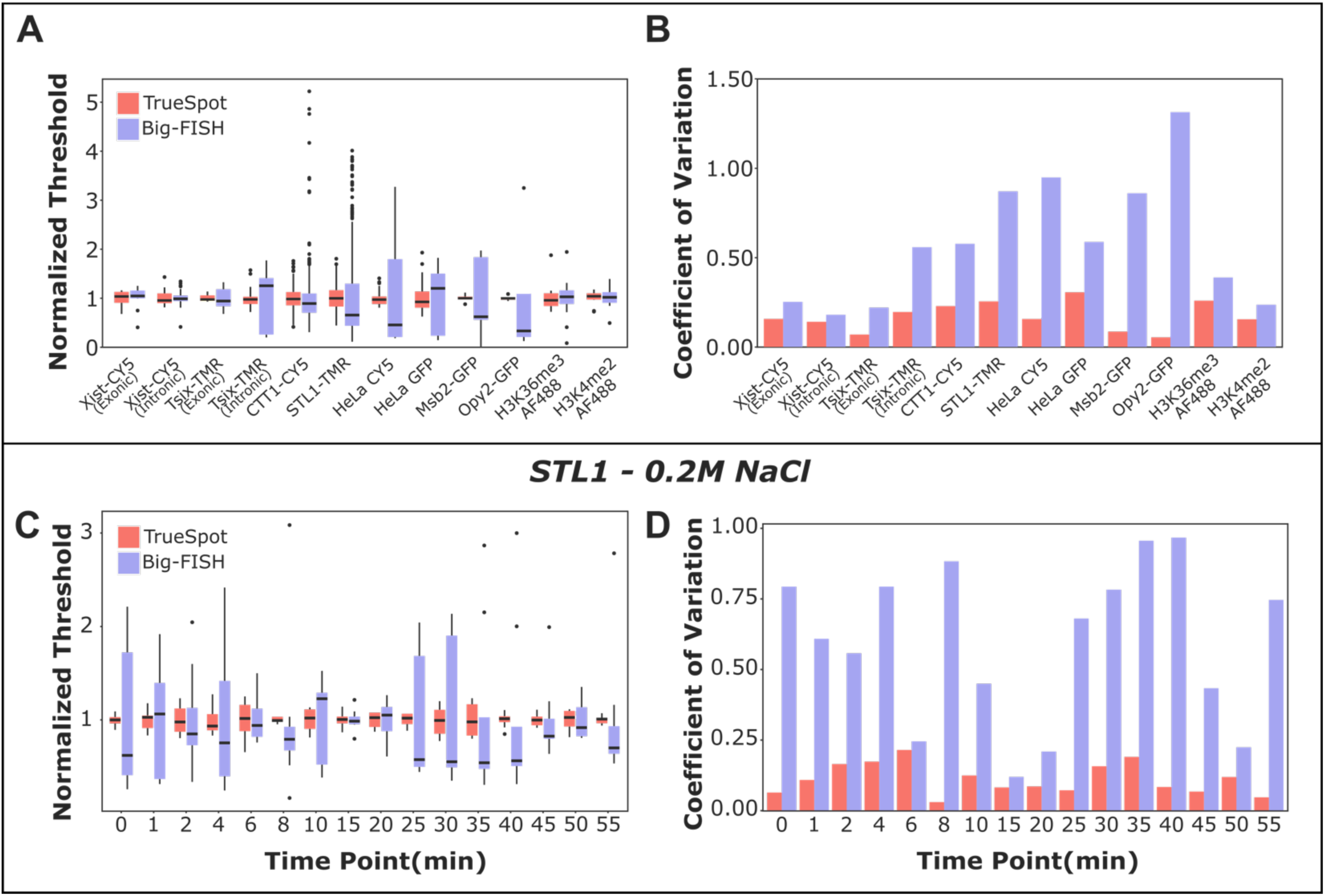
TrueSpot is consistent in threshold selection within image batches. Variation in threshold selection between images in 12 experimental groups by tools with automated threshold selection (TrueSpot (red), Big-FISH (blue)). (A) Distribution of selected thresholds normalized to batch average (box bounds and midline represent 25^th^, 50^th^, and 75^th^ percentiles, whiskers extend to further values within 1.5 * IQR from box edges, dots represent outliers). (B) Coefficient of variation (standard deviation / mean) for each batch. (C-D) Variation in threshold selection across osmotic stress time course images (STL1-TMR channel, 0.2M peak stimulation), grouped by time point. Analyses of additional time courses can be seen in Figures S28 and S29.

The distribution of threshold values normalized to each group mean (Figure 6A) and the coefficient of variation (ratio of the standard deviation to the mean) (Figure 6B) were plotted for each major group of experimental images. (Image counts and exact values can be found in Table S10). The results from TrueSpot tended to display considerably lower coefficients of variation and median threshold selections in closer agreement to group means than the results from Big-FISH. Overall, this shows that TrueSpot had more consistent threshold selection between similar images.

This trend was for the most part retained when the CTT1-CY5 and STL1-TMR groups from the *S. cerevisiae* time courses were broken down by experiment and time point (Figures 6C-D, S28, S29). The contrast in threshold selection consistency between the two tools was most prominent for STL1-TMR in the 0.2M experiment (Figures 6C-D) and, consistent with the previous discussions on Big-FISH’s overcalling behavior on this image group (Figures 4C-D, 5B), the gap in coefficient of variation between the two tools tended to be greater at time points where lower RNA expression was expected (Figure 6D).

While threshold selection consistency is an informative metric to assess tool behavior in tandem with other analyses, it cannot serve as a direct proxy measurement of overall performance alone as it rests on the aforementioned assumption that images assigned to the same group have roughly the same signal and noise intensity value ranges post-filtering. Though we have found that this is often a reasonable assumption to make for images in the same batch (for example, each biological replicate for each *S. cerevisiae* experiment [^24^] was done on the same day), or even using the same microscopy setup and parameters on different days, it is not a reliable one nor is it necessarily globally applicable due to the possibility of genuine outliers or the processing pipelines of particular tools exaggerating small differences between similar images, leading to very different, but still appropriate threshold selections.

## Discussion

We have developed a software tool for fluorescent signal localization and quantification that is automated and widely applicable. We believe that it addresses the limitations of existing tools, namely the prevalence of unavoidable manual steps and the risk of performance loss the more an input deviated from an ideal mid-high expression RNA-FISH image. TrueSpot can be run on batches all the way from raw image files to per-cell signal quantification statistics without any mandatory manual steps, though fine tuning is available. Our threshold selection algorithm is flexible and its strategy of searching for a range of potential values allows for easier troubleshooting and simplified front-end tuning. As we show, TrueSpot works well both on simulated images and a variety of experimental images, and it is quite consistent in its detection and threshold selection behavior. We also showed that our automatic threshold selection algorithm could replicate the results of a time course experiment that had previously been analyzed using manually chosen thresholds, indicating that in these cases our automated threshold selection performed on par with a human curator.

Furthermore, we highlighted the importance of using a variety of experimentally produced image data in tandem with simulated data for benchmarking detection tool performance. The strength of simulated images lies in the ability to have an objective ground truth set and seed parameters. However, even when seeding for high noise or variability and applying additional filters after initial simulated image generation, we found it difficult to truly reproduce the real-world imperfection we observed in our experimental images. Additionally, the experimental images we used came from different experiments and even different labs, and we tried to ensure that our pool had some diversity in probe, target, and cell types. We included images such as those of GFP fused membrane proteins that would not normally be processed by tools designed for RNA-FISH, and found that our tool and some of the other tools tested worked surprisingly well, while at the same time tools would not perform as well as expected on seemingly classic RNA-FISH images with clear, even puncta.

The major downside to using experimental images for benchmarking is the lack of objective ground truth sets. Instead, tool calls must be compared against reference sets that were generated by hand. Because image stacks that contain no instances of dense signal clustering, blurring, or dim puncta that teeter on the border between signal and noise are quite rare, this manual reference creation strategy inevitably brings with it an element of subjectivity. Thus it is important to consider when interpreting such performance metrics that it is the tool’s agreement with human assessors that is being measured, not its objective detection ability.

There are additional caveats to consider when evaluating software performance in this matter. Usage hindrances and circumstantial errors in wrapper and analysis scripts can make it difficult to assess a tool’s true maximum potential, and the developer’s product will generally be at an unintended advantage due simply to a deeper familiarity with their own program. However, usability and flexibility are still important. A tool that is difficult to use properly or sensitive to deviations from the use cases it was tested on can produce low quality results that are time consuming to fix or not even caught at all.

With this framing, let us revisit the above results from the RS-FISH and deepBlink tools. We chose to include these tools in our assessment because they use different strategies than the classic Laplacian of Gaussian filter that we use and we were interested in their potential.

Despite technically possessing a command line interface and batch processing capabilities, RS-FISH seems to be primarily intended for manual use via the Fiji GUI. Indeed, we found RS-FISH to be so sensitive to small changes in input parameters that it is imprudent to *not* use the GUI to finely tune these parameters for each and every batch. Even then, if a batch is sufficiently heterogeneous the quality of the output may be quite mixed. Thus, it is very likely that despite initial manual tuning via the Fiji plugin, image-specific ideal parameters could not be found and applied to many of the benchmarked images. Our takeaway is not that RS-FISH is a subpar tool, rather, that because it requires extensive manual work to operate at its full potential, and our objective is to reduce or eliminate manual input, it may not be the most practical tool to address the specific issues we have discussed.

deepBlink utilizes a deep learning approach instead of the standard filter-based image processing methods. This allows it to bypass any threshold step entirely allowing the prediction module to operate entirely automatically. Unfortunately, as is an inherent problem with machine learning approaches, the performance of the prediction step is highly dependent upon the quality and composition of the training set used to create the model that the predictor uses. Pre-trained models provided by the developer or community can be incredibly useful, but there are many subtle factors that can make datasets unique enough to throw off such models. The above data are an example of mixed results on different image sets yielded by a developer provided general model. It may thus be necessary to train a new model using a subset of the data of interest. This, however, is not always a trivial task. Training requires an “answer key” for each input item, and the best way to get such an answer key for an experimental image is to manually choose what to consider a spot. Essentially, the need to train a new model loops back into another potentially labor intensive manual step that reintroduces human subjectivity.

One other important issue to note about deepBlink specifically is that as of writing, it only processes images in 2D. The prediction module accepts image stacks of 3 or more dimensions, but these are processed slice-by-slice, producing an output with many redundant spot calls that need additional user-side post-processing to merge -an additional step that can introduce opportunity for error. The training module does not accept images with more than 2 dimensions at all. This forces the user to make additional compromises to create a model from a set of 3D image stacks. Taking model sensitivity and coordinate conversion into consideration, deepBlink’s relatively poor performances on our benchmark tests should not rule it out as a useful tool. As with RS-FISH, our criticism is not of its capabilities, but rather, the difficulties involved in consistently bringing out its best with many kinds of datasets, and the risk of inaccurate results going undetected should it fall short.

However, this apparent inability to consider more than two dimensions at once is a major criticism we have of deep learning approaches in general at this point in time. Slice-by-slice processing requires extra work on the user side to merge the results into a 3D call set. Use of maximum intensity projections seems to be more common, but these risk a loss in resolution in turn yielding depressed signal measurements (Figure 4C,F,G). Furthermore, the magnitude of signal obfuscation increases as the amount of signal increases. This introduces an additional signal clipping threshold above which higher signal levels are indistinguishable from each other. These factors have the potential to lead to output data that are so noisy or distorted that biological interpretation may be significantly impacted.

TrueSpot and Big-FISH were designed specifically to process both 2D and 3D images and to circumvent the kinds of issues with unavoidable manual steps discussed above by automating as much as possible, including threshold selection. Additionally, we designed TrueSpot to cover the handful of problems we had with Big-FISH. Though Big-FISH’s automatic threshold selection process appeared to perform on low expression RNA-FISH images about as well as it did on high expression images in our tests, variable intensity caused a noticeable decrease in performance. Big-FISH seemed to struggle when provided images with blurrier signal as seen with both GFP groups. Our main concern however, was that the automated threshold selector would sometimes pick extreme, inappropriate threshold values and manual examination of these cases did not make it clear why it behaved this way on some images, but not others in the same batch. As starkly demonstrated in Figure 5, relying solely on the auto-thresholder could produce usable results as in the case of CTT1-CY5, but could also yield extremely messy, nonsensical results as in the case of STL1-TMR. At time of writing, neither Big-FISH’s auto-thresholding wrapper function nor its component functions appear to accept tuning arguments that can directly influence the threshold selection given a constant spot count curve.

The primary remaining concern we have with both tools is their persistent difficulty with thresholding images with high variability in signal intensity. TrueSpot is at the very least consistent, robustly identifying the approximate center of the first elbow feature at default settings. However, it may actually be the start of the elbow or the end of the elbow that make for preferable threshold selections for many images. TrueSpot has parameter presets that favor precision or recall, but the choice of which to implement for a given image batch still lies with the user. Thus, while we believe we have succeeded in reducing the need for subjective human intervention in the quantification process, we have not quite eliminated it entirely.

We think it is important to highlight that threshold selection for such images with high signal variability might be a general issue. It is theoretically possible to design an algorithm that can detect any specific part of an ‘L’ bend, but *whether* it’s the start, the end, or the middle that best represents the signal-noise boundary may come down to user preference. As noted prior regarding the problems with manual reference sets, it can be difficult for even humans to agree on whether a particular point in this signal/noise overlap region is signal or noise. Thus, though we aim to reduce subjectivity as much as possible, some small amount of human input may be inevitable.

## Materials & Methods

### Image Acquisition

Sample images from the Neuert lab were acquired using a Nikon Eclipse Ti microscope and Hamamatsu Orca Flash C11440-22CU 4v2 CMOS camera. The microscopy setup also included a Nikon Perfect Focus System (TI-PFS-CON 596216), Excelitas fluorescent light source (X-cite Series 120 Q), and 100x VC DIC lens (Nikon MRD01901). Pixel size in the x and y dimension was 65nm. The Micromanager software application was used to control the microscope and save the image data.

mESCs were imaged at 100X in stacks of 69 or 81 z-slices spaced 300nm apart. All images included a DAPI channel (Semrock Brightline DAPI-5060C-NTE-ZERO; 20ms exposure; channel 1, all images) for nucleus identification and a light transmission channel “TRANS” (10ms exposure; channel 4 in Xist/Tsix combined images, channel 5 in Xist/Tsix/histone mark and Xist or Tsix only images). The lncRNA Xist was probed using CY5 (Semrock Brightline NIK-0013 RevA-NTE-ZERO; 1000ms exposure; channel 2). The lncRNA Tsix was probed using either TMR (Semrock Brightline SpGold-B-NTE-ZERO; 1000ms exposure; channel 3) or AF594 (Semrock Brightline NIK-0012 RevA-NTE-ZERO; 1000ms exposure; channel 3). Histone marks H3K36me3 and H3K4me2 were immunofluorescently probed using AF488 as a fluorophore, but were imaged using a yellow fluorescent protein (YFP) filter (Semrock Brightline YFP-2427B-NTE; 500ms exposure; channel 4).

The STL1 and CTT1 timecourse images in *S. cerevisiae* were acquired at 100X in z-stacks of 25 images 200nm apart. As with the mESC images, all images of this set include DAPI (20ms exposure; channel 3) and TRANS (20ms exposure; channel 4) channels. CTT1 was probed with CY5 (1000ms exposure; channel 1), and STL1 was probed with TMR (1000ms exposure, channel 2). Specifications can also be found in^24^.

Additional technical specifications for the Msb2-GFP and Opy2-GFP images are the same as those specified in Jashnsaz, et al. 2021^31^. Images were acquired at 100X in stacks of 13 images 500nm apart. Images of this set have two channels - GFP (Semrock Brightline FITC-2024B-NTE-ZERO; 150ms exposure, channel 1) and TRANS (10ms exposure, channel 2).

Test images of H-128 (HeLa) cells shared by the lab of Brian Munsky at Colorado State were acquired using an Olympus IX81 confocal microscope at 60X (160nm per pixel in x and y) and an EMCCD camera (AndoriXon Ultra 888) using the Slide Book software. Image stacks were comprised of 27 z-slice images 500nm apart and four channels. Channels used in this analysis were those intended to visualize MCP-GFP targeted to a reporter gene containing MS2 loops (488nm; 100ms exposure; channel 2) and a CY5 smiFISH probe for that same reporter RNA (647nm; 300ms exposure; channel 4). Additionally, images included channels for DAPI (405nm; 100ms exposure; channel 1) and a cytosolic marker (561nm; 100ms exposure; channel 3)^26^.

### Image Simulation

The Mueller lab’s Sim-FISH tool v0.2 (https://github.com/fish-quant/sim-fish)^17^ was used to generate 30 simulated image stacks using parameters similar to the properties of the experimental test images. Generation of simulated images applied common values to some Sim-FISH parameters (random_n_spots = True, random_n_clusters = True, random_n_spots_cluster = True, subpixel_factors = None). Variable parameters for each image simulation are outlined in Table S11. The python wrapper script used to generate these simulations is included as “simfish_wrapper.py” in supplementary data.

To simulate the dense clusters of diffuse non-punctate signals as observed in our GFP tagged protein images, simulated stacks were first generated using the parameters specified in the Supplementary Table, then a 3D Gaussian filter (xy radius = 7, z radius = 2, amount = 4) was applied (Equation 3) to blur the simulated image.

#### Equation 3. Gaussian Filter

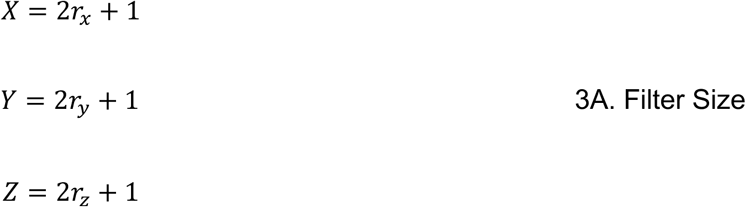

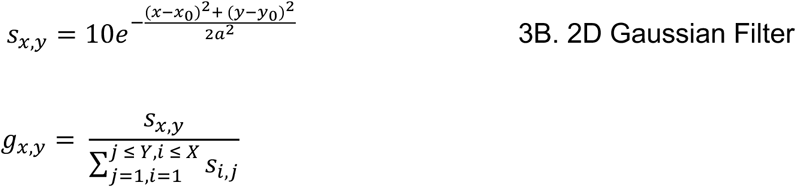

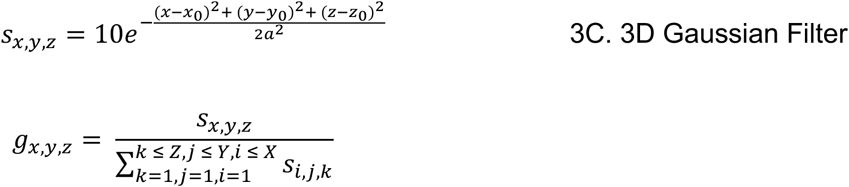

Three sets of 512 x 512 x 16 simulated images varying simulation parameters were also generated using Sim-FISH. The first set was comprised of 5 sets of 7 images generated by varying a single parameter while holding the other 4 constant. The python script used to generate this set of simulated images is also included as “simfish_wrapper_2.py”, and the parameters for each image instance are also listed in Table S11. Varied parameters were total spot count, amplitude variability (random_amplitude), background variability (random_noise), intensity average (amplitude), and cluster density (n_clusters and n_spots_cluster).

The second (simvarmass) and third (simytc) Sim-FISH sets were large sets (1000 and 500 images respectively) generated while varying amplitude variability, background variability, intensity average, background average (simytc only), and spot count randomly and simultaneously (see simfish_wrapper_3.py and simfish_wrapper_4.py). The common parameters and randomizer ranges for these simulations are listed in Table S11. Parameters for individual images are included in a csv table for each set and can also be found in the simulated image mat files.

Two additional filters were applied to all simytc images after initial simulated image generation (see repository script test/test_importsimytc_230330.m). To simulate uneven illumination on a given plane favoring the center of the image, an edge darkening factor between 0.01 and 0.10 was randomly generated for each simulated image stack. This factor was used to generate a plane filter with values proportionate to the selected edge darkening factor and a given pixel position’s distance from the center of the plane. For each plane, a small amount of noise (up to 5%) was introduced to the darkening filter as well before application. The second filter effect was intended to simulate out-of-focus z planes particularly near the extreme top and bottom of the stack. A gaussian blur with a radius of 5 was applied to each z plane, with the strength scaled based upon the distance of the plane from the approximate center of the stack. The maximum strength for each stack was a random value between 1.1 and 2.5. All parameters for additional filter applications as well as the unfiltered simulated stacks are available in the simulated images’ respective mat files.

### External Datasets

The *C. elegans* embryo and simulated images shared by the Preibisch lab for benchmarking in their GitHub repository for RS-FISH^16^ (https://github.com/PreibischLab/RS-FISH) were included in our set of test images.

### Cell Segmentation

Automated cell segmentation was performed on image stacks that included TRANS and DAPI channels for cell boundary and nucleus recognition respectively. This was done using a tool we developed previously^23^ to generate masks demarcating individual cells and their nuclei. The GUI version was used for all test images originating from the Neuert lab rather than the command-line based current release version (which can be run using the celldissect/A0_GUI_seg_outside_generalized.m script). The internal algorithms are unchanged.

Cell segmentation masks for the HeLa groups were provided by the Munsky lab as TIF files and converted into MATLAB binary files for processing.

### Background Masking

Cell segmentation and TRANS channel data were used to generate boolean background masks for applicable test images. 2D projections of 3D image stacks were produced by calculating the standard deviation of intensity values in each xy pixel position over z in the TRANS channel. When the resulting distribution of projection values was bimodal, the local minimum between the peaks was used as a threshold value. When the distribution was not cleanly bimodal, a standard deviation threshold was chosen by enhancing the contrast of the projection and finding the first local minimum in the first derivative approximation after the distribution peak. The first derivative approximation value in at this point in the curve was divided by an arbitrary constant and the threshold value was selected when the first derivative approximation fell below this value. The cell segmentation mask was inverted to pass pixels that were not called inside of a cell by the segmentation software, and this mask was multiplied by the mask of pixels that fell below the previously described threshold in the standard deviation projection to produce an initial background mask. Small holes in the mask were then filled using MATLAB Image Processing Kit methods. This process was automated for all images it was applied to.

### Spot Detection

Spot detection was performed by applying a Laplacian of Gaussian transformation to individual slices in raw image stacks and searching for local intensity maxima in 3D. Before filter application, automated removal of “dead pixels” was performed to reduce false positive calls resulting from pixel positions with no data due to ordinary defects in the imaging hardware. The dead pixel detection algorithm defined dead pixels as non-border xy positions where the intensity value after application of an edge detection filter (Equation 4) was greater than or equal to the mean intensity of its z-slice plus 3 standard deviations for at least half of the z-slices sampled (the greater of 6 or one-quarter of the total z-slices). Dead pixels were subsequently “cleaned” by setting their intensity values in each z-slice to the mean of the values of the eight adjacent pixels in 2D. Raw images where the difference between the maximum and minimum intensities was less than 256 were automatically linearly rescaled to a range of 0 - 511 before further processing.

#### Equation 4. Edge Detection Filter

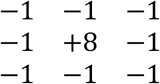

The 2D Gaussian filter was generated according to Equation 3 using standard parameters across all images (xyradius = 7, amount = 2). A 3 x 3 edge detection filter (Equation 4) was then applied to each Gaussian-filtered z-slice. Edges of each slice up to the Gaussian xy radius were trimmed after edge detection filter application by setting the pixel intensity values to zero.

The filtered image stack was then tested against a range of intensity threshold values. This test range was automatically determined for each image. The minimum default was 10 a.u., but for dimmer images where the 80th percentile intensity value in the filtered image fell below 10, this was lowered to 5 or 1 (see RNA_Threshold_Common.suggestScanThreshold). The maximum scan default was set to 500 a.u., but was raised to either the value of the 99th of 99th percentile (top 0.1%) of the filtered image with some leeway or one tenth the maximum intensity value of the filtered image depending upon the ratio of these two values. If the automatically determined scan maximum fell below 500 a.u., the latter default was used.

For each test threshold, voxels with intensity values below the test threshold were filtered out by setting to zero before local maxima identification was performed in 3D. The coordinates of local maxima voxels with intensity values above the mean intensity for the z-slice the voxel was positioned on were returned as the spot callset for the test threshold.

Parameters used for each test image are outlined in Table S12.

### Threshold Selection

Automated threshold selection was performed by computationally searching for abrupt leveling out in curves derived from plotting the number of spots detected at each intensity test threshold versus the threshold value. For these analyses, the curves used for threshold selection were primarily “window score” curves, though the initial count curve and diff curve (absolute value of the first derivative approximation) were used in some cases as well. Window score curves were derived from spot count versus threshold plots by calculating the ratio of variance to mean (Fano factor) over a sliding window across the diff curve. For each test image, a range of window sizes was used to produce multiple window score curves (see Table S3 for thresholding parameters).

Two approaches were applied to each curve to produce a range of possible threshold values. The first approach was comprised of a simple two-piece linear fit. User/preset specified whether the curve itself or a log10 projection of the curve was used for fit. The x-coordinates of the breakpoint and the point where the right-hand fit line first intersects the target curve were saved as threshold candidates. The second approach noted the lowest intensity threshold values where the y value of the curve fell at or below the median y plus the median absolute deviation (MAD) of y multiplied by a range of factors (−1.0 to 1.0 in increments of 0.25). A weighted average of the resulting candidates was taken according to adjustable weighting parameters (Table S3) and rounded to determine the final threshold call.

The RNAThreshold.m script contains all functions related to TrueSpot’s threshold selection process. All preset parameters can be found in Table S3.

### Signal Quantification

The signal quantification step was comprised of both automatically determining how many puncta were in each cell and its nucleus and calculating the total signal intensity by fitting gaussians to called spots and indentifying dense clusters and clouds that could not be easily fitted to a simple gaussian point model.

Gaussian fitting is performed on the voxels surrounding local maxima found by applying the filter and intensity threshold from the spot detection and thresholding steps. Fitting for each maximum was performed on rectangular prism shaped crops around each maximum (xy radius = 4, z radius = 2). Parameters for Equation 3B were fitted to the putative spot in 2D, one z-slice at a time, and the parameters for the spot in 3D were inferred from these. Total signal of a fitted spot was determined by summing up the background subtracted intensity of all voxels within the volume of the fitted ellipsoid (full width at half maximum used for xy radii).

Due to the RNA distribution of STL1 and CTT1 being sufficiently spread out so as to easily resolve individual spots at 100X resolution, we deemed cloud detection unnecessary for the *S. cerevisiae*timecourse dataset. Fixed thresholds derived from the average of all auto-detected thresholds in an image batch (same biological replicate, same channel) were used for the quantification step of the *S. cerevisiae*timecourse dataset (Table S6). Fitted spots were counted for each cell in each image, and the average RNA per cell, number of ON cells (At least 8 transcripts per cell for channel 1, 2 transcripts per cell for channel 2 - see Li &Neuert 2019^24^) and average RNA per ON cell were calculated.

### Tool Benchmarking

Big-FISH 0.6.2^17^ was run using a python wrapper script (bigfish_wrapper.py) in a virtual environment running Python 3.6.3 on the Vanderbilt ACCRE computing cluster. Within the wrapper, the rescaling step was included for all test images. For experimental images, the out-of-focus slice detection function was used to remove the blurriest 20% of slices, and for simulated images, no slices were removed for analysis. Big-FISH performs detection and threshold selection independent of any manually provided scan range, though to dump coordinates for individual test thresholds, a standard scan range of 10 - 1000 was used. This range was adjusted to include automatically determined thresholds that fell outside. Subpixel fitting for the callset at the automatically selected threshold was also run for most test images. Spot coordinates for each test threshold, threshold selection, and subpixel fits were imported into MATLAB structures for further analysis. Coordinates were converted from 0-based to 1-based for ease of use in MATLAB.

RS-FISH 2.3.1^16^ was built from a clone of the GitHub repository, and run headless on the ACCRE cluster via the rs-fish binary. Instances of RS-FISH were run using Java runtime version 13.0.2. All parameters for runs except for the DoG threshold were held constant for each image. The sigma parameter was calculated as the ratio of the expected point radius to the xy voxel dimension, divided by 2.25. The anisotropy parameter was left at 1.0. These were determined prior to batch runs using RS-FISH’s interactive Fiji plugin on Windows on representative test images. DoG threshold was used as the independently varied parameter with a scan range of 4 x 10^−4^ to 0.1 with a step size of 4 x 10^−4^. RS-FISH output tables were imported into MATLAB with 0- based to 1-based coordinate conversion.

deepBlink 0.1.4^18^ was run in a virtual environment using Python 3.8.6 on the ACCRE cluster. For comparison against truthsets and other tool outputs, deepBlinkcallsets for individual slices were merged into single 3D callsets after MATLAB import and conversion to 1-based coordinates. Merging was performed by clustering spots with at least a 50% call probability in adjacent z planes that were within 4 pixels of each other in xy (see RNACoords.mergeSlicedSetTo3D). Call probability was used as a proxy for threshold in curve visualization and tool comparison.

A second model for testing on the simulated data was generated using a small subset of simulated images (n = 100). The image stacks were broken into individual slices and fed into deepBlink’s training module with label sets generated from the ground truth keys. Label sets for each slice included all ground truth spots centered on that slice. All commands used and the list of images used for the training subset are provided in the analysis data repository.

### Maximum Intensity Projection Assessment

Maximum intensity projections (MIPs) were pre-generated from source stacks by taking the maximum intensity at a given x,y position across all z slices. For each source stack, one MIP was generated from the full stack, and one MIP from the range of z-slices that was manually determined to be most in-focus for the batch (z-slices 20-42 inclusive for mESC histone images, 10-16 inclusive for *S. cerevisiae* time course images).

These pre-generated MIPs were provided to TrueSpot and Big-FISH as input using the same xy parameters as their 3D counterparts. For cases where results were compared against a reference set upon import and the MIP had been derived from a trimmed stack, reference set calls outside of 2 z-slices from the trim edges were removed.

Per-cell counts were derived using the same cell masks (see Cell Segmentation above) as were used for the full 3D stacks. Fixed threshold values for the *S. cerevisiae* time course images were the same as the per biological replicate auto-threshold averages used in the time course analyses. Fixed threshold values for the mESC histone images were derived from the mean of the threshold values at maximum F-Score for each target and tool using the images with curated reference sets. When comparing counts from MIPs derived from trimmed stacks with their 3D counterparts, 3D calls outside of 2 z-slices from the trim edges were removed from the 3D counts.

### Truth Set Generation and Performance Quantification

To assess the performance of these software tools on experimental image data, manual “truth sets” were created for a subset of test images. Truth sets were created by manually selecting spots on an image using a both a max z projection and individual slice views as guides. Initial spot selection was agnostic to computer generated callsets, however spot coordinates were snapped to nearest computer calls for evaluation. Each image stack that was assessed was assessed against truth sets generated by at least one individual.

Performance quantification was conducted using measures of recall or sensitivity (Equation 1A), precision (Equation 1B), and F-Score (Equation 1C). False positives were defined as voxel positions called by the automated tool at a given test threshold that were not in the truth set. False negatives were voxel positions that were in the truth set, but were not called automatically at a given test threshold. True positives were voxel positions that were both in the truth set and called by the automated tool at the given test threshold. A call was considered a true positive if its coordinates were within 4 voxel units in xy and 2 z planes of a reference set entry and that entry had not already been matched with a closer call.

Area under the precision-recall curve (PR-AUC) was calculated for each test image/tool pair by calculating the area of the polygon generated by connecting all of the sensitivity/precision mapped points and extrapolating to the x and y axes. Signal-to-noise ratio (SNR) was calculated for simulated images using the mean amplitude over the standard deviation of the background (μ_A_ / σ_B_). The spot density proxy measurement was calculated as the average number of spots per 9×9×5 box (Total Spots / [(Width/9) * (Height/9) * (Depth/5)]).

Statistical tests were performed in R. The commands and scripts used for these as well as the output tables are available in the analysis data repository.

In cases where a reference set only covered and/or a tool was known to analyze part of an image stack, only calls within the intersection of these regions were included in precision and sensitivity calculations. For all simulated images, 7 pixels were trimmed from all edges in x and y regardless of tool. A comparison of the PR-AUC and F-Scores from TrueSpot when the simulated images are trimmed versus untrimmed can be seen in Figure S30.

### Simulated Image Quality Control

To assess the artificiality of simulated images by evaluating “cleanliness”, we used zero voxel proportion (ZVP) as a metric. ZVP was calculated from the LoG filtered image (see “Spot Detection” section above) by dividing the number of voxels with a value of zero by the total number of voxels in the stack. Simulated images with a ZVP at or above 0.7 were filtered out for most simulated image assessments, including the results displayed in Figure 3A-B. See supplementary results for additional analyses.

## Code and Data Availability

All MATLAB code comprising our software pipeline is open source and available on <u>Zenodo</u> (https://zenodo.org/records/13345321).

The parameters used to run Big-FISH on test images are summarized in Table S12. All wrapper and analysis scripts and tables may also be downloaded from the <u>Zenodo</u> repository. mESC RNA and histone test images^25^ as well as the yeast membrane protein images, TIFs of simulated image stacks, call set and analysis files, and other large data files are linked in this repository and are available at (https://www.ebi.ac.uk/biostudies/studies/S-BIAD1316). Yeast RNA test image stacks^24^can be downloaded from the Image Data Resource (IDR) at (https://doi.org/10.17867/10000118).

## Acknowledgements

We would like to thank Luis Aguilera, Linda Forero, and Brian Munsky from the Munsky lab at Colorado Statefor sharing experimental images with us to use for benchmarking.We would also like to thank Eric Gamazon, Bingshan Li, Jason Hughes, John Adams, and our poster and presentation audiences for providing suggestions and feedback on the software, benchmarking tests, and manuscript.Finally, we thank our funding sources: the Vanderbilt Big Biomedical Data Science (BIDS) training program (B.H., 5T32LM012412), Vanderbilt Integrated Training in Engineering and Diabetes (ITED) training program (B.H., 5T32DK101003), Vanderbilt Molecular Biophysics training program (B.K., 5T32GM008320),the NIH Director’s New Innovator Award (G.N., 1DP2GM114849), the National Institute of General Medical Sciences (G.N., 5R01GM115892; 5R01GM140240), the Vanderbilt Advanced Computing Center for Research and Education (ACCRE) (1S10OD023680), and the Vanderbilt Basic Sciences Dean’s Faculty Fellowship.

## Author Contributions

G.N. wrote the initial LoG spot detection and quantification code, which was refined and adapted to mammalian cells by B.K., and later generalized, modularized and optimized by B.H.. The thresholding approach was developed by G.N. and B.H., and implemented and tested by B.H..Experimental manual reference sets were created by B.H. and B.K..Benchmarking tests and analyses were developed by G.N. and B.H., and run by B.H. for all tools and test images. B.H. generated all figures and simulated images, and wrote the base drafts and majority of this manuscript and all public data and code documentation. H.J. conducted the experiments resulting in the GFP-tagged Msb2 and Opy2 images in yeast that were used for benchmarking. B.K. conducted the experiments that produced the Xist, Tsix, and histone mark images in mESCs that were used for benchmarking.

## References

1 Vera, M., Biswas, J., Senecal, A., Singer, R. H. & Park, H. Y. Single-Cell and Single-Molecule Analysis of Gene Expression Regulation. Annual Review of Genetics 50, 267–291 (2016). 10.1146/annurev-genet-120215-034854

2 Hnisz, D., Shrinivas, K., Young, R. A., Chakraborty, A. K. & Sharp, P. A. A Phase Separation Model for Transcriptional Control. Cell 169, 13–23 (2017). 10.1016/j.cell.2017.02.007

3 Adivarahan, S. et al. Spatial Organization of Single mRNPs at Different Stages of the Gene Expression Pathway. Molecular Cell 72, 727–738.e725 (2018). 10.1016/j.molcel.2018.10.010

4 Statello, L., Guo, C. J., Chen, L. L. & Huarte, M. in Nature Reviews Molecular Cell Biology Vol. 22 96–118 (Nature Research, 2020).

5 Escalante, L. E. & Gasch, A. P. The role of stress-activated RNA-protein granules in surviving adversity. RNA 27, rna.078738.078121–rna.078738.078121 (2021). 10.1261/rna.078738.121

6 Femino, A. M., Fay, F. S., Fogarty, K. & Singer, R. H. Visualization of Single RNA Transcripts in Situ. Science 280, 585–590 (1998). 10.1126/science.280.5363.585

7 Raj, A., van den Bogaard, P., Rifkin, S. A., van Oudenaarden, A. & Tyagi, S. Imaging individual mRNA molecules using multiple singly labeled probes. Nature Methods 5, 877–879 (2008). 10.1038/nmeth.1253

8 Chen, K. H., Boettiger, A. N., Moffitt, J. R., Wang, S. & Zhuang, X. RNA imaging. Spatially resolved, highly multiplexed RNA profiling in single cells. Science (New York, N.Y.) 348, aaa6090–aaa6090 (2015). 10.1126/science.aaa6090

9 Eng, C.-H. L. et al. Transcriptome-scale super-resolved imaging in tissues by RNA seqFISH+. Nature 568, 235–239 (2019). 10.1038/s41586-019-1049-y

10 Moffitt, J. R. et al. High-throughput single-cell gene-expression profiling with multiplexed error-robust fluorescence in situ hybridization. Proceedings of the National Academy of Sciences 113, 201612826–201612826 (2016). 10.1073/pnas.1612826113

11 Mueller, F. et al. FISH-quant: automatic counting of transcripts in 3D FISH images. Nature Methods 10, 277–278 (2013). 10.1038/nmeth.2406

12 Tsanov, N. et al. smiFISH and FISH-quant - a flexible single RNA detection approach with super-resolution capability. Nucleic Acids Research, gkw784-gkw784 (2016). 10.1093/nar/gkw784

13 Lionnet, T. et al. A transgenic mouse for in vivo detection of endogenous labeled mRNA. Nature Methods 8, 165–170 (2011). 10.1038/nmeth.1551

14 Perkel, J. M. Starfish enterprise: finding RNA patterns in single cells. Nature 572, 549–551 (2019). 10.1038/d41586-019-02477-9

15 Kowalczyk, G. J. et al. dNEMO: a tool for quantification of mRNA and punctate structures in time-lapse images of single cells. Bioinformatics 37, 677–683 (2021). 10.1093/BIOINFORMATICS/BTAA874

16 Bahry, E. et al. RS-FISH: precise, interactive, fast, and scalable FISH spot detection. Nature Methods 2022 19:1219, 1563–1567 (2022). 10.1038/s41592-022-01669-y

17 Imbert, A. et al. FISH-quant v2: a scalable and modular tool for smFISH image analysis. RNA 28, 786–795 (2022). 10.1261/RNA.079073.121/-/DC1

18 Eichenberger, B. T., Zhan, Y., Rempfler, M., Giorgetti, L. & Chao, A., Jeffrey. deepBlink: threshold-independent detection and localization of diffraction-limited spots. Nucleic Acids Research 49, 7292–7297 (2021). 10.1093/nar/gkab546

19 Niu, Z. et al. Piscis: a novel loss estimator of the F1 score enables accurate spot detection in fluorescence microscopy images via deep learning. bioRxiv (2024). 10.1101/2024.01.31.578123

20 Mantes, A. D. et al. Spotiflow: accurate and efficient spot detection for imaging-based spatial transcriptomics with stereographic flow regression. bioRxiv, 2024.2002.2001.578426 (2024). 10.1101/2024.02.01.578426

21 Laubscher, E. et al. Accurate single-molecule spot detection for image-based spatial transcriptomics with weakly supervised deep learning. Cell Syst 15, 475–482.e476 (2024). 10.1016/j.cels.2024.04.006

22 Lyubimova, A. et al. Single-molecule mRNA detection and counting in mammalian tissue. Nature protocols 8, 1743–1758 (2013). 10.1038/nprot.2013.109

23 Kesler, B., Li, G., Thiemicke, A., Venkat, R. & Neuert, G. Automated cell boundary and 3D nuclear segmentation of cells in suspension. Scientific Reports 9 (2019). 10.1038/s41598-019-46689-5

24 Li, G. & Neuert, G. Multiplex RNA single molecule FISH of inducible mRNAs in single yeast cells. Scientific Data 6 (2019).

25 Kesler, B. K. & Neuert, G. Transcriptional stochasticity reveals multiple mechanisms of long non-coding RNA regulation at the Xist – Tsix locus. manuscript in revision in Nature Communications (2024).

26 Vo, H. D., Forero-Quintero, L. S., Aguilera, L. U. & Munsky, B. Analysis and design of single-cell experiments to harvest fluctuation information while rejecting measurement noise. Frontiers in Cell and Developmental Biology 11 (2023). 10.3389/fcell.2023.1133994

27 Munsky, B., Li, G., Fox, Z. R., Shepherd, D. P. & Neuert, G. Distribution shapes govern the discovery of predictive models for gene regulation. Proceedings of the National Academy of Sciences 115, 7533–7538 (2018). 10.1073/pnas.1804060115

28 de Nadal, E., Ammerer, G. & Posas, F. Controlling gene expression in response to stress. Nature Reviews Genetics 12 (2011). 10.1038/nrg3055

29 Neuert, G. et al. Systematic Identification of Signal-Activated Stochastic Gene Regulation. Science 339, 584–587 (2013). 10.1126/science.1231456

30 Mitchell, A., Wei, P. & Lim, W. A. Oscillatory stress stimulation uncovers an Achilles heel of the yeast MAPK signaling network. Science 350, 1379–1383 (2015). 10.1126/science.aab0892

31 Jashnsaz, H., Fox, Z. R., Munsky, B., Neuert, G. & Fox R, Z. Building predictive signaling models by perturbing yeast cells with time-varying stimulations resulting in distinct signaling responses. STAR Protocols 2, 100660–100660 (2021). 10.1016/J.XPRO.2021.100660

