## Supplemental Text for "TrueSpot: A robust automated tool for quantifying signal puncta in fluorescent imaging"

### Supplementary Results

#### Artificial aspects of simulated images are revealed by idealized result curves

As the details of the spot count curves output by the LoG tools (TrueSpot and Big-FISH) can be very difficult to resolve on a linear scale due to the extreme oversensitivity at the lowest thresholds, all spot count curves were examined using a logarithmic (base 10) projection. The qualitative features of the initial output of the simulated image analyses that stood out to us first were the sharpness of bends and frequent presence of clean plateaus in the spot count curves relative to those produced by experimental images (representative outputs are shown in Figures S1-S3). These features were seen even in the outputs from simulated images with low signal-to-noise ratio (SNR, defined in these analyses as the ratio of mean signal amplitude to standard deviation in background) (Figure S4) and high amplitude variability (Figure S2). It is possible that extreme amplitude variability parameters beyond our dataset's maximum may yet result in curves with more gradual or diagonal features as seen in experimental images with high signal variability (illustrated in Figure 1G-I). Still, it appears evident that though superficially similar, simulated RNA-FISH images tend to retain an unrealistic "cleanliness" that is not typically observed in experimental images, even when seemingly extreme noise addition parameters are provided to the generator. The presence of a clean signal feature in a spot count curve in turn generally produces plots of F-Score versus threshold (Equation 1C) and precision (Equation 1B) versus recall (Equation 1A) with similarly clean attributes such as sharp bends and smooth or straight lines (Figures S1-S2). Thus, the optimal threshold or threshold range for signal detection is often clear and the auto-thresholding algorithms have less difficulty finding it than they do given real experimental images (Figure S1C).

Interestingly, we observed a fourth common curve shape in the simulated dataset that we did not see in the experimental dataset (Figures S2J-L and S3). These curves are characterized by their lack of an initial sharp dropoff, generally starting at or just above the signal plateau before gently dropping off toward zero. As the target of TrueSpot's and Big-FISH's threshold selection algorithms is the resolution of this sharp drop feature, the tools behave unpredictably in such cases where the even the lowest testable threshold value does not produce an extreme overcount. Thus, this curve type is the result of abnormally efficient noise filtration during the detection stage. This is generally not a phenomenon we observe with experimental images and may be an indication that even simulated images generated with high noise level parameters retain an element of artificiality.

In an attempt to filter out the most unrealistic images from the simulated image pool, we calculated the proportion of zero value voxels (ie. the proportion of voxels in a channel stack that have an intensity value of zero to the total number of voxels) in every simulated (Figure S3A-B) and experimental (Figure S3A,C) image in our dataset after application of TrueSpot's LoG filter as a proxy measurement representing cleanliness. Indeed, there was some correlation between SNR and zero voxel proportion (ZVP) (Figure S5D, Spearman = 0.555, p =$6.96 \times 10^{-129}$) in that images with higher SNR tended to have higher ZVP, though there was a small subpopulation of high SNR images that had low ZVP (possibly due to simply having a higher spot count and thus more signal voxels).

Since we had observed that these "no dropoff" images were problematic for the tools' threshold selection, we also plotted ZVP against F-Score for both TrueSpot (Figure S5E) and Big-FISH (Figure S5F) to see whether there was a correlation between ZVP and thresholder performance. Though there did not appear to be a correlation for the vast majority of images, TrueSpot had a small dribble of high ZVP images that TrueSpot was not able to threshold properly likely due to the aforementioned "no dropoff" issue leading to a significant nonparametric anticorrelation overall (Spearman: -0.216, p = $2.86 \times 10^{-18}$). Interestingly, Big-FISH's performance actually increased with ZVP for the subpopulation of images where it performed generally poorly, leading to a slight but significant positive correlation (Spearman: 0.221, $p = 3.21 \times 10^{-19}$).

For quality control purposes, we still decided to filter out the most unrealistic simulated images. The cleanest experimental image produced a filtered result with a ZVP of 0.699 (Figure S5A,C). Thus, simulated images with a ZVP at or above 0.7 were removed from the main analysis pool.

Area under the precision-recall curve (PR-AUC) and F-Score versus SNR distributions for the outputs of all four tested tools on the simulated image dataset using various zero voxel proportion filter values is shown in Figure S31. Importantly, the shapes of the main distributions are unchanged regardless of ZVP cap, and while the ZVP cap is not unreasonably restrictive, only a handful of outliers appear to be subject to the filter.

**Subpixel fitting of simulated images produces mixed results**

We performed subpixel fitting on the simulated dataset using all four tools to compare their ability to pinpoint signal origins with ground truth sets. The results showing the distribution of fit distances from the true coordinates of matched reference set points from all Sim-FISH generated images are shown in Figure S32 (TrueSpot n = 2149054, Big-FISH n = 2903759, RS-FISH n = 2653673, deepBlink n = 1903796).

All tools performed well in xy (Figure S32A, statistics in Table S13A), with at least 75% of all spot fits falling below half a pixel in all four cases, and under a quarter pixel for TrueSpot (median = 0.091, $75^{th}$ percentile = 0.194, mean = 0.162), Big-FISH (median = 0.054, $75^{th}$ percentile = 0.137, mean = 0.147), and RS-FISH (median = 0.105, $75^{th}$ percentile = 0.217, mean = 0.217). There were few fits exceeding 1 pixel in distance from the matched reference point, though it appeared to occur more with deepBlink (median = 0.259, $75^{th}$ percentile = 0.439, mean = 0.402) than any other tool. Big-FISH performed the tightest subpixel fits while TrueSpot and RS-FISH performed only marginally weaker and comparably to each other.

The three 3D tools display interestingly disparate behavior when fitting in the z direction (Figure S32B, statistics in Table S13B). Big-FISH (median = 0.032, mean = 0.081) and RS-FISH perform similarly in z as in xy, but TrueSpot has small, but visible, subpopulations of spots fitted one or more slice away from their reference position.

Though the median z fit distance for TrueSpot is on par with RS-FISH's (TrueSpot median = 0.1, mean = 0.267; RS-FISH median = 0.11, mean = 0.237), around 18.6% (399173 of 2149054) of spot fits fell over half a slice away from their ground truth reference. Still, even TrueSpot's 75th percentile sits at around one-third of a pixel (75$^{th}$ percentile = 0.340), meaning that while its z fitting could use some improvement, it is far from nonfunctional. There are two likely explanations for this subpopulation of spots that are less well fitted in z, that provide the best initial targets for tuning. One is that TrueSpot uses the average of three different approaches for z fitting (attempting fits using xz, yz, and intensity), and one or more of these approaches may need additional troubleshooting or may not be well suited for this simulated dataset. The other is that TrueSpot usually calls spots that are clustered tight enough to overlap as a single signal point. If two spots are overlapping, the z coordinate called for the cluster, and thus used as the starting point for the fit, may be relatively far from the reference z coordinate for either spot.

We also calculated the total xyz distance (in voxels) for all of the 3D fits (Figure S32C, statistics in Table S13C). Unsurprisingly, the distributions were reflective of their xy and z counterparts, with Big-FISH tending to have the tightest overall fits (median = 0.0835, mean = 0.1843). Again, TrueSpot and RS-FISH had similar distributions for the bulk of the fits (TrueSpot median = 0.2206, mean = 0.3639; RS-FISH median = 0.193, mean = 0.3627), but TrueSpot had a subpopulation of spots around a distance of 1.25, most likely due to a poorer z fit. However, for even TrueSpot 88.5% of fits were under 1 voxel away from their "true" value and 74.1% were under half a voxel away, once again indicating that the majority of even the poorer fits were not extreme in their error.

There are a couple of additional points worth noting regarding our benchmarking of subpixel fitting. The first is that we only included the images generated with Sim-FISH in the test set, and Sim-FISH not only shares a development team with Big-FISH, but Big-FISH was tested extensively using Sim-FISH images[17]. There is nothing wrong with this approach, but it is likely that Big-FISH is somewhat calibrated to perform particularly well on these and similar simulated images. The other note of import is that the ground truth set used only integer coordinates and during assessment of fitting, the assumption was made that the exact integer coordinates represented the true center of the gaussian and origin of the signal.  This is probably a fair assumption to make for a simulated image set, but it is very unrealistic. An ideal test would include experimental images in the benchmarking set, but obtaining a ground truth of subpixel signal origins independent of any software tool would be thoroughly impractical.

**SNR of simulated images has little impact on overall detection performance of tools utilizing traditional methods**

We wanted to determine how the noisiness of the simulated images affected tool detection performance, so we plotted PR-AUC against SNR for all four tested tools across the full simulated dataset (Figure S8A, statistics in Table S14B). There was no significant correlation between SNR and PR-AUC for RS-FISH (Spearman p = 0.966), though oddly there was a very mild anticorrelation between SNR and maximum recall (Spearman = -0.379, p = 2.94 x $10^{-55}$) (Figure S6B, statistics in Table S14A). Conversely, TrueSpot and Big-FISH showed no significant correlation between maximum recall and SNR (Spearman: TrueSpot p = 0.654; Big-FISH p = 0.185), but positive correlation between SNR and PR-AUC. In TrueSpot's case, the correlation is quite weak (Spearman = 0.069, p = 0.0063), but in Big-FISH's case, this correlation had a more substantial magnitude (Spearman = 0.341, p = 1.53 x $10^{-44}$). Indeed, there is a visible population of low SNR images in Figure S8A that Big-FISH did not perform well on. This suggests that even with simulated images, Big-FISH's ability to sift out true signal spots from noise spots is more dependent upon SNR than either TrueSpot's or RS-FISH's.

deepBlink's performance was also very sensitive to SNR when the developer smFISH model was used. The positive correlation between SNR and performance is starkly evident for both maximum recall (Spearman = 0.66, p = 4.23 x $10^{-197}$) and PR-AUC (Spearman = 0.651, p = 3.15 x $10^{-190}$). However, this relationship dissipated considerably when the custom model trained on images within the same simulated data set was used (Max Recall: Spearman = 0.108, p = 5.8 x $10^{-5}$; PR-AUC: Spearman = 0.092, p = 6.51 x $10^{-4}$).

Big-FISH showed an even higher correlation between SNR and F-Score (Spearman = 0.58, p = 9.62 x $10^{-144}$; Figure S8B, statistics in Table S14C) than with PR-AUC, as illustrated in Figure S8. This is an indication that not only precision, but automated threshold selection is sensitive to low SNR. In contrast, TrueSpot showed no significant relation between SNR and F-Score (Spearman p = 0.123).

**Measurements of performance tend to drop as spot density increases**

In order to assess whether expression level, or total spot count, had an impact on the performance of any of the benchmarked tools, we first plotted the number of ground truth spots versus spots called at the selected threshold for TrueSpot and Big-FISH to look for general trends (Figure S8C).

Figure S8C reveals not only a propensity for TrueSpot to undercall and Big-FISH to overcall in the minority of cases where a poor threshold selection is made, but an increased prevalence and degree of undercalling by both tools as the number of ground truth spots increases. The latter phenomenon is very likely due to increasing spot density moreso than raw spot count, as tools may either have difficulty distinguishing signals that are too close together or be built to call tightly clustered spots as a single signal, the net intensity and extent of which is to be determined at a later quantification step.

Thus, to quantitatively assess relationships between performance metrics and spot density, we used a proxy measurement that corrected for total image size (spots per 9 x 9 x 5 box) rather than the number of ground truth spots directly (Figure S33). Unsurprisingly, Spearman tests showed anticorrelation between all three metrics for all five tool variations and spot density across the board (Table S15).

RS-FISH and deepBlink Retrained showed the weakest anticorrelations, with PR-AUC for RS-FISH scoring the lowest in magnitude (Spearman = -0.056, p = 0.025). However, it could be that these distributions do not have as significant a fall across increasing spot density simply because their performance did not start as high as the other tools at low spot density.

There is also an interesting trend with the relationship between F-Score and spot density for Big-FISH specifically. Threshold F-Score for the subpopulation of images that performed very poorly actually increases along with spot density, even though the overall trend for the full image set is the opposite (Figure S33C). As seen in Figure S8B, these low F-Score images also tend to have very low SNR. So perhaps this phenomenon is the result of higher spot density leading to a higher proportion of total signal in these noisy images, allowing Big-FISH to perform marginally better.

**Degree of amplitude variation in simulated images has little impact on performance for most tools**

As high signal amplitude variability is prone to producing gently sloped spot count curves that auto-thresholders tend to struggle with in experimental images (Figures 1G-I, 2K-O), we tested whether the degree of amplitude variability in our set of simulated images affected tool performance.

For all tools, trends were similar between maximum recall (Figure S34A) and PR-AUC (Figure S34B). TrueSpot showed a very weak anticorrelation (Table S16, Max recall: Spearman = -0.144, p = $7.63 \times 10^{-9}$, PR-AUC: Spearman = -0.086, p = $5.80 \times 10^{-4}$), and deepBlink with the default model showed a very weak correlation (Max recall: Spearman = 0.093, $p = 2.02 \times 10^{-4}$, PR-AUC: Spearman = 0.078, $p = 1.85 \times 10^{-3}$). Big-FISH also showed a weak anticorrelation, though the p-value for PR-AUC was not within significant range (Max recall: Spearman = -0.174, $p = 2.77 \times 10^{-12}$, PR-AUC: p = 0.092).

RS-FISH showed stronger anticorrelation for both measures than TrueSpot or Big-FISH with both Spearman estimates around -0.28 (Table S16), but the relationship was still subtle enough that it is difficult to visually resolve downward slopes in Figure S34A-B.

The most surprising finding from this analysis was the very strong anticorrelation between amplitude variability and both maximum recall (Spearman = -0.769, $p = 2.19 \times 10^{-219}$) and PR-AUC (Spearman = -0.770, $p = 8.27 \times 10^{-272}$) from deepBlink Retrained. This is interesting because as Figure 3A and Figure S6A demonstrated, the retrained model actually performed worse on Sim-FISH images than the default model. However, as can be seen in Figure S34A-B, performance was actually near perfect for the small number of images with extremely low amplitude variability. The strong effect amplitude variability has on performance from this retrained model provides insight into why the model may have performed so poorly overall. It is possible that we overfitted the model when training. It is also possible that there is an element inherent to deepBlink in either its training or prediction that operates under the assumption that all spots within a given image must be around the same intensity. The training set included images with a variety of simulation parameters including amplitude variability, amplitude levels, and SNR, so it is unlikely that this pattern is due simply to the model only recognizing spots with specific parameters.

Finally, we examined the relationship between amplitude variability and F-Score for the tools with automatic threshold selection (Figure S34C). As with the other measures, TrueSpot showed a mild anticorrelation (Spearman = -0.063, $p = 1.23 \times 10^{-2}$). Conversely, Big-FISH showed an overall mild positive correlation (Spearman = 0.080, $p = 1.30 \times 10^{-3}$), likely due again to the small subpopulation of extreme low performers.

**Using batch-wise fixed thresholds reduces outliers, but has minimal effect on time course measurements**

In the original analysis of these time course data[24], a single fixed threshold value was selected for each of the five replicates in each of the two channels using manual inspection (threshold values are listed in Table

S6). To compare automatic per-image and batch fixed thresholding approaches, we generated 500 simulated images with simulation parameters reflective of the 2 yeast time course channels (250 CY5-Like (CY5L), 250 TMR-Like (TMRL)). We plotted the number of ground truth spots versus detected spots by TrueSpot and Big-FISH using the per-image auto-determined thresholds and fixed thresholds set to the average of the auto-determined thresholds for each of the two simulated channels (Figures S22, S23).

Unsurprisingly, the prevalence and relative distance of outliers was reduced using the fixed thresholds in all four cases (Spearman and Pearson test outputs in Table S8). However, use of the fixed threshold in the case of TrueSpot on the CY5L batch also noticeably pulled the fit line down away from $y = x$ (Figure S22), showing that the integration of the outlier threshold picks was driving the threshold too high for most images in the batch and depressing the detected spot count. This effect was as prominent in the TrueSpot TMRL batch (Figure S23).

In the case of Big-FISH, a handful of extreme overcounts in the CY5L group pulled the mean up so much that when it was used as a fixed threshold, overcounting became more common, though the magnitude less extreme (Figure S22). The situation was similar in the TMRL group (Figure S23), though there was a greater population of initial extremes resulting in less of a dampening of overcount magnitude under the fixed threshold.

We then re-derived on-proportion (Figure S24) and average-per-on measurements (Figure S25) from the experimental time course data using fixed thresholds set to the averages of all per-image thresholds per biological replicate, experiment, and channel (Figures S24 and S25 display results from 0.4M replicate 2).

In Big-FISH, fixing the threshold using any of the three batch averages tended to reduce noisiness (Figure S24A vs. S24B-D, both channels; Figure S25A vs. S25B-D TMR.) In fact, qualitative noise (standard deviation between technical replicates at a time point) reduction was the only apparent difference present between fixed and per-image thresholds for most replicates.

We used the paired t-test to quantitatively compare divergence between time course datasets, testing variable threshold vs. fixed thresholds, fixed thresholds vs. other fixed thresholds, and variable and fixed thresholds vs. reference. We found that while there were many curve comparisons with very low p values, the magnitude of the difference between these "significantly different" curves was usually very small (See Table S9). Thus in most cases, any significant differences present between a variable threshold curve and a fixed threshold curve or between two curves from different fixed thresholds were too small to assert that fixing or not fixing the threshold seriously impacted the biological measurement. The one notable exception to this is the Average per ON curves produced by the output from Big-FISH on the TMR channel images where differences could range from insignificant to 29 transcripts per cell. This is likely an artifact of Big-FISH's extremely wide range of threshold selections made across this image batch (Figures 5B, S21). Interestingly, this did not appear to likewise impact the scores on the counterpart ON proportion curve.

Notably, noise reduction using threshold fixing appeared to only occur in only specific regions of a per-image curve that was noisy to begin with. Additionally, most time points with seemingly high noisiness were unaffected by threshold fixing, indicating that threshold fixing only affected final RNA counts from the most egregious outliers. Furthermore, threshold fixing on a mean skewed by such outliers was not able to increase the resemblance between the noisiest dataset and the manual reference curve, pointing to an inability to for threshold fixing to improve accuracy without manual input. Perhaps use of the set median as a fixed threshold instead of the mean would address this issue, though this too would be contingent upon the distribution of the threshold selections and homogeneity of the image batch.

**Use of automated thresholding alters per-cell spot counts in an unpredictable manner in 2D but not 3D**

Collapsing 3D stacks to 2D maximum intensity projections (MIPs) alters the number of spots found at a given intensity threshold simply by virtue of the number of possible calls being smaller. We suspected that this would affect automatic threshold selection behavior, especially in cases where the shape of the resulting spot count curve was significantly altered. Thus, we compared 2D and 3D per-cell spot counts using both per-image automated thresholding (“variable”) and batch-fixed thresholds.

Indeed, use of automatically selected variable thresholds instead of fixed thresholds did appear to alter spot counts and data distributions for both tools with both the *S. cerevisiae* time course images and the mESC histone mark IF images (Figure S35, Table S5). The trend of 2D counts running lower than 3D counts persisted in the vast majority of cases (H3K4me2 TrueSpot being an interesting exception, though the anomaly did not persist when out-of-focus z-slices were removed (Figures S13, S35)). However, the impact automated thresholding had on per-cell spot count distributions differed between the two tools.

For Big-FISH, usage of automated threshold selection tended to increase correlation (as shown by mildly increasing Pearson estimates in Table S5) as well as agreement between 2D and 3D spot counts (Figure S35). Usage of automated thresholding also consistently increased spot counts relative to usage of fixed thresholds for all tested groups in 2D (Figure S36). This does not hold true in 3D – to the contrary, spot counts tended to *decrease* in time course groups when automated thresholding was used. The latter observation is likely merely the result of the fixed threshold used for the time course groups being derived from the mean automatically selected threshold for each biological replicate batch and Big-FISH's propensity to produce extreme outliers (Figures 4,6,S14-S17, S28-S29). What is more interesting is the difference in distribution shapes between 2D and 3D plots of variable versus fixed threshold counts (Figure S36). In 3D, these plots are more spread out and tend to stay vaguely linear, but in 2D the points tend to be more clustered, often forming vertical lines on the left edge of the graph at time points when expression is expected to be low. This is indicative of disproportionate overcalling with automated thresholding relative to fixed threshold use in 2D, but not as much in 3D.

Similar clustering trends can be seen in the TrueSpot results as well. However, the most striking feature of the TrueSpot data is that use of automated thresholding on 2D MIPs greatly increases the overall noisiness of the spot count distributions in a way that is not seen in 3D (Figures S35-S26, Table S5). This seems potentially indicative of relatively erratic threshold selection in 2D that does not occur in 3D. Indeed, we observed many MIP spot count curves with little to no initial dropoff, a feature that is not in their 3D counterparts (Figure S37). Such curves are produced when there is relatively little background noise to compare against the signal, a phenomenon we also frequently observed with particularly clean simulated images (Figure S3). Lack of background information to compare against can increase the difficulty of selecting a threshold, particularly for an automated thresholder.

**Supplementary Figures**

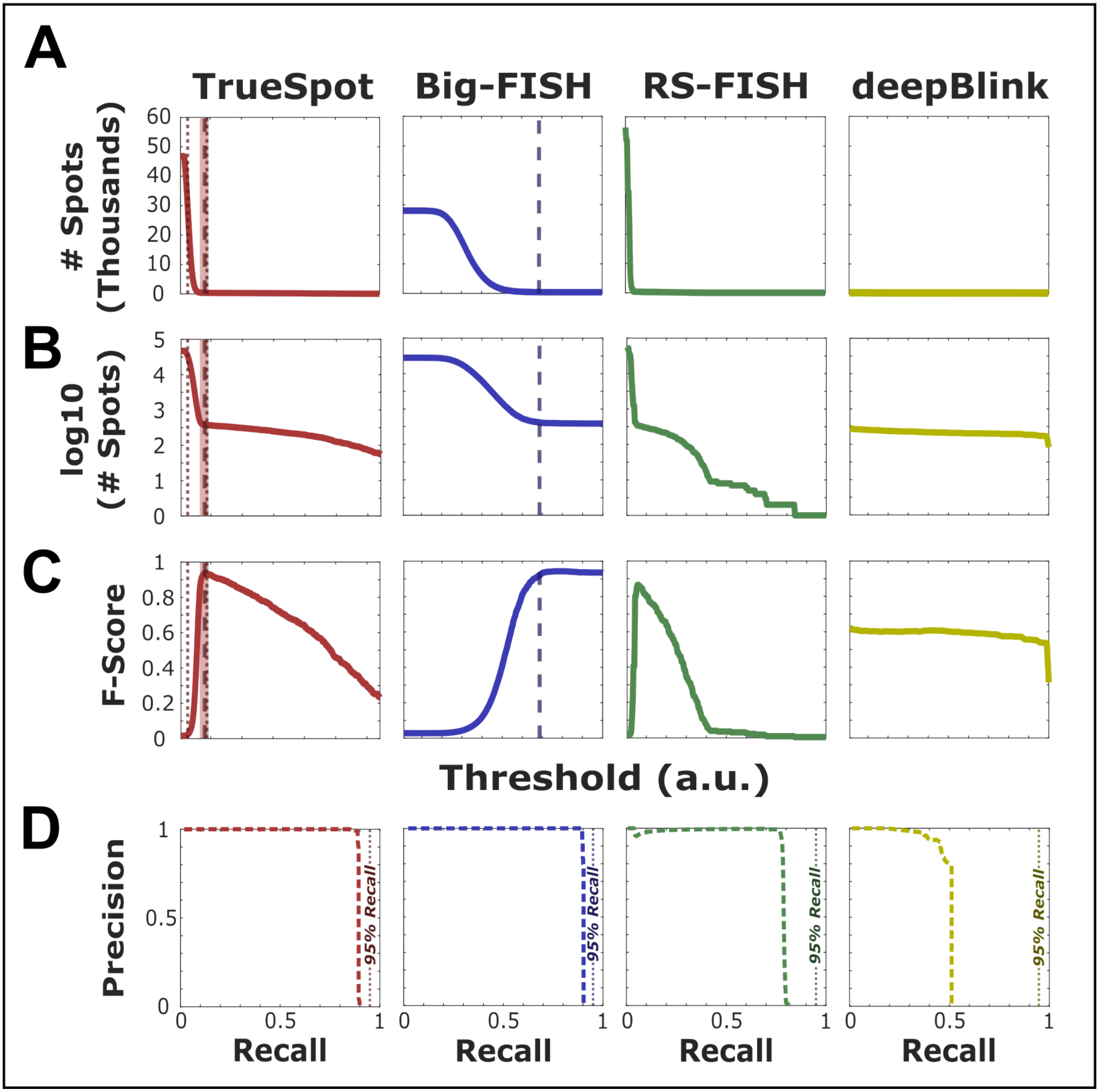


**Figure S1. Simulated images tend to produce clean, sharp spot count curves.** (A-D) Example spot count ((A) linear scale and (B) logarithmic scale), (C) F-Score, and (D) Precision-Recall curves produced by running each tool (TrueSpot (red), Big-FISH (blue), RS-FISH (green), deepBlink using default smiFISH model (yellow)) on a single example simulated image. Threshold values are arbitrary and only comparable between tools in that they range from lenient (lower value) to strict (higher value). Automatically selected threshold values are shown by dashed vertical line for TrueSpot and Big-FISH. TrueSpot mean ± standard deviation (shaded rectangle), minimum, and maximum (dotted lines) of threshold suggestion pool additionally shown.

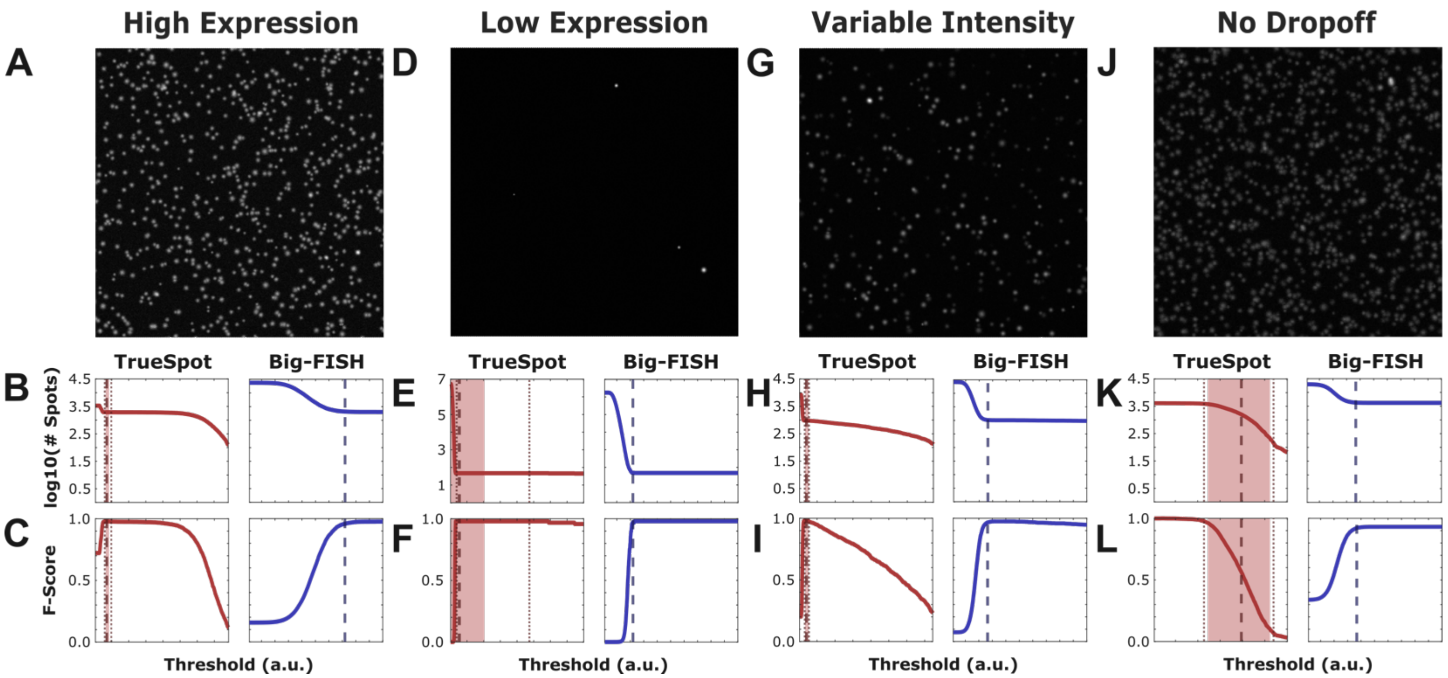


**Figure S2. Simulated images produce cleaner filtered outputs and sharper curves than experimental images.** Samples of images with simulated puncta spots from each of the three experimental image archetypes: High expression (A-C), Low Expression (D-F), and Variable Intensity (G-I). Additionally, there is a fourth type ("No Dropoff", J-L) observed only among simulated images. (B,E,H,K) Spot count curves (logarithmic projection) and (C,F,I,L) F-Score curves for both TrueSpot (left, red) and Big-FISH (right, blue). Dashed lines demarcate automatically selected threshold. Shaded area in TrueSpot graphs represents mean of suggestion pool ± standard deviation of thresholds, dotted lines represent pool minimum and maximum.

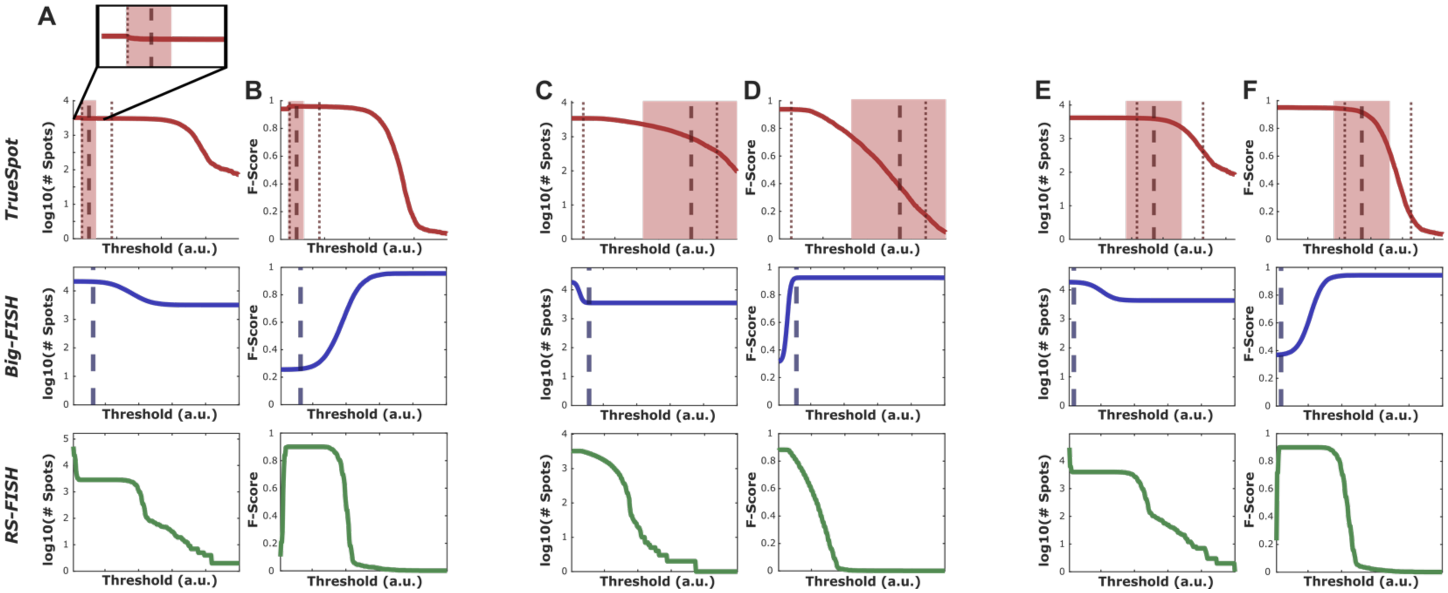


**Figure S3. Curves with no initial dropoff seen by TrueSpot still display variation.** (A,C,E) Spot count (y logarithmic scaled) and (B,D,F) F-Score curves produced by TrueSpot (top, red), Big-FISH (middle, blue), and RS-FISH (bottom, green) for three "no dropoff" example images. (A-B) TrueSpot's curve starts at the low end of the signal range. It contains a very small notch at the lower end, which is detected by the automatic thresholder. Big-FISH sees the signal to noise transition as shorter, more gradual curve, but its automated thresholder does not appear to recognize it, instead choosing a far too lenient threshold. Initial sharp dropoff is seen by RS-FISH, but neither of the others. (C-D) Big-FISH is the only one of the three tools to see a sharp noise to signal transition which its thresholder is able to identify. TrueSpot's and RS-FISH's lowest thresholds fall well into the signal range. TrueSpot's threshold selection is very uncertain and of low quality due to the lack of noise to compare against. (E-F) As with (A-B), TrueSpot and Big-FISH do not see a sharp noise to signal transition and Big-FISH's threshold selector is unable to identify the gradual transition curve. TrueSpot sees a much longer signal plateau than in (C) and no notch as in (A), leading to a threshold prediction that falls in between (B) and (D) in terms of certainty and quality.

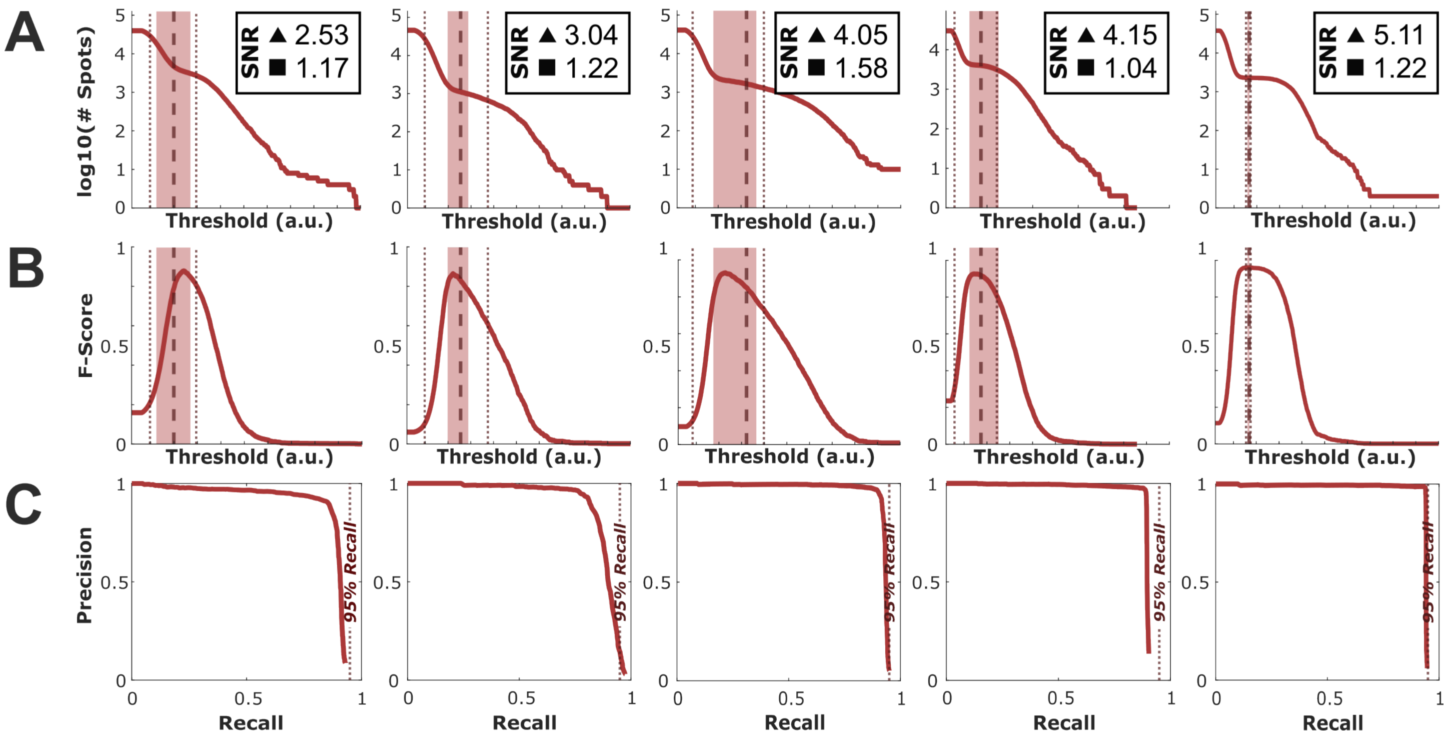


**Figure S4. TrueSpot performs well on simulated images with low SNR.** (A) The spot count (logarithmic projection), (B) F-Score, and (C) PR curves produced by TrueSpot for five example simulated images with low SNR. SNR measurements indicated in inset boxes for each column. SNR measure denoted by triangle is $\mu_A/\sigma_B$ (mean signal amplitude / background standard deviation), and measure denoted by square is $\mu_A/\mu_B$ (mean signal amplitude / mean background level). Dashed lines indicate automatically selected threshold, shaded regions represent mean value of threshold suggestion pool ± standard deviation, and dotted lines represent pool minimum and maximum. Dotted line in PR graphs marks 95% recall.

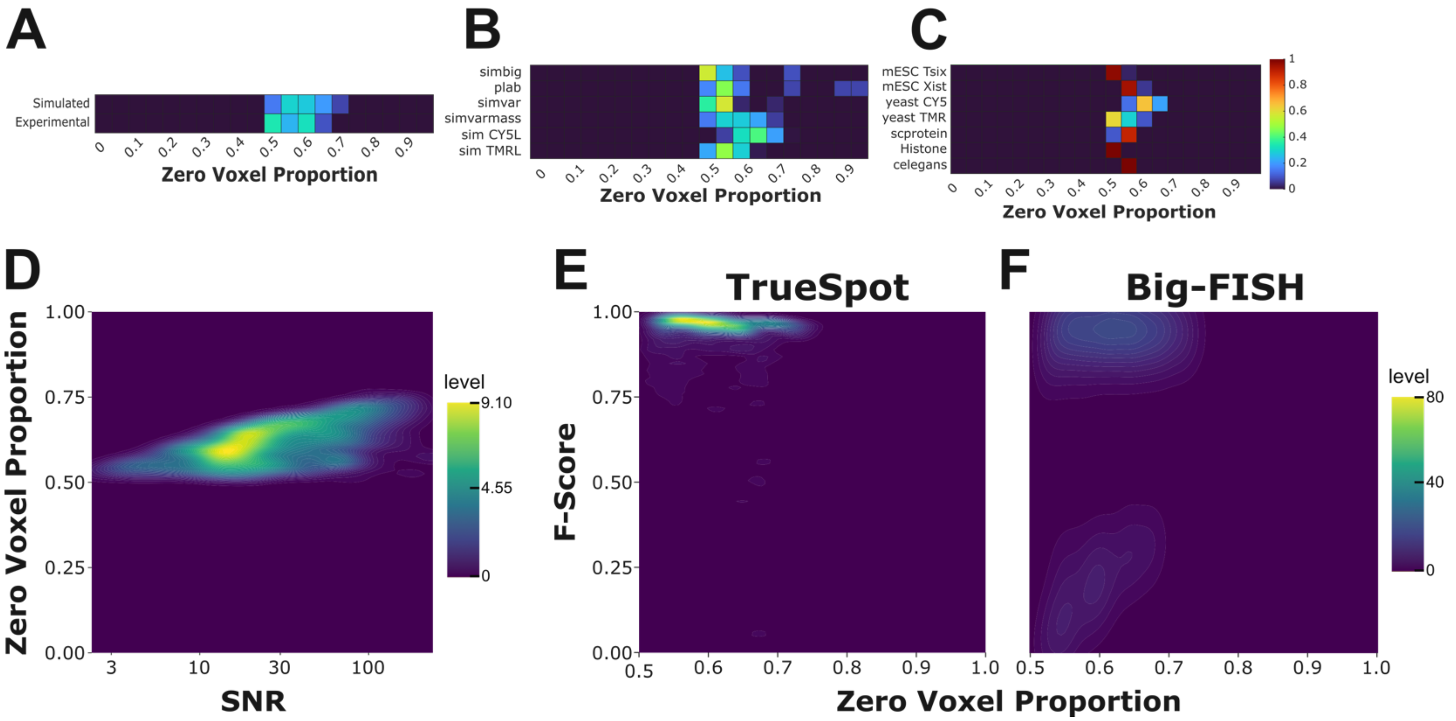


**Figure S5. Simulated images tend to produce cleaner filtered outputs than experimental images.** (A-C) A breakdown of test images by zero voxel proportion after applying the TrueSpot LoG filter as a proxy measurement for image "cleanliness". (A) Split into simulated and experimental images. (B) Simulated images split by batch. (C) Experimental images split by group. (D) A density map of the zero voxel proportion of simulated images (n = 1590) plotted against SNR (log scale). (E-F) Density maps of F-Score (tools with automatic thresholding) versus ZVP for all simulated images. Low F-Score outliers tend to have high ZVP for (E) TrueSpot and low ZVP for (F) Big-FISH.

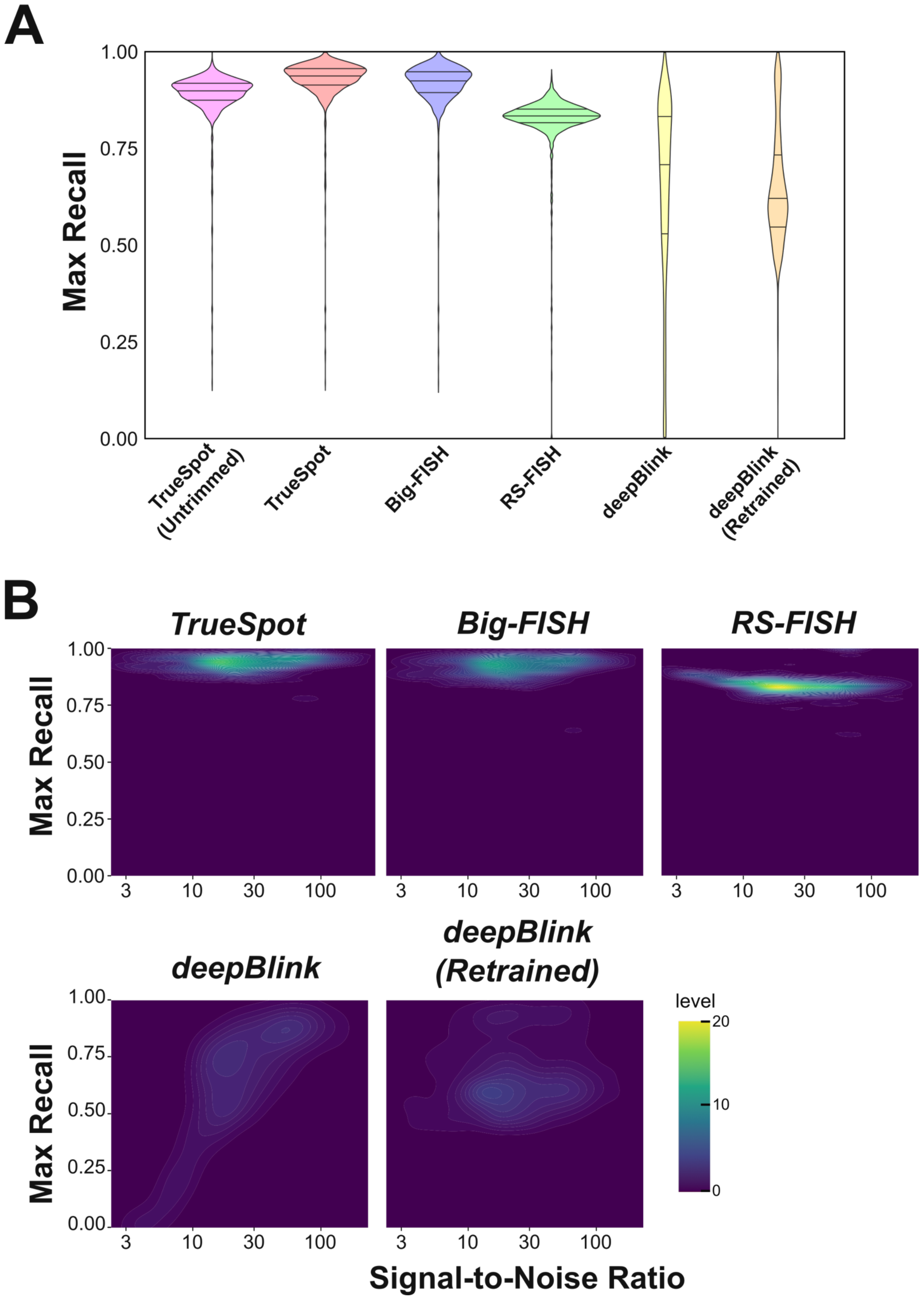
A
Max Recall
1.00
0.75
0.50
0.25
0.00
TrueSpot (Untrimmed)
TrueSpot
Big-FISH
RS-FISH
deepBlink
deepBlink (Retrained)
B
TrueSpot
Big-FISH
RS-FISH
deepBlink
deepBlink (Retrained)
Max Recall
3
10
30
100
level
20
10
0
Signal-to-Noise Ratio

1

**Figure S6. Maximum recall distributions tend to reflect PR-AUC distributions.** Distributions of maximum recall values for tested tools (ZVP ≤ 0.7) (A) Violin plots of maximum recall values across all Sim-FISH simulated images (n = 1472, deepBlink Retrained n = 1367) for all tools and TrueSpot untrimmed. (B) Plots of maximum recall versus signal-to-noise ratio (SNR, calculated as $\mu_{Signal} / \sigma_{Background}$ from simulation parameters). SNR axis is on a logarithmic scale. Mann-Whitney and Spearman correlation calculations are shown in Tables S1A and S14A respectively.

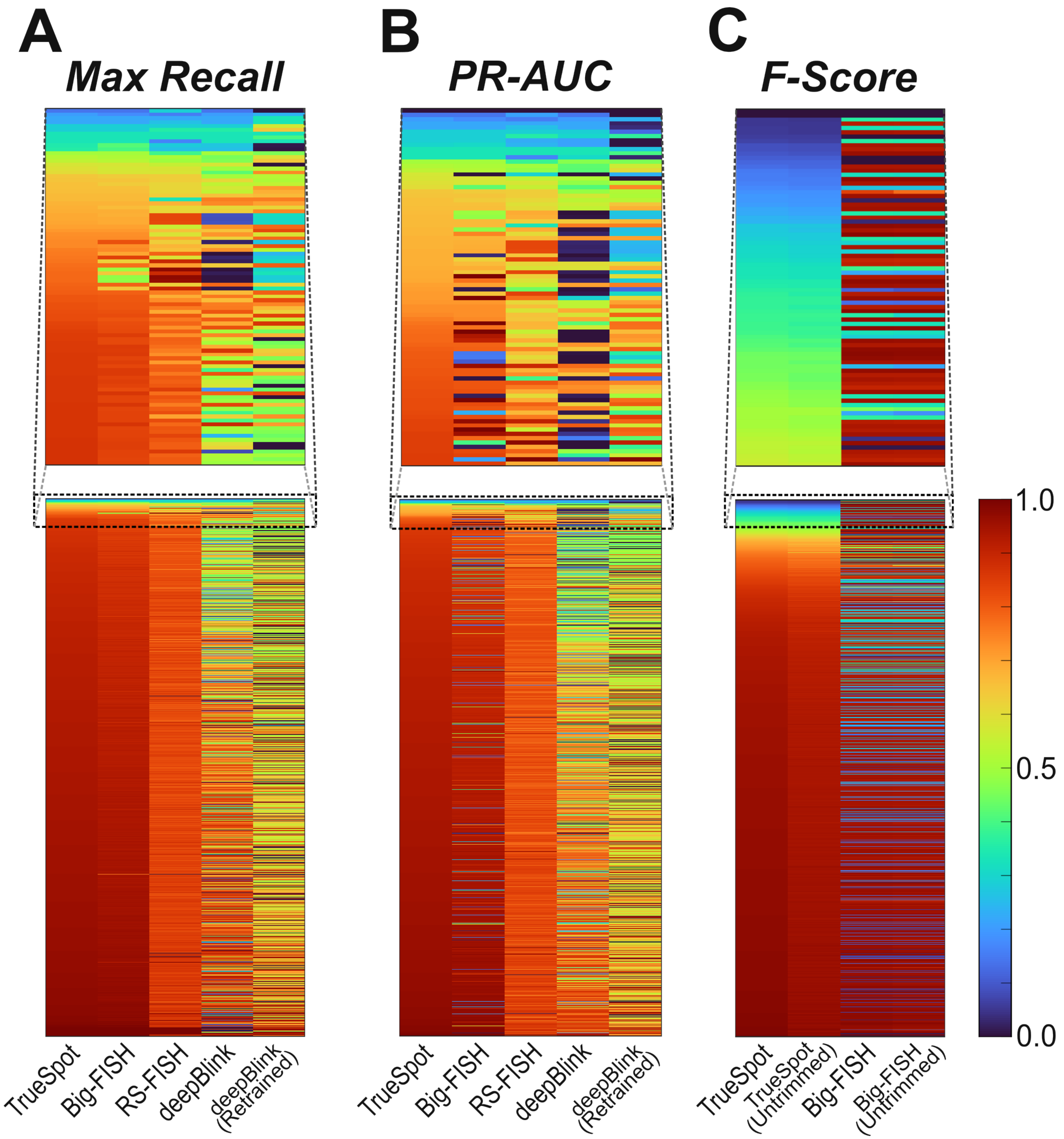


**Figure S7. A subset of simulated images score poorly on performance metrics regardless of tool.** Heatmaps of maximum recall (A), PR-AUC (B), and F-Score (C) for each individual simulated image (rows, n = 1599, no ZVP cap) and each tool tested (columns). Rows are sorted by TrueSpot (trimmed) score independently for each heatmap (ie. rows are not in the same order between A, B, and C).

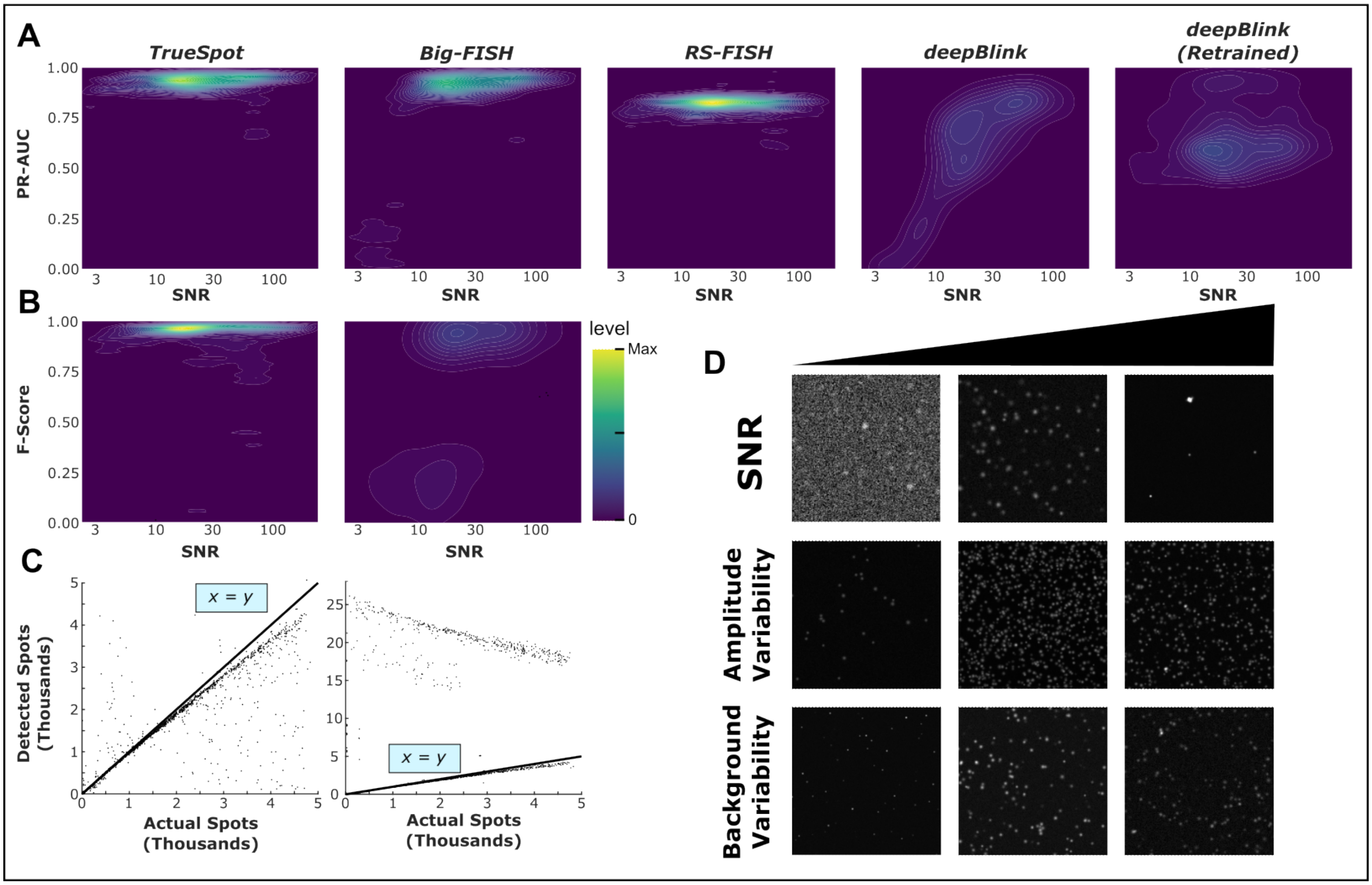


**Figure S8. Specific image properties may or may not impact tool performance.** (A) Density maps of PR-AUC versus SNR ($\mu_{Signal}$ / $\sigma_{Background}$, scaled logarithmically for visualization) for all tested simulated images (ZVP ≤ 0.7, n = 1472), four tools and deepBlink retrained (n = 1369). Spearman correlations are shown in Table S14B. (B) Density maps of F-Score at automatically selected threshold versus SNR for all tested simulated images (ZVP ≤ 0.7), only tools with automatic threshold selection. Spearman correlations are shown in Table S14C. (C) Scatter plots of number of actual spots generated for each simulated image versus number of spots detected by each tool with automatic thresholding. *x* = *y* line marked for reference. (D) Examples of simulated images across ranges of SNR, amplitude variability, and background variability.

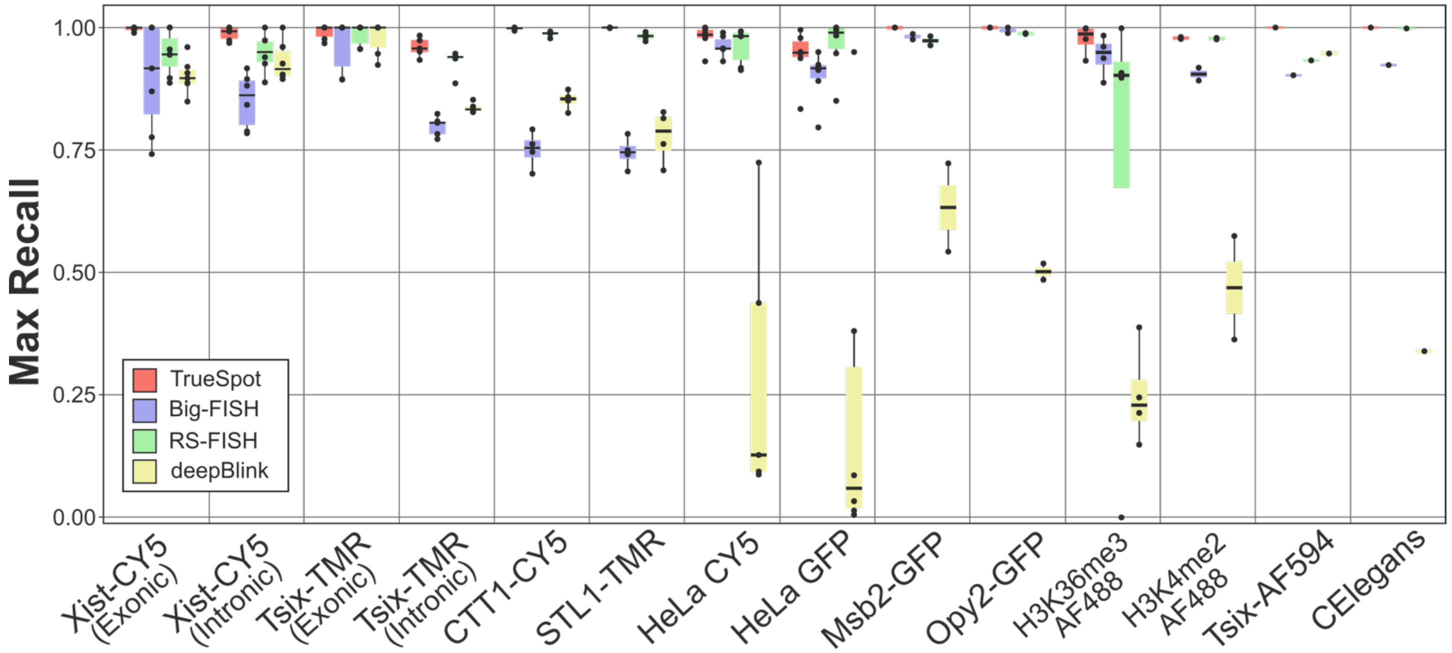


**Figure S9. Maximum recall of all tools trends high on experimental RNA-FISH images.** Box-and-whiskers plot for maximum recall on benchmarked experimental images. Box boundaries represent 25th and 75th percentiles, midline represents median, and whiskers represent furthest values within 1.5 * interquartile range (IQR) from box edges. Spots represent individual images tested. Counts and types for image groups can be seen in Figure 3C-D. Statistics are shown in Table S2A.

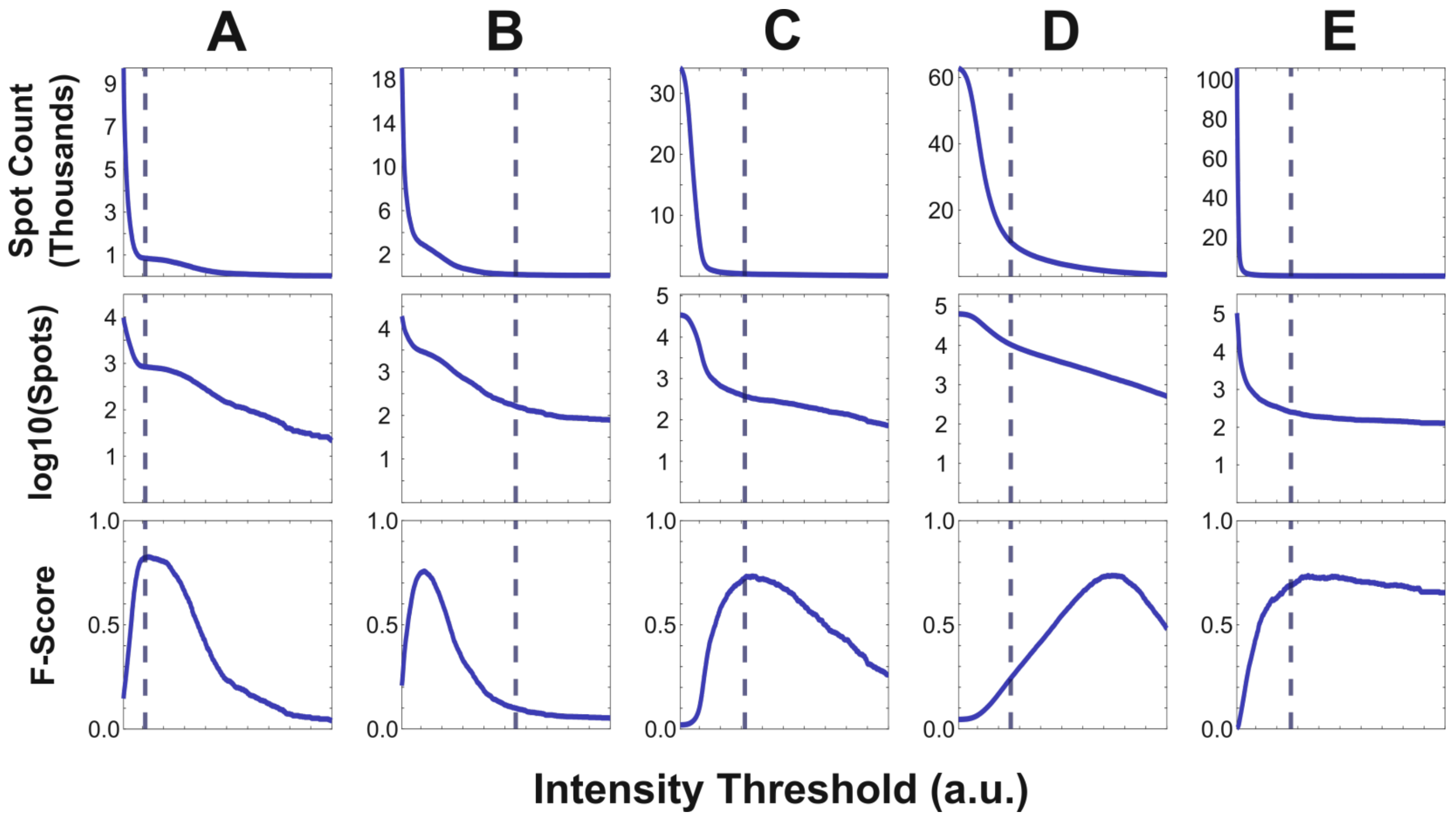


**Figure S10. Curve attributes that lead to extreme threshold selections by Big-FISH are not easily defined.** Examples of images with “gradual bend” curves produced by Big-FISH illustrating variable performance in threshold selection despite apparently similar curve shapes. (A-C) STL1-TMR, (D) H3K36me3-AF488, (E) Tsix-TMR (Intronic).

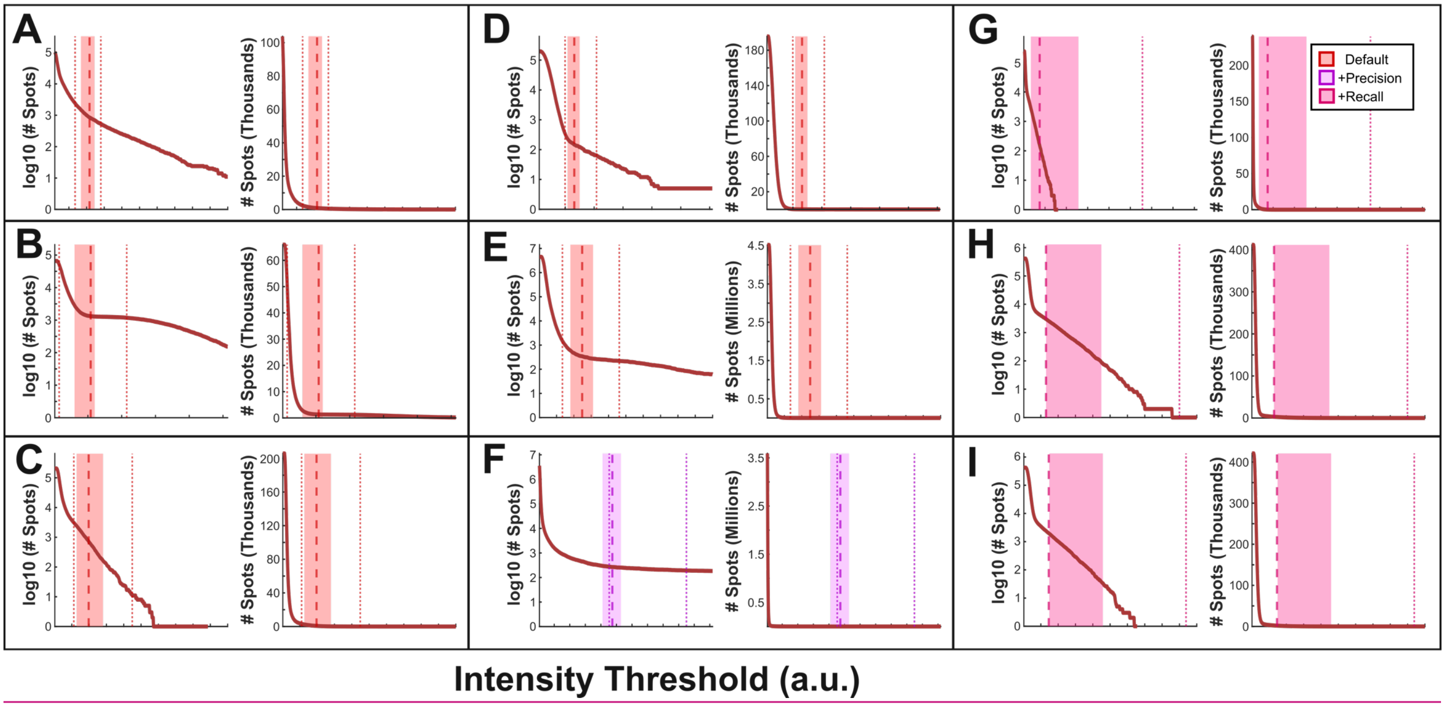


**Figure S11. TrueSpot's threshold selection algorithm consistently finds the approximate center of gradual curves at default settings.** Examples of spot count curves (log scale, left; linear scale, right) and threshold selections produced by TrueSpot for experimental images with high amplitude variability. Examples from six different experimental groups were used: (A) Msb2-GPF, (B) CElegans, (C) HeLa GFP, (D) HeLa CY5, (E) Tsix-TMR (Exonic), (F) Tsix-TMR (Intronic), (G) H3K36me3-AF488, (H-I) H3K4me2-AF488. Range colors reflect thresholder preset.

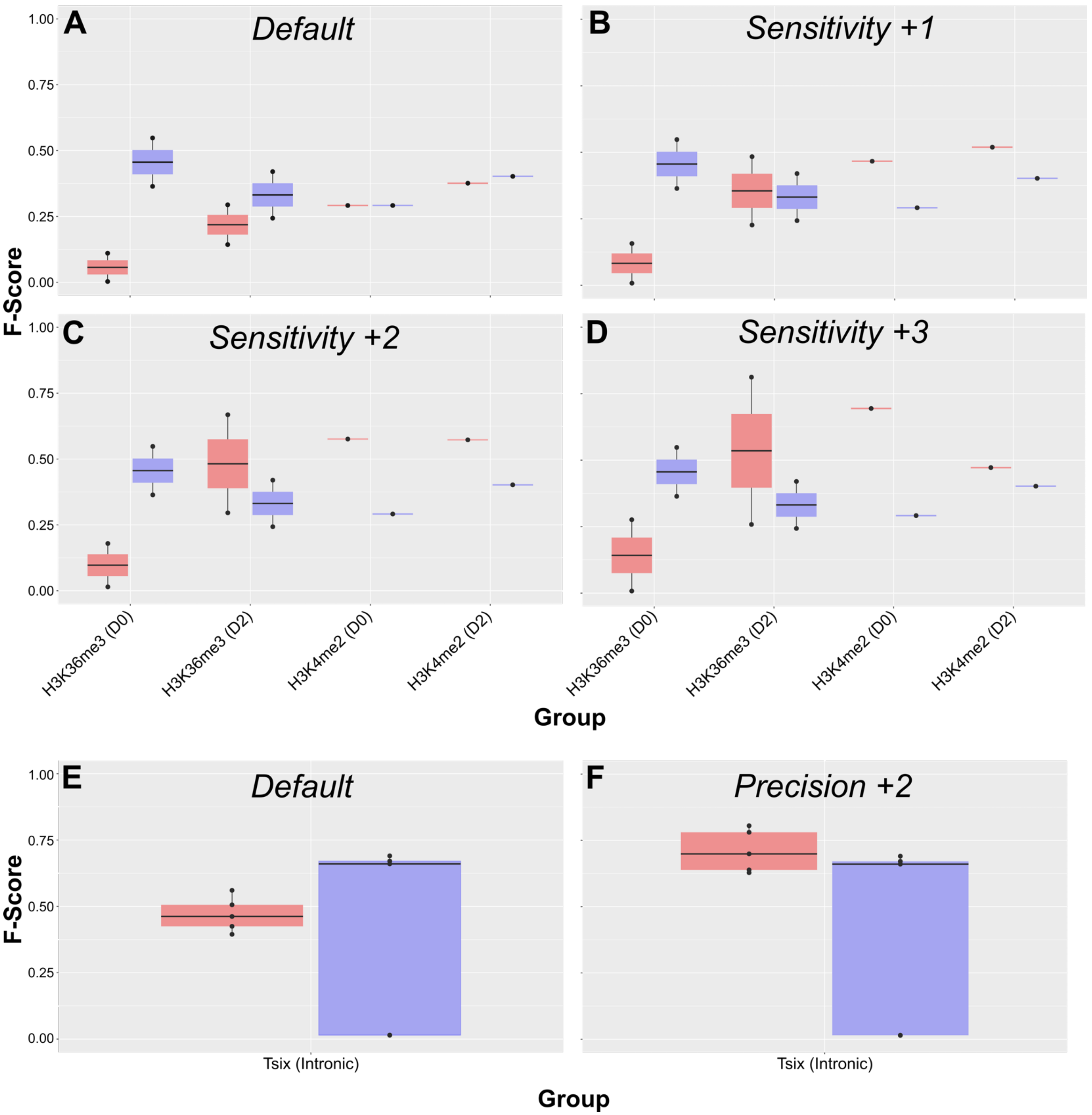


**Figure S12. Thresholder preset choice affects TrueSpot's thresholding performance.** Comparison of F-Scores at automatically selected threshold over a range of threshold presets for the four histone subgroups (A-D) and Tsix Intronic group (E-F). TrueSpot is shown in red. Big-FISH is included in blue for reference.

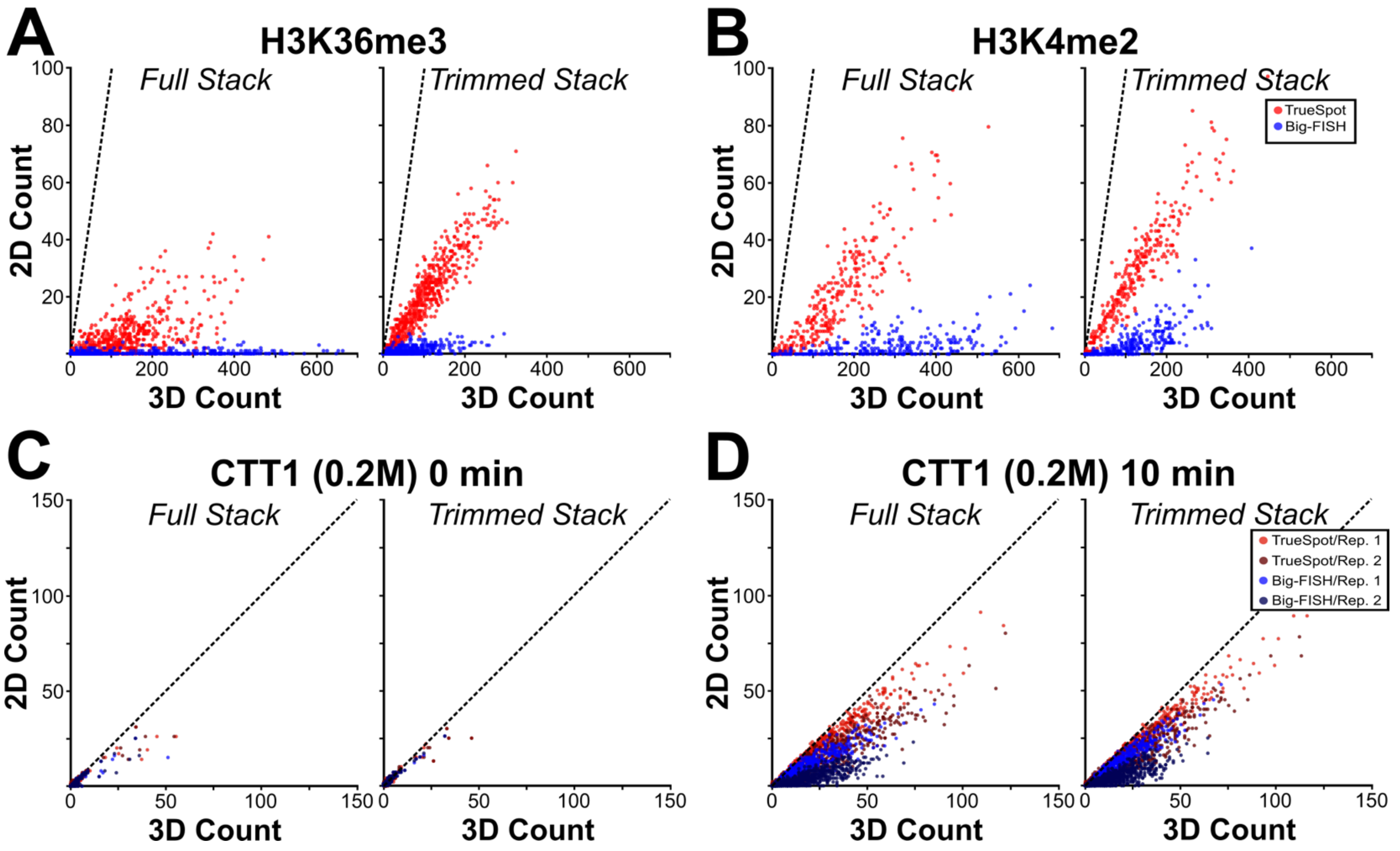


**Figure S13. Using maximum intensity projections from trimmed stacks does not resolve undercounting.** Per-cell 2D/maximum intensity projection versus 3D spot counts across both histone groups (A-B) and two sample CTT1 (0.2M) time points (C-D) using both full and trimmed image stacks (fixed thresholds). Dashed lines denote $y = x$. Both biological replicates are included for CTT1.

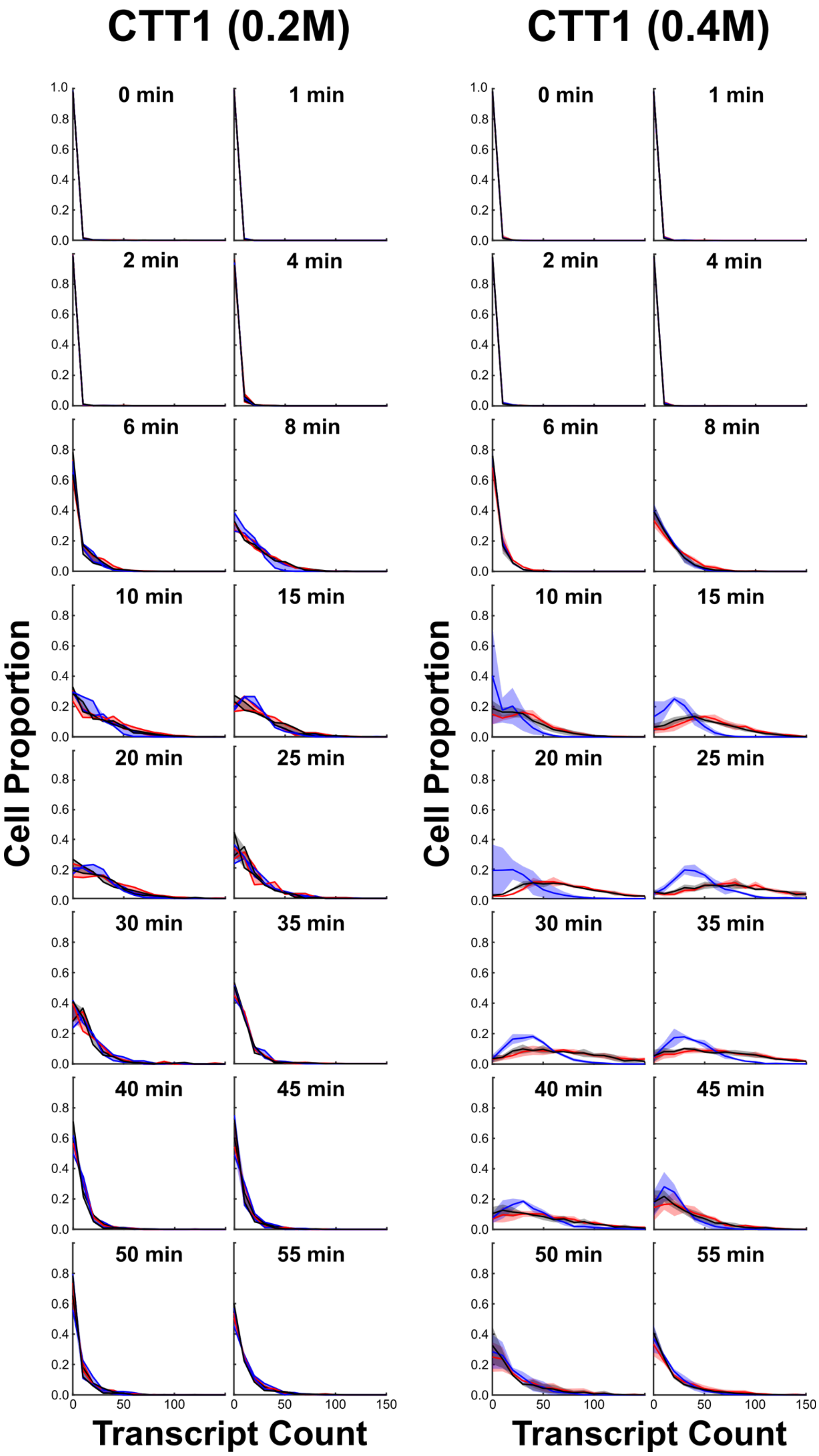
CTT1 (0.2M)
CTT1 (0.4M)
0 min
1 min
2 min
4 min
6 min
8 min
10 min
15 min
20 min
25 min
30 min
35 min
40 min
45 min
50 min
55 min
Cell Proportion
Transcript Count
0
50
100
150
0.0
0.2
0.4
0.6
0.8
1.0

1

**Figure S14. TrueSpot and Big-FISH tend to show agreement on per-cell transcript count with manually thresholded reference.** Probability distribution plots (bin size = 10) of transcript/spot count per cell for CTT1 time course experiments after 0.2M (n = 2 biological replicates) and 0.4M (n = 3) applied hyperosmotic stress. Distributions shown for TrueSpot with auto-thresholding (red), Big-FISH with auto-thresholding (blue), and previously published reference derived from TrueSpot prototype with manual thresholding (black). (0.2M) Lines represent measures from biological replicates, fill covers region in between the two replicates. (0.4M) Lines represent mean measures of biological replicates, fill represents standard deviation.

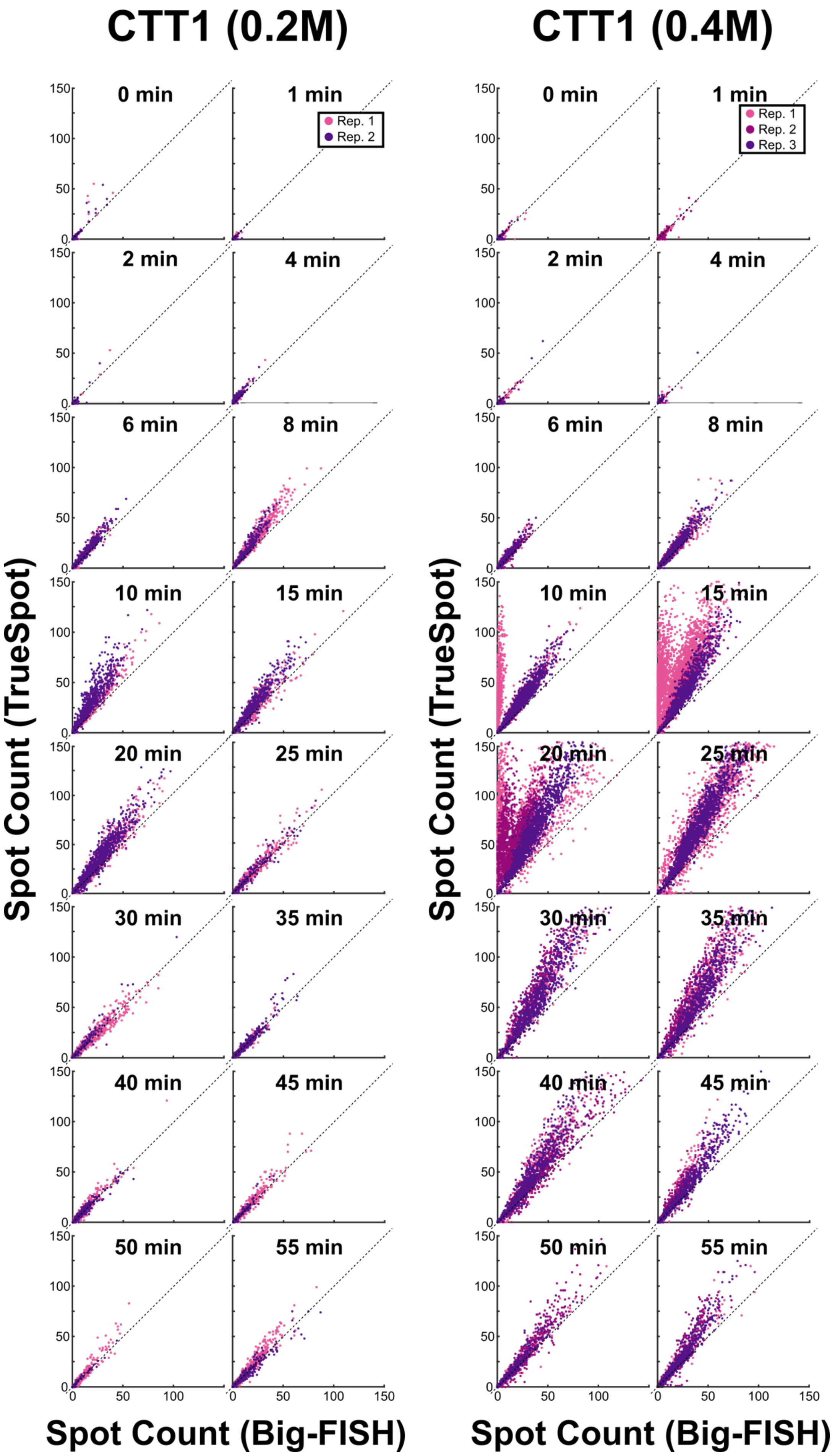
CTT1 (0.2M)
CTT1 (0.4M)
0 min
1 min
2 min
4 min
6 min
8 min
10 min
15 min
20 min
25 min
30 min
35 min
40 min
45 min
50 min
55 min
Rep. 1
Rep. 2
Rep. 3
Spot Count (TrueSpot)
Spot Count (Big-FISH)
0
50
100
150

1

**Figure S15. TrueSpot and Big-FISH tend to agree on per-cell CTT1 transcript count.** Scatterplots of per-cell CTT1 transcript/spot count produced by TrueSpot with auto-thresholding versus Big-FISH with auto-thresholding at each time point. Each point represents a single cell, point fill color reflects biological replicate (0.2M NaCl stimulation: n = 2; 0.4M: n = 3). $y = x$ line marked for each scatterplot.

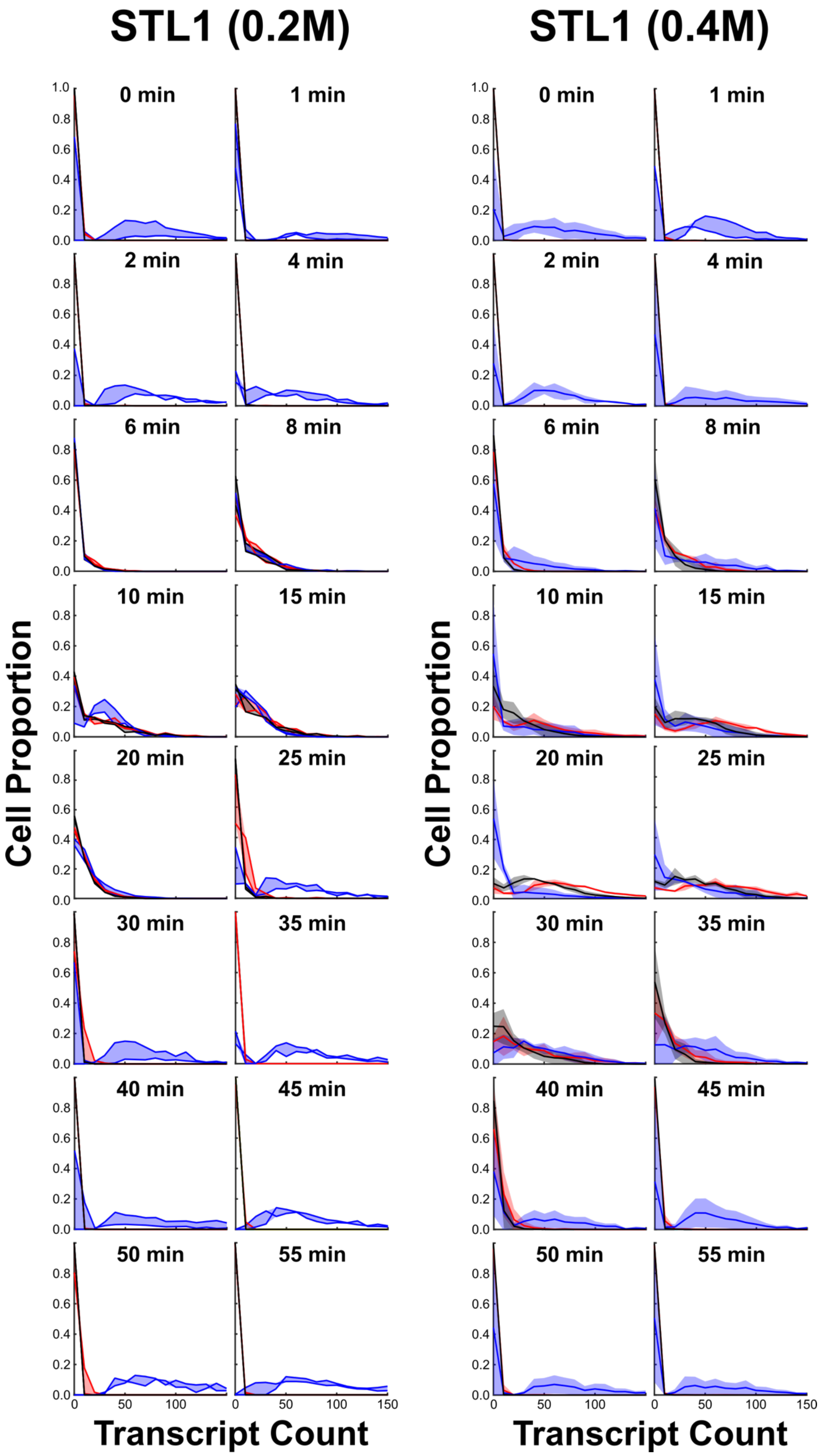

STL1 (0.2M)
STL1 (0.4M)
0 min
1 min
2 min
4 min
6 min
8 min
10 min
15 min
20 min
25 min
30 min
35 min
40 min
45 min
50 min
55 min
Cell Proportion
Transcript Count


1

**Figure S16. Big-FISH calls far more STL1 spots per cell than TrueSpot or manual thresholder at most time points.** Probability distribution plots (bin size = 10) of transcript/spot count per cell for STL1 time course experiments after 0.2M (n = 2 biological replicates) and 0.4M (n = 3) applied hyperosmotic stress. Distributions shown for TrueSpot with auto-thresholding (red), Big-FISH with auto-thresholding (blue), and previously published reference derived from TrueSpot prototype with manual thresholding (black). (0.2M) Lines represent measures from biological replicates, fill covers region in between the two replicates. (0.4M) Lines represent mean measures of biological replicates, fill represents standard deviation.

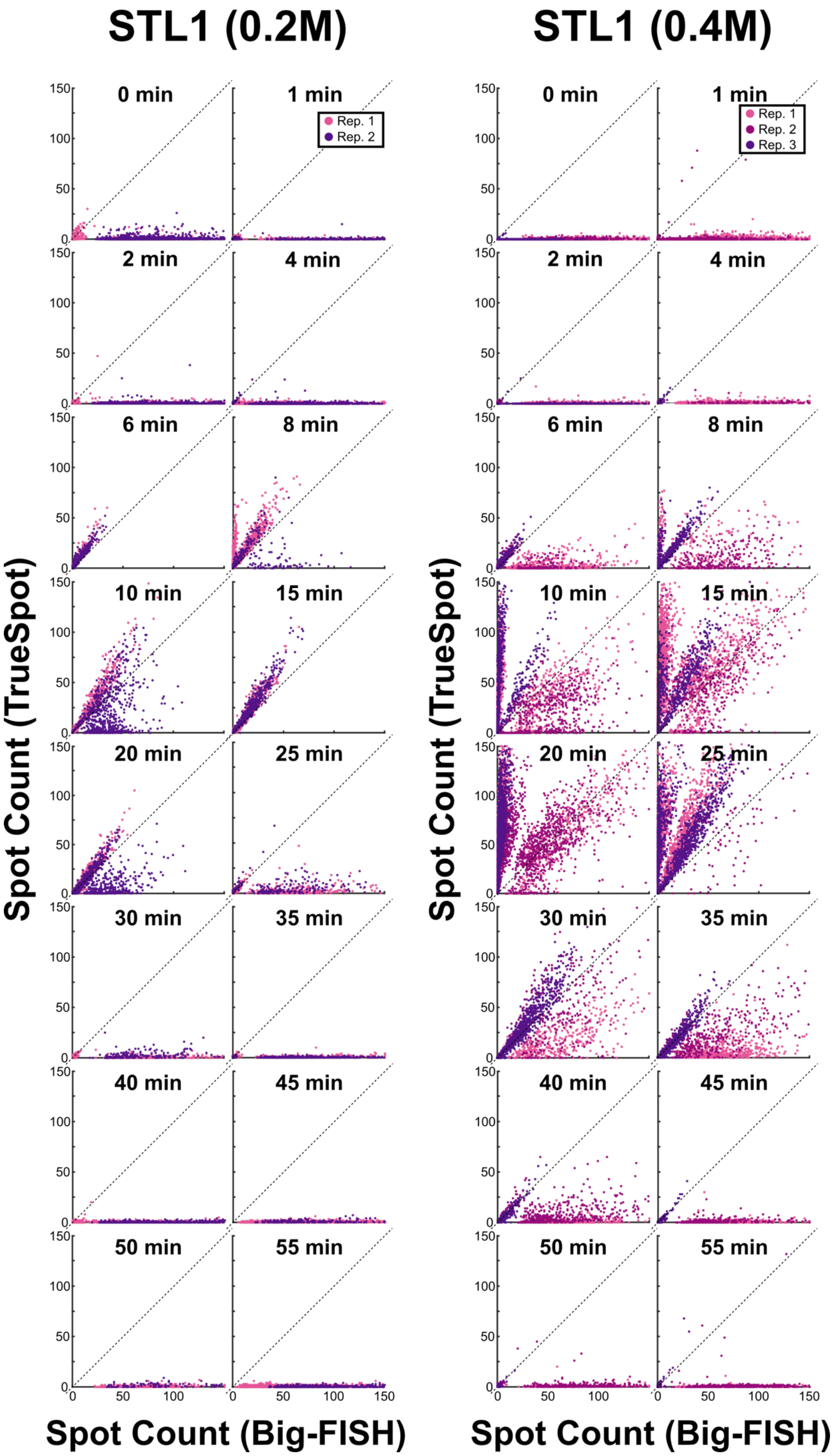

STL1 (0.2M)
STL1 (0.4M)
0 min
1 min
2 min
4 min
6 min
8 min
10 min
15 min
20 min
25 min
30 min
35 min
40 min
45 min
50 min
55 min
Rep. 1
Rep. 2
Rep. 3
Spot Count (TrueSpot)
Spot Count (Big-FISH)
0
50
100
150


1

**Figure S17. Big-FISH tends to overcall per-cell STL1 spots relative to TrueSpot.** Scatterplots of per-cell STL1 transcript/spot count produced by TrueSpot with auto-thresholding versus Big-FISH with auto-thresholding at each time point. Each point represents a single cell, point fill color reflects biological replicate (0.2M NaCl stimulation: n = 2; 0.4M: n = 3). $y = x$ line marked for each scatterplot.

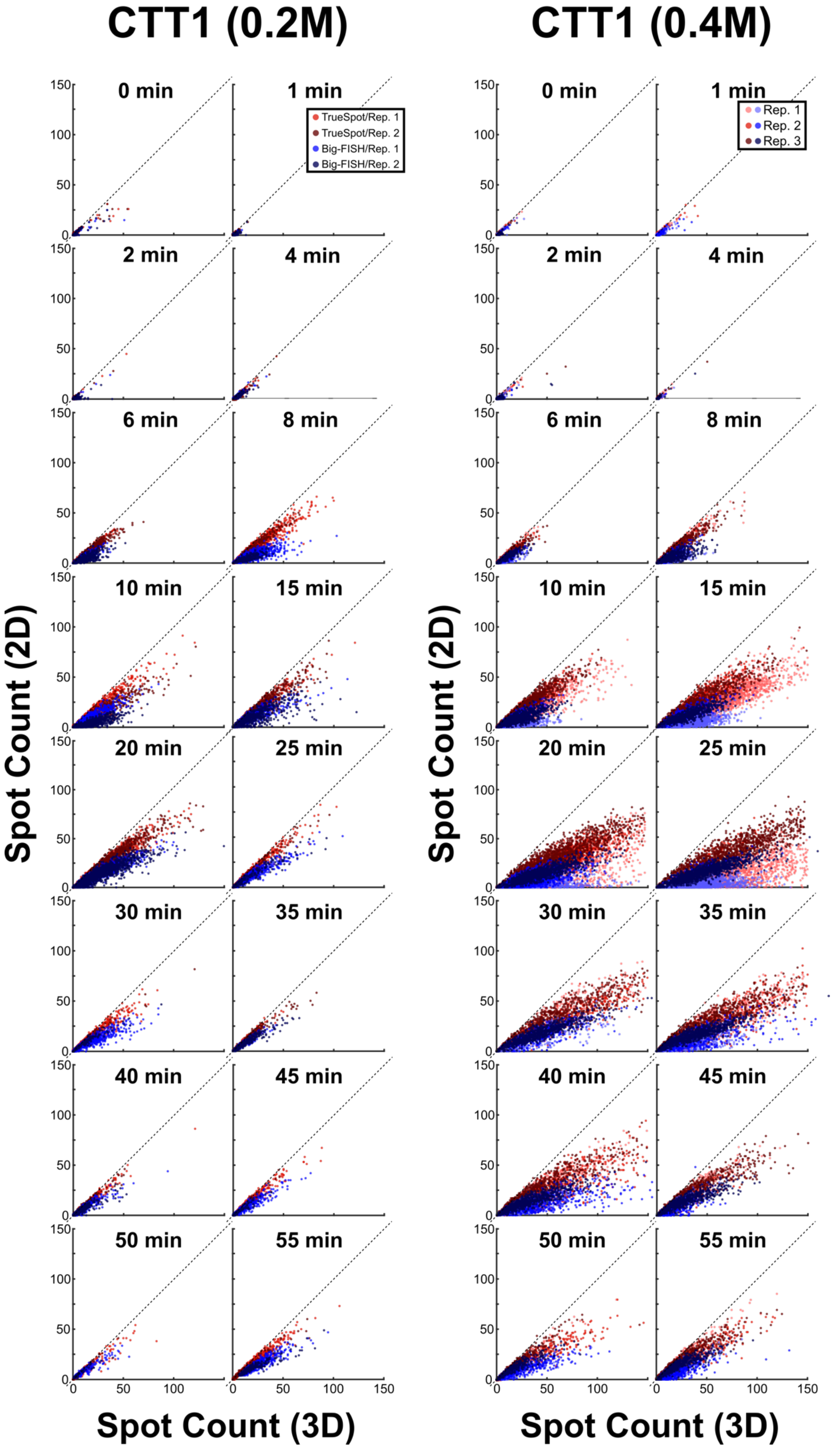
CTT1 (0.2M)
CTT1 (0.4M)
0 min
1 min
2 min
4 min
6 min
8 min
10 min
15 min
20 min
25 min
30 min
35 min
40 min
45 min
50 min
55 min
TrueSpot/Rep. 1
TrueSpot/Rep. 2
Big-FISH/Rep. 1
Big-FISH/Rep. 2
Rep. 1
Rep. 2
Rep. 3
Spot Count (2D)
Spot Count (3D)
0
50
100
150

**Figure S18. Per-cell spot counts from maximum intensity projections are lower and have a possible non-linear relationship with 3D counterparts.** Scatterplots of per-cell spot counts derived from 2D maximum intensity projections plotted against spot counts from analysis of full 3D image stack for all CTT1 time points. Stacks were not trimmed. Fixed thresholds. Each point represents a single cell, fill color represents tool and biological replicate (0.2M n = 2, 0.4M n = 3).

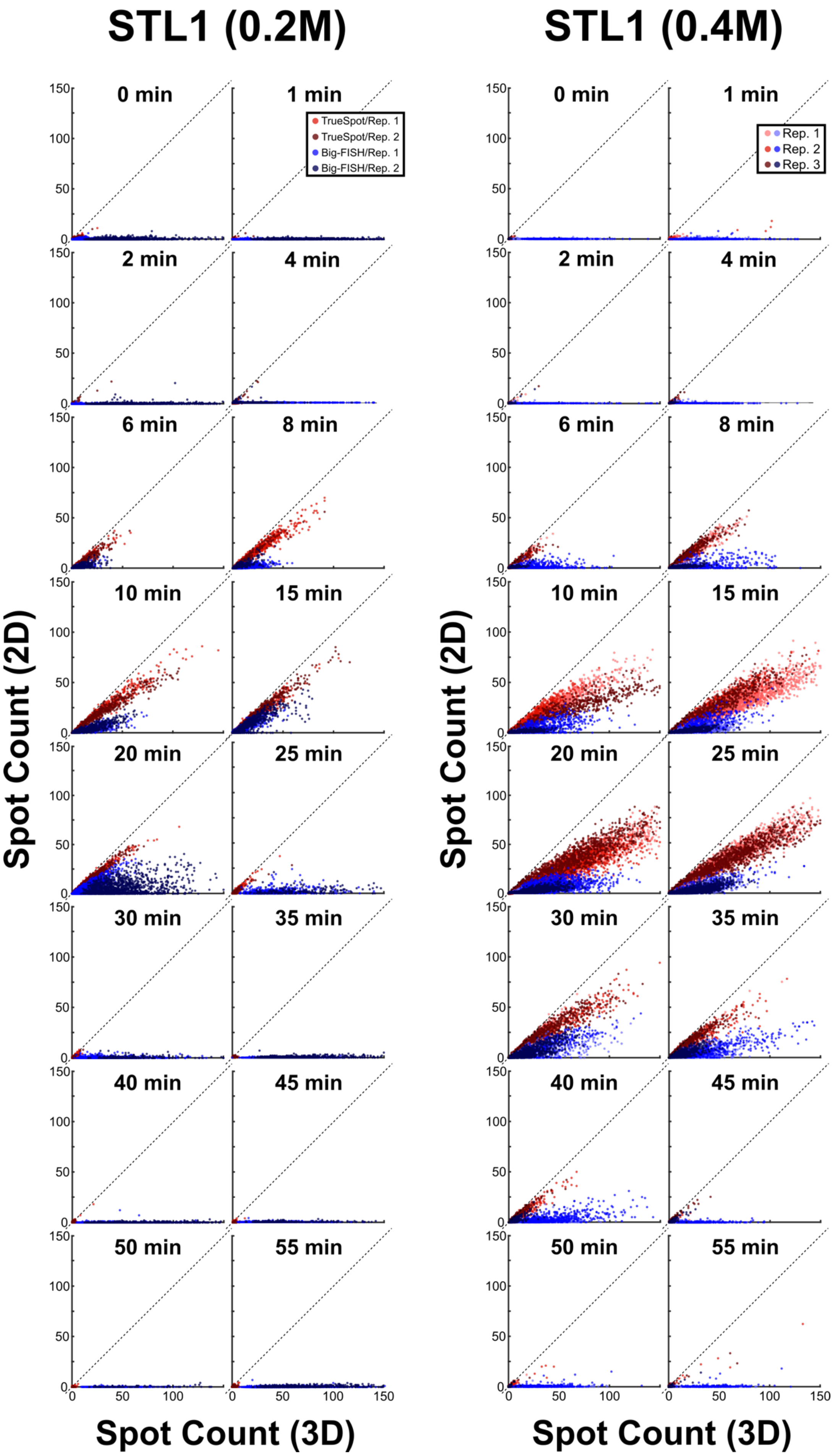

STL1 (0.2M)
STL1 (0.4M)
0 min
1 min
2 min
4 min
6 min
8 min
10 min
15 min
20 min
25 min
30 min
35 min
40 min
45 min
50 min
55 min
TrueSpot/Rep. 1
TrueSpot/Rep. 2
Big-FISH/Rep. 1
Big-FISH/Rep. 2
Rep. 1
Rep. 2
Rep. 3
Spot Count (2D)
Spot Count (3D)


1

**Figure S19. Per-cell spot counts from maximum intensity projections are lower and have a possible non-linear relationship with 3D counterparts.** Scatterplots of per-cell spot counts derived from 2D maximum intensity projections plotted against spot counts from analysis of full 3D image stack for all STL1 time points. Stacks were not trimmed. Fixed thresholds. Each point represents a single cell, fill color represents tool and biological replicate (0.2M n = 2, 0.4M n = 3).

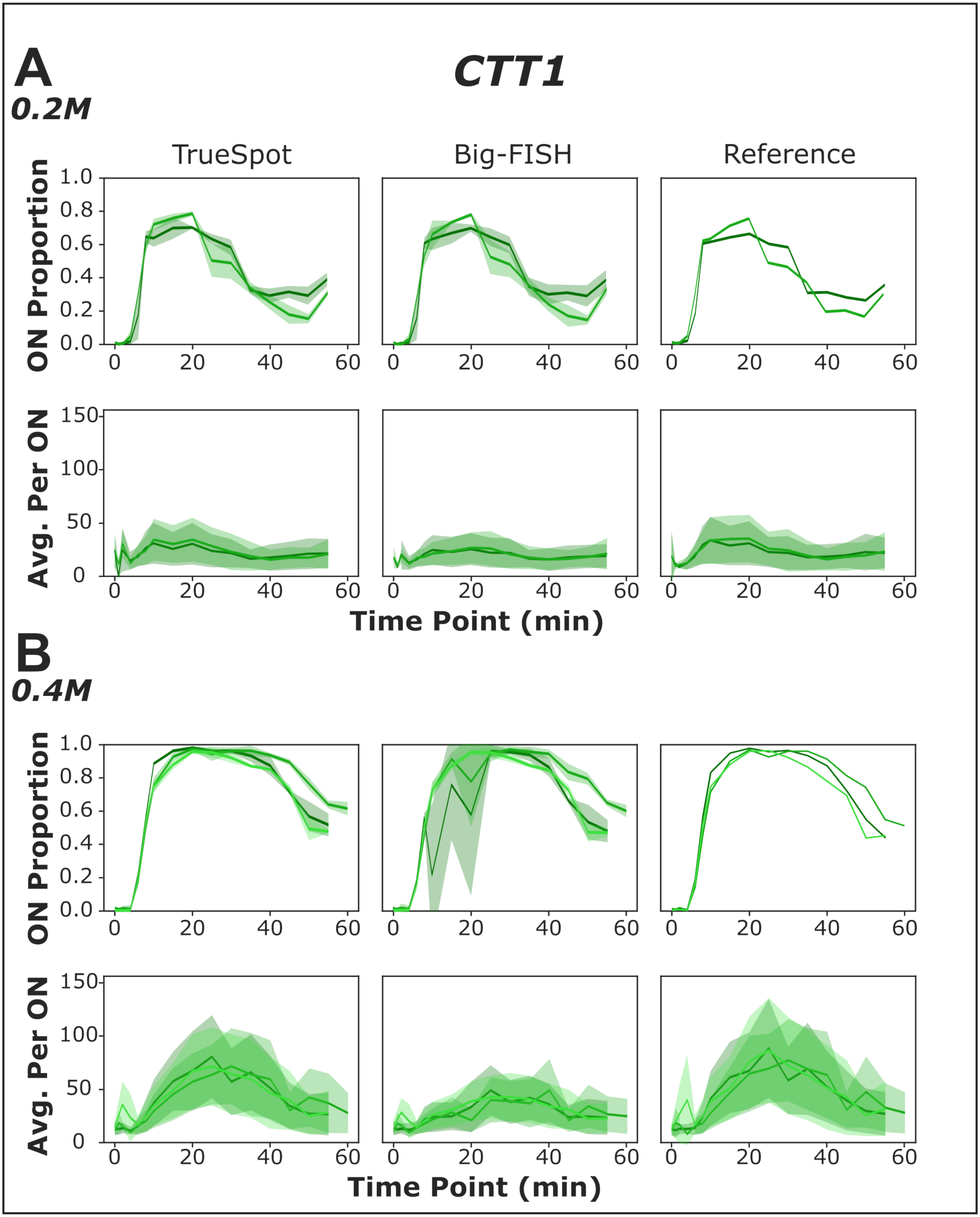
A
CTT1
0.2M
TrueSpot
Big-FISH
Reference
ON Proportion
1.0
0.8
0.6
0.4
0.2
0.0
0
20
40
60
Avg. Per ON
150
100
50
0
Time Point (min)
B
0.4M

1

**Figure S20. TrueSpot and Big-FISH can replicate CTT1 time course curves derived from manual thresholding.** (A) Measurements of ON proportion of total cells and (B) average transcripts per ON cell derived from analysis of experimental time course images tracking CTT1-CY5 expression upon osmotic stress induction. Comparison of TrueSpot and Big-FISH fully automated spot detection pipelines (using the same cell segmentation mask from TrueSpot) with measurements from previously published data using manual thresholding with a prototype of TrueSpot ("Reference"). Each line represents a biological replicate (0.2M n = 2, 0.4M n = 3), mean value across technical replicates (ON proportion) or all ON cells (Average per ON). Shaded areas are mean ± standard deviation of technical replicates (individual images within biological replicates, n = 3-9) at each time point.

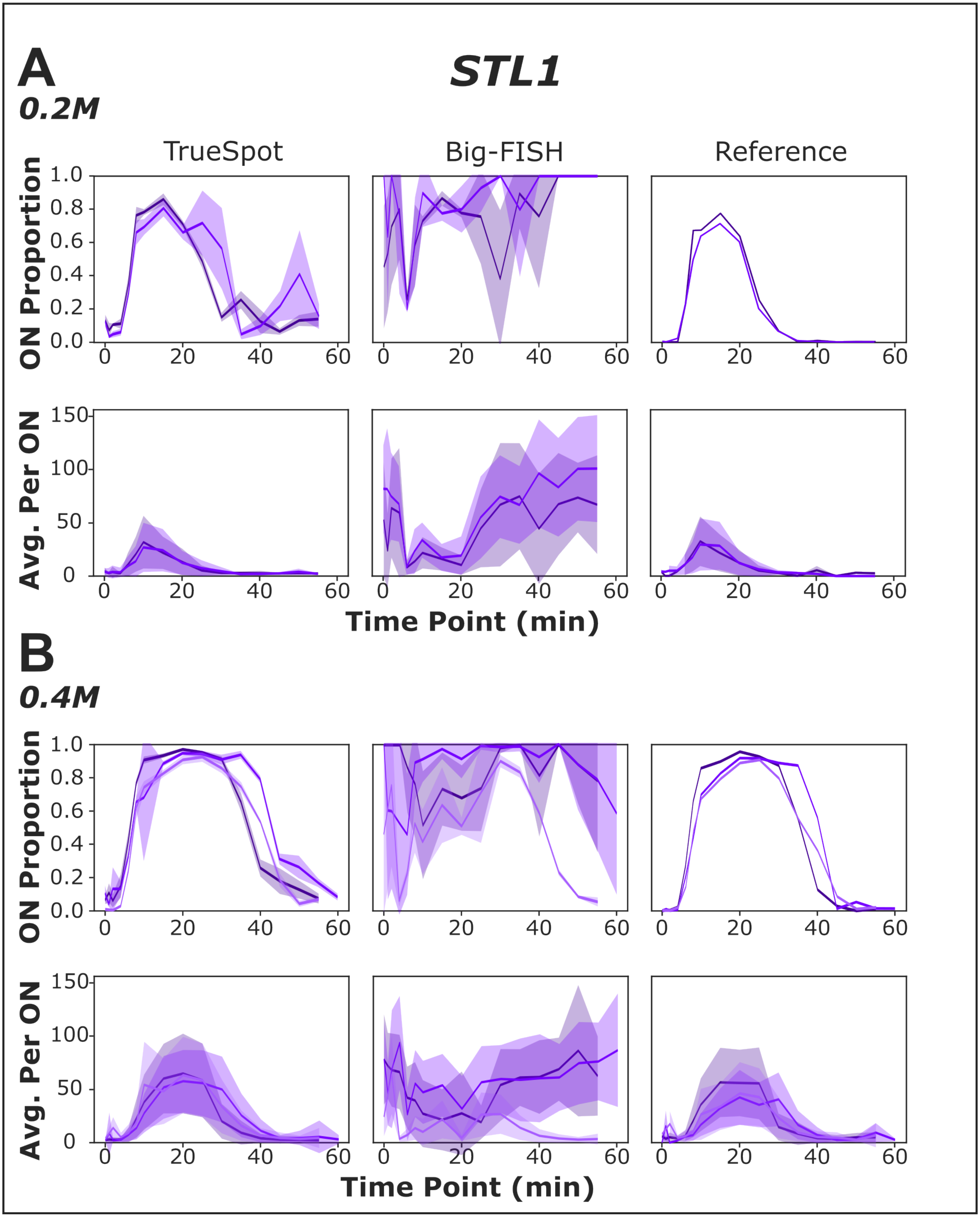
A
STL1
0.2M
TrueSpot
Big-FISH
Reference
ON Proportion
Avg. Per ON
Time Point (min)
B
0.4M
ON Proportion
Avg. Per ON
Time Point (min)

1

2

**Figure S21. TrueSpot can replicate STL1 time course curves derived from manual thresholding, but Big-FISH cannot.** (A) Measurements of ON proportion of total cells and (B) average transcripts per ON cell derived from analysis of experimental time course images tracking STL1-TMR expression upon osmotic stress induction. Comparison of TrueSpot and Big-FISH fully automated spot detection pipelines (using the same cell segmentation mask from TrueSpot) with measurements from previously published data using manual thresholding with a prototype of TrueSpot ("Reference"). Each line represents a biological replicate (0.2M n = 2, 0.4M n = 3), mean value across technical replicates (ON proportion) or all ON cells (Average per ON). Shaded areas are mean ± standard deviation of technical replicates (n = 3-9) at each time point.

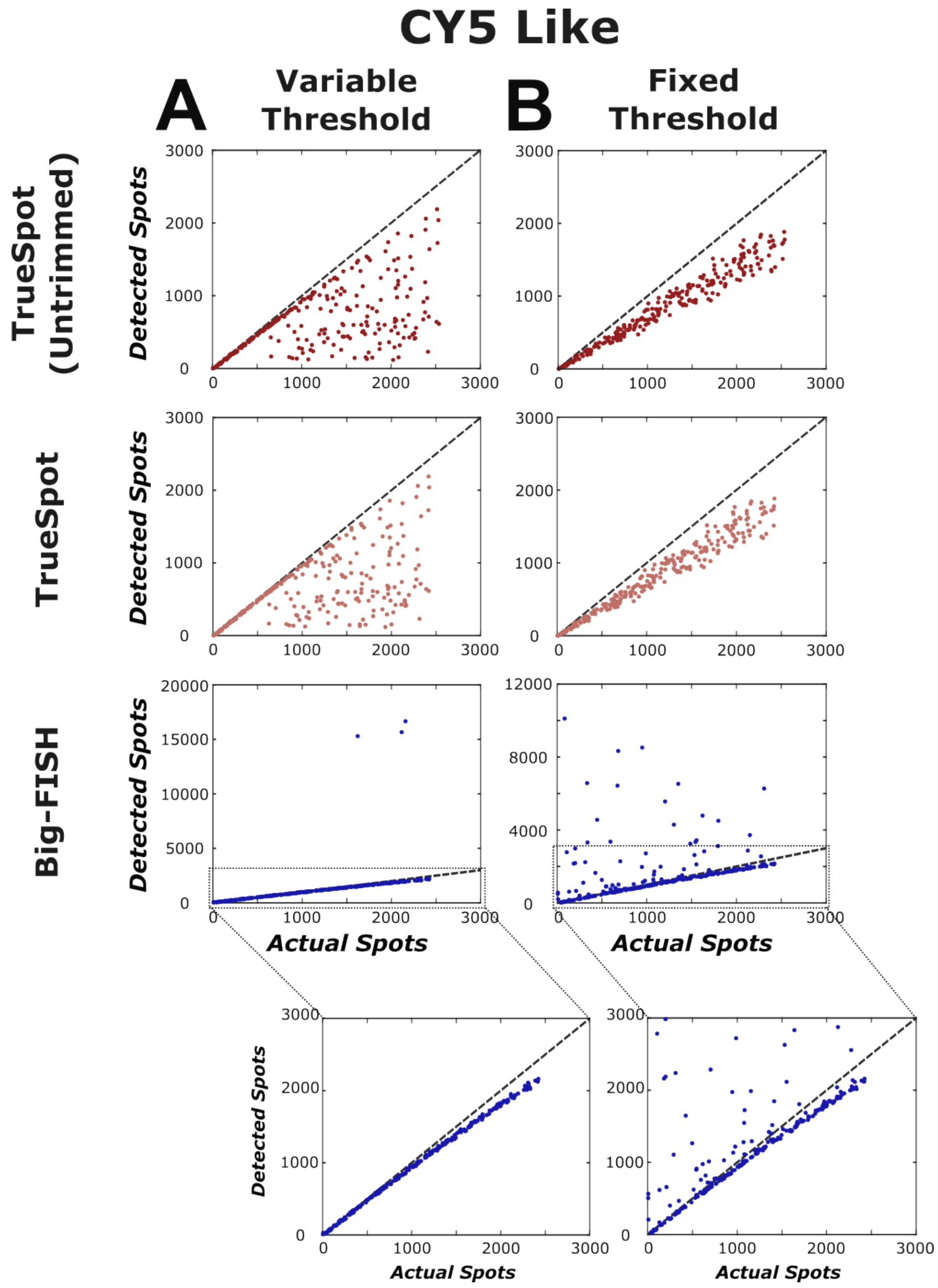


**Figure S22. Using a fixed threshold across similar CY5-Like simulated images does not necessarily improve performance over image specific thresholds.** Scatter plots of tool detected versus actual (ground truth) spot counts for all CY5-Like simulated images (n = 250) at (A) per-image auto-selected thresholds and (B) mean of per-image auto-selected thresholds (per tool) applied as a fixed threshold to all images. Dashed lines represent x = y. Bottom insets show y-scale zoomed-in views of Big-FISH plots.

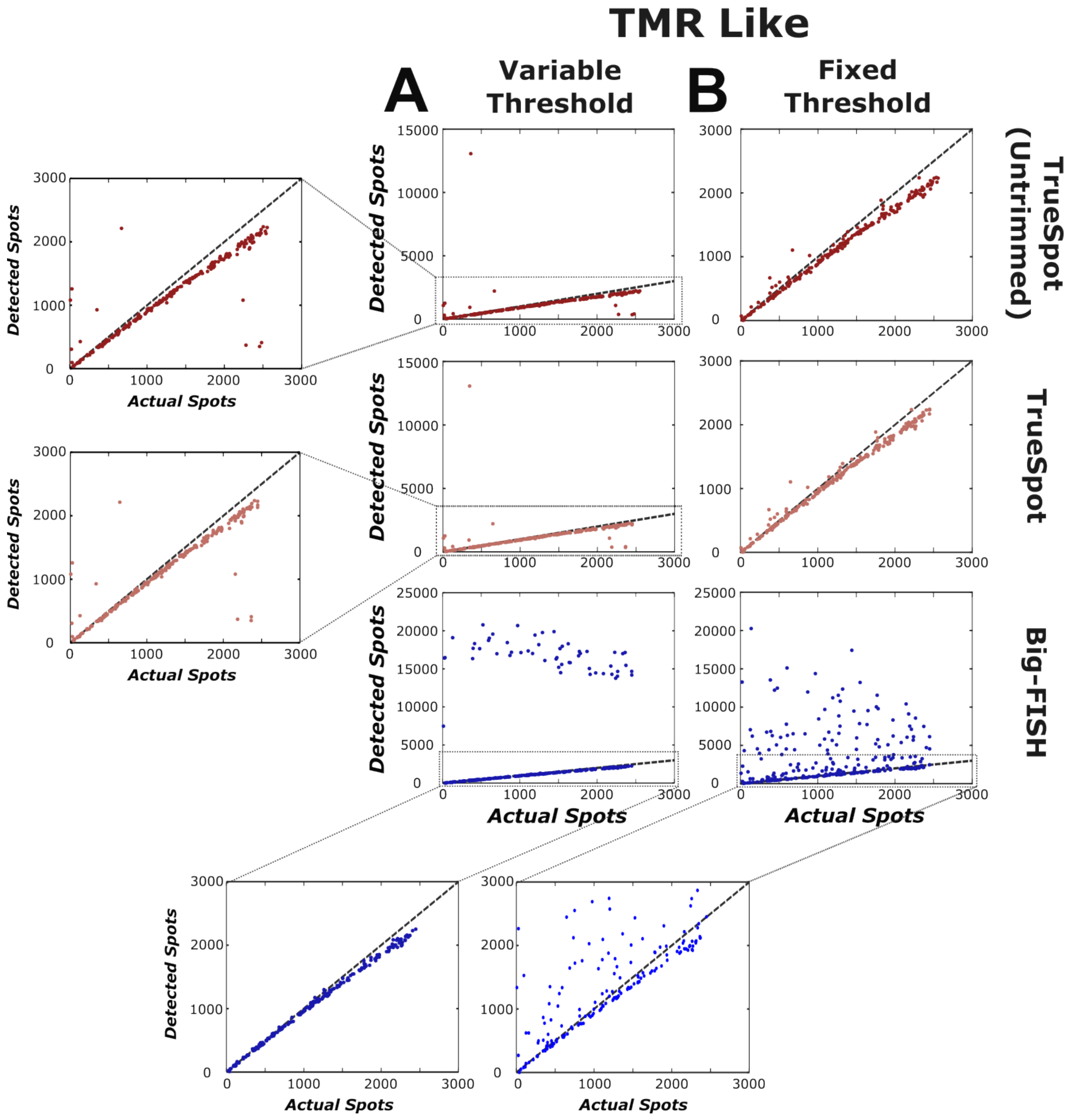


**Figure S23. Using a fixed threshold across similar TMR-Like simulated images does not necessarily improve performance over image specific thresholds.** Scatter plots of tool detected versus actual (ground truth) spot counts for all TMR-Like simulated images (n = 250) at (A) per-image auto-selected thresholds and (B) mean of per-image auto-selected thresholds (per tool) applied as a fixed threshold to all images. Dashed lines represent x = y. Bottom insets show y-scale zoomed-in views of Big-FISH plots, left insets show y-scale zoomed-in views of variable threshold TrueSpot (untrimmed and trimmed) plots.

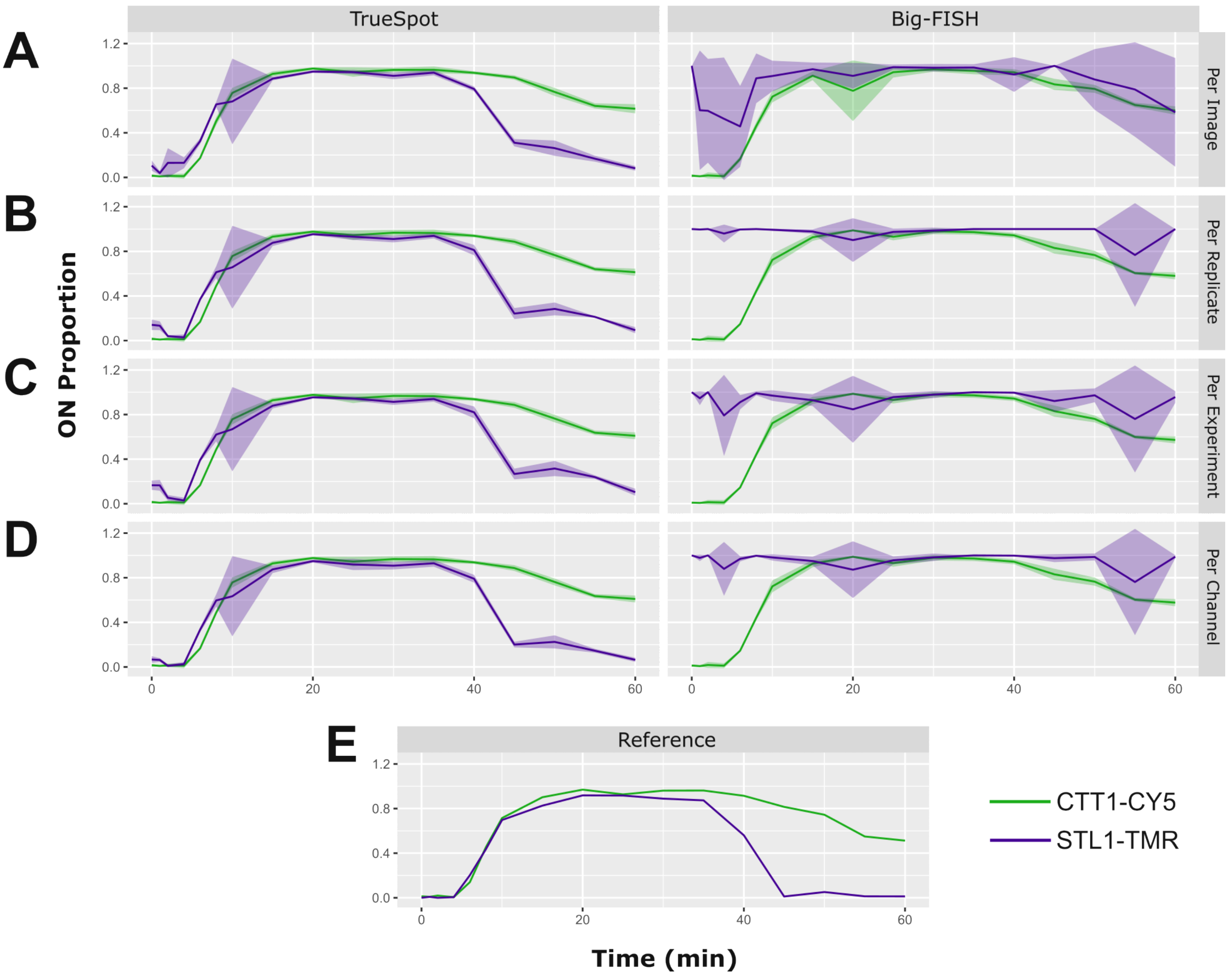


**Figure S24. Using group fixed thresholds over per-image thresholds may produce modest smoothing, but does not necessarily improve accuracy of ON proportion prediction.** ON cell proportion across CTT1 (green) and STL1 (purple) osmotic stress time courses (0.4M) for one biological replicate (E2R2) derived using different threshold selection approaches. Lines represent mean, shaded areas represent mean ± standard deviation across technical replicates (n = 3 - 9 per time point). Plots shown for both tools with automatic threshold selection (TrueSpot (left) and Big-FISH (right)). Threshold types tested are (A) per-image automatically selected ("Per Image"), (B) mean of per-image selections across biological replicate as fixed threshold ("Per Replicate"), (C) mean of per-image selections across experiment (salt concentration) as fixed threshold ("Per Experiment"), and (D) mean of per-image selections across all images in full experiment set as fixed threshold ("Per Channel"). (E) Published reference derived from manual channel-fixed threshold selection and a prototype of TrueSpot.

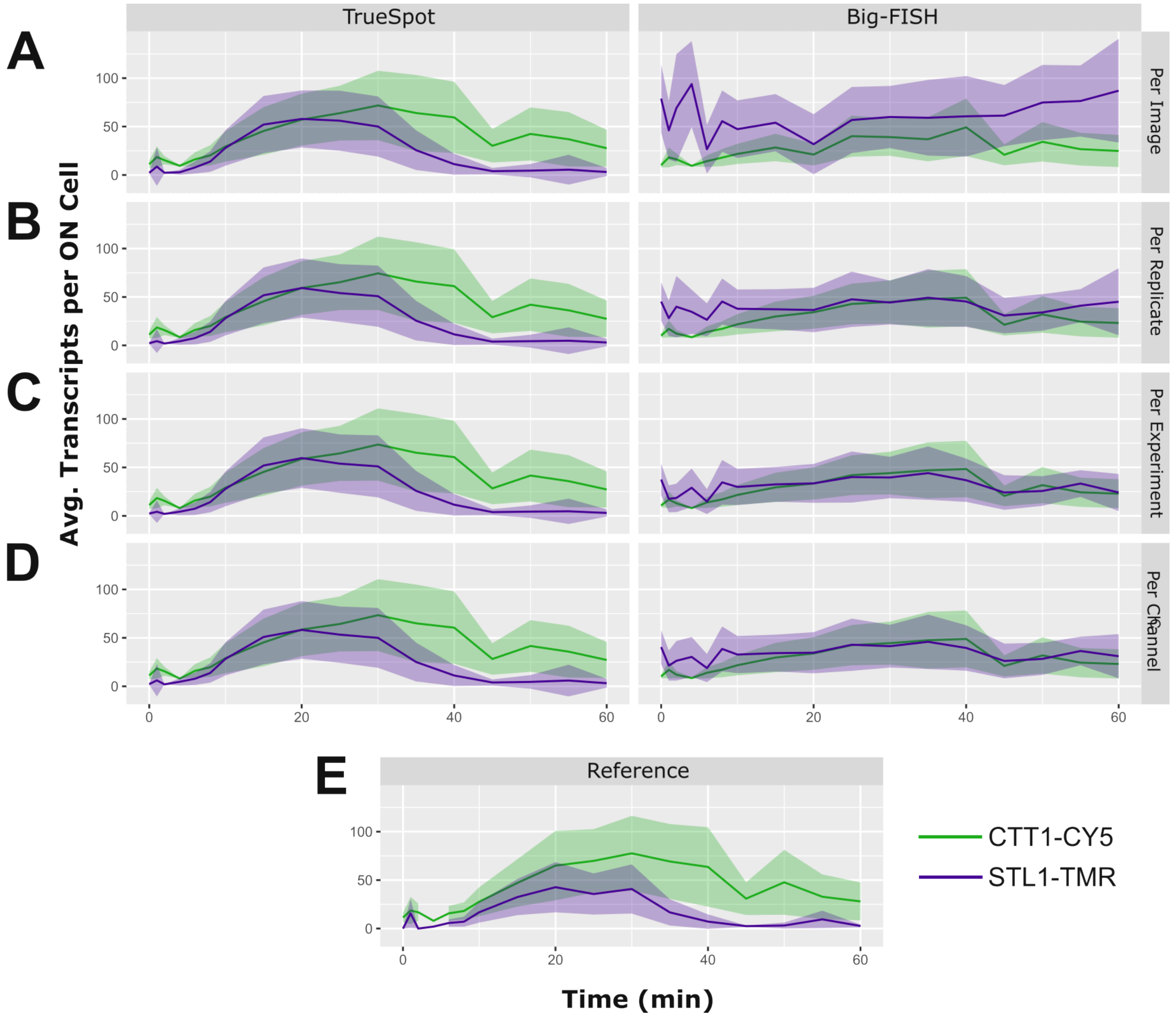


**Figure S25. Using group fixed thresholds over per-image thresholds can provide a very modest degree of smoothing, but does not necessarily improve accuracy of average transcript count per ON cell prediction.** Average transcripts per ON cell across CTT1 (green) and STL1 (purple) osmotic stress time courses (0.4M) for one biological replicate (E2R2) derived using different threshold selection approaches. Lines represent mean, shaded areas represent mean ± standard deviation across all cells in all technical replicates. Plots shown for both tools with automatic threshold selection (TrueSpot (left) and Big-FISH (right)) and published reference derived from manual channel-fixed threshold selection and a prototype of TrueSpot (bottom). Threshold types tested are (A) per-image automatically selected ("Per Image"), (B) mean of per-image selections across biological replicate as fixed threshold ("Per Replicate"), (C) mean of per-image selections across experiment (salt concentration) as fixed threshold ("Per Experiment"), and (D) mean of per-image selections across all images in full experiment set as fixed threshold ("Per Channel"). (E) Published reference derived from manual channel-fixed threshold selection and a prototype of TrueSpot.

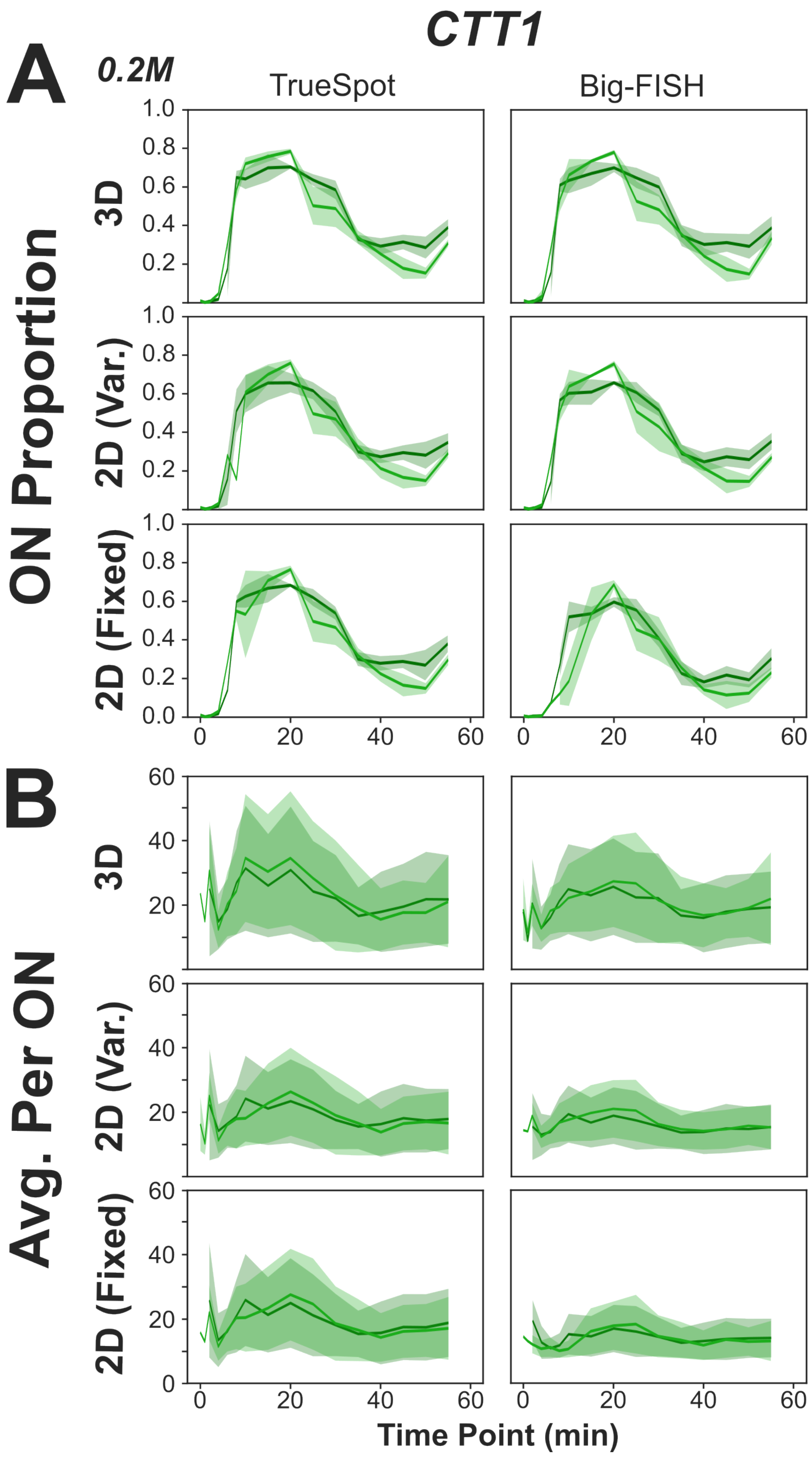
CTT1
A
0.2M
TrueSpot
Big-FISH
ON Proportion
3D
2D (Var.)
2D (Fixed)
B
Avg. Per ON
3D
2D (Var.)
2D (Fixed)
Time Point (min)

1

**Figure S26. Use of 2D maximum intensity projections depresses measured signal.** Calculated proportion of "on" cells (≥ 8 transcripts/spots) and average transcripts/spots per "on" cell for each time point from the 0.2M CTT1 time course. Measurements derived from TrueSpot on the left, Big-FISH on the right. Lines represent mean, filled regions represent standard deviation for each of 2 biological replicates. Full stacks were used. Results using both fixed and automatically determined (variable) thresholds are shown for maximum intensity projections ("2D Fixed" and "2D Var.").

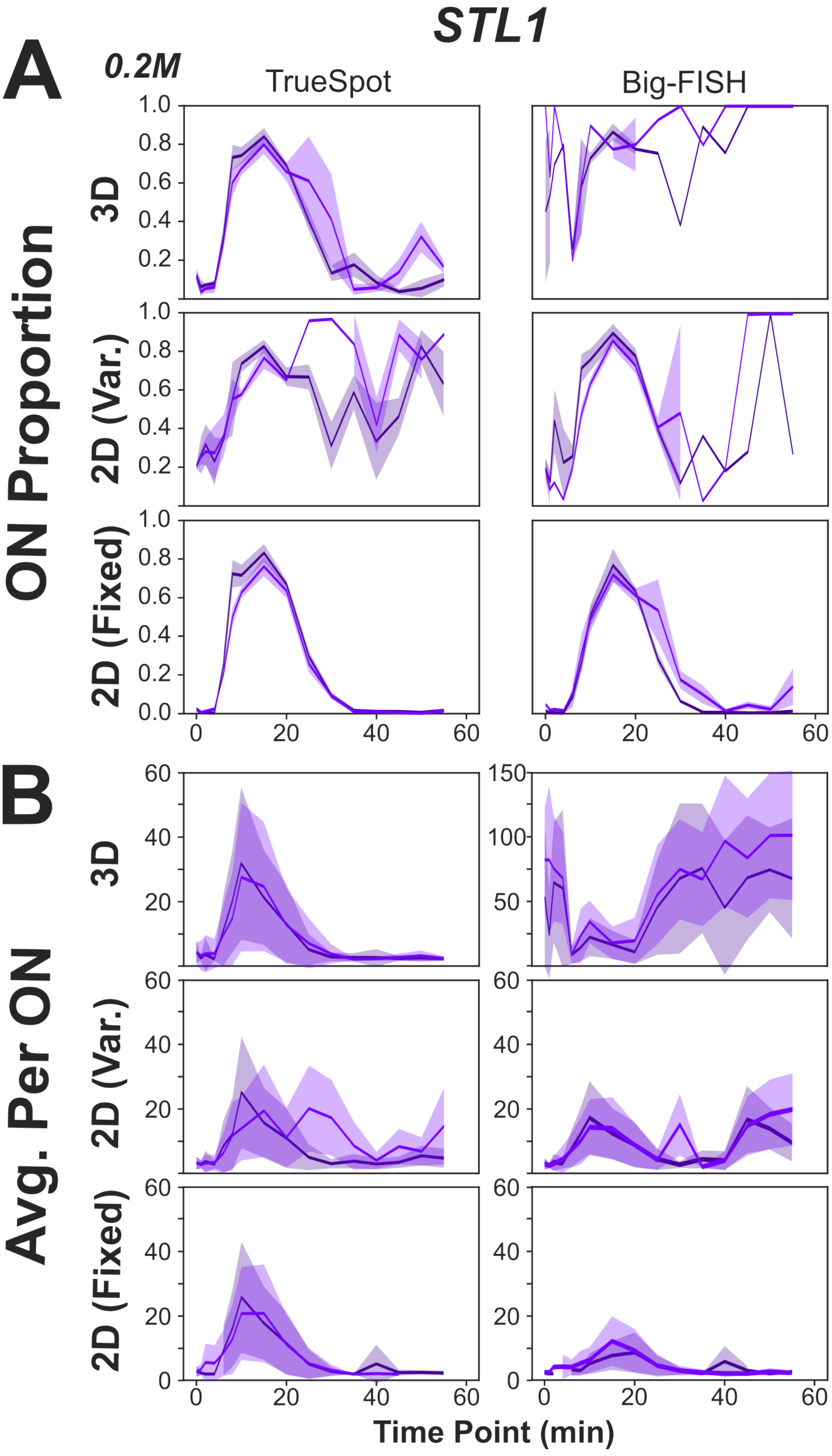

STL1
A
0.2M
TrueSpot
Big-FISH
ON Proportion
3D
2D (Var.)
2D (Fixed)
B
Avg. Per ON
Time Point (min)


1

**Figure S27. Use of 2D maximum intensity projections affects automatic threshold selection behavior.** Calculated proportion of “on” cells (≥ 2 transcripts/spots) and average transcripts/spots per “on” cell for each time point from the 0.2M STL1 time course. Measurements derived from TrueSpot on the left, Big-FISH on the right. Lines represent mean, filled regions represent standard deviation for each of 2 biological replicates. Full stacks were used. Results using both fixed and automatically determined (variable) thresholds are shown for maximum intensity projections (“2D Fixed” and “2D Var.”).

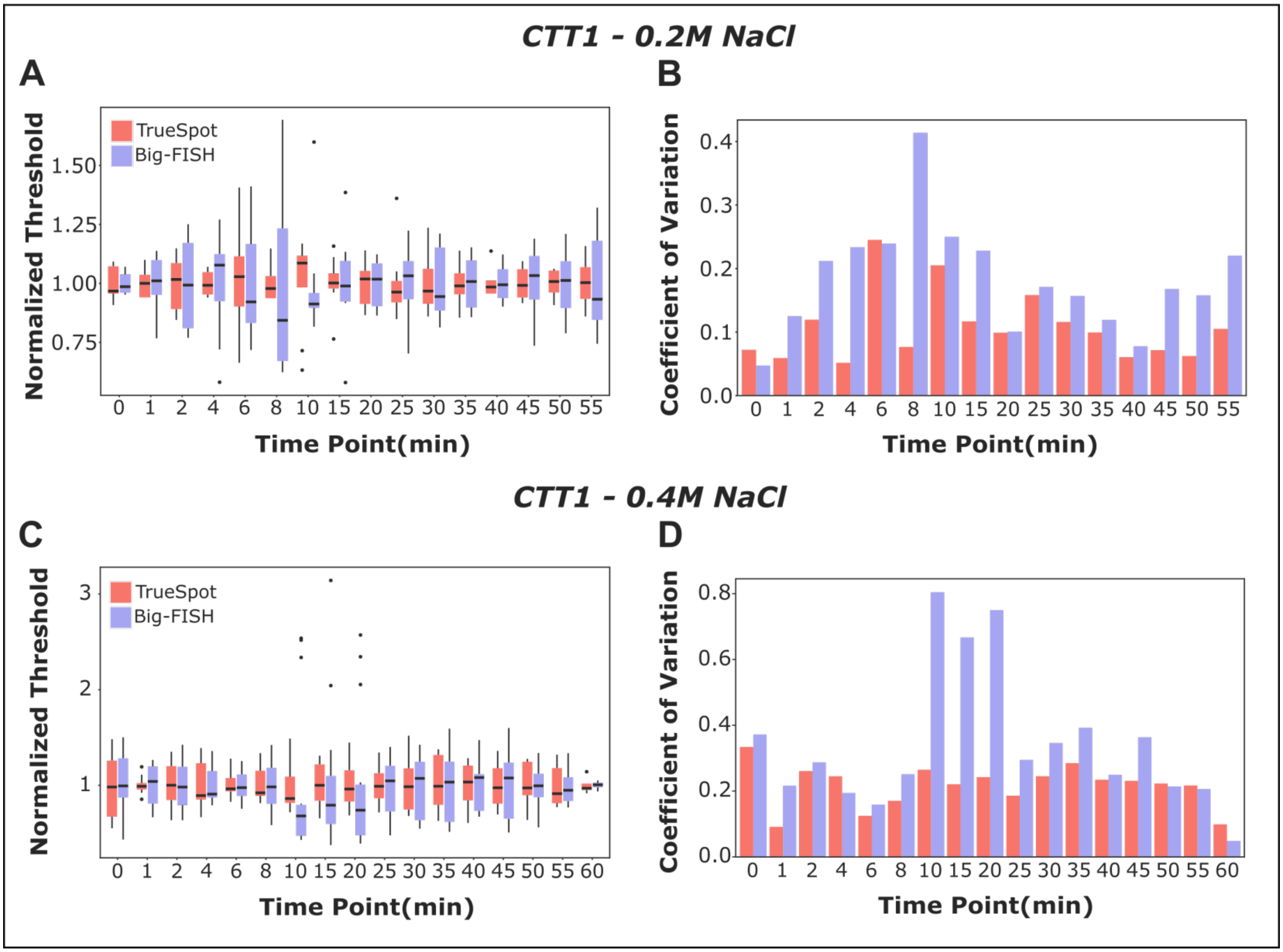


**Figure S28. TrueSpot and Big-FISH vary similarly in threshold selection at most CTT1 timepoints.** Variation in threshold selection across CTT1-CY5 osmotic stress time course images, grouped by time point and experiment by tools with automated threshold selection (TrueSpot (red), Big-FISH (blue)). (A-B) Experiment with maximum salt concentration of 0.2M, (C-D) experiment with maximum salt concentration of 0.4M. (A,C) Distribution of selected thresholds normalized to batch average (box bounds and midline represent 25th, 50th, and 75th percentiles, whiskers extend to further values within 1.5 * IQR from box edges, spots represent outliers). (B,D) coefficient of variation (standard deviation / mean) for each batch. Statistics are shown in Table S10.

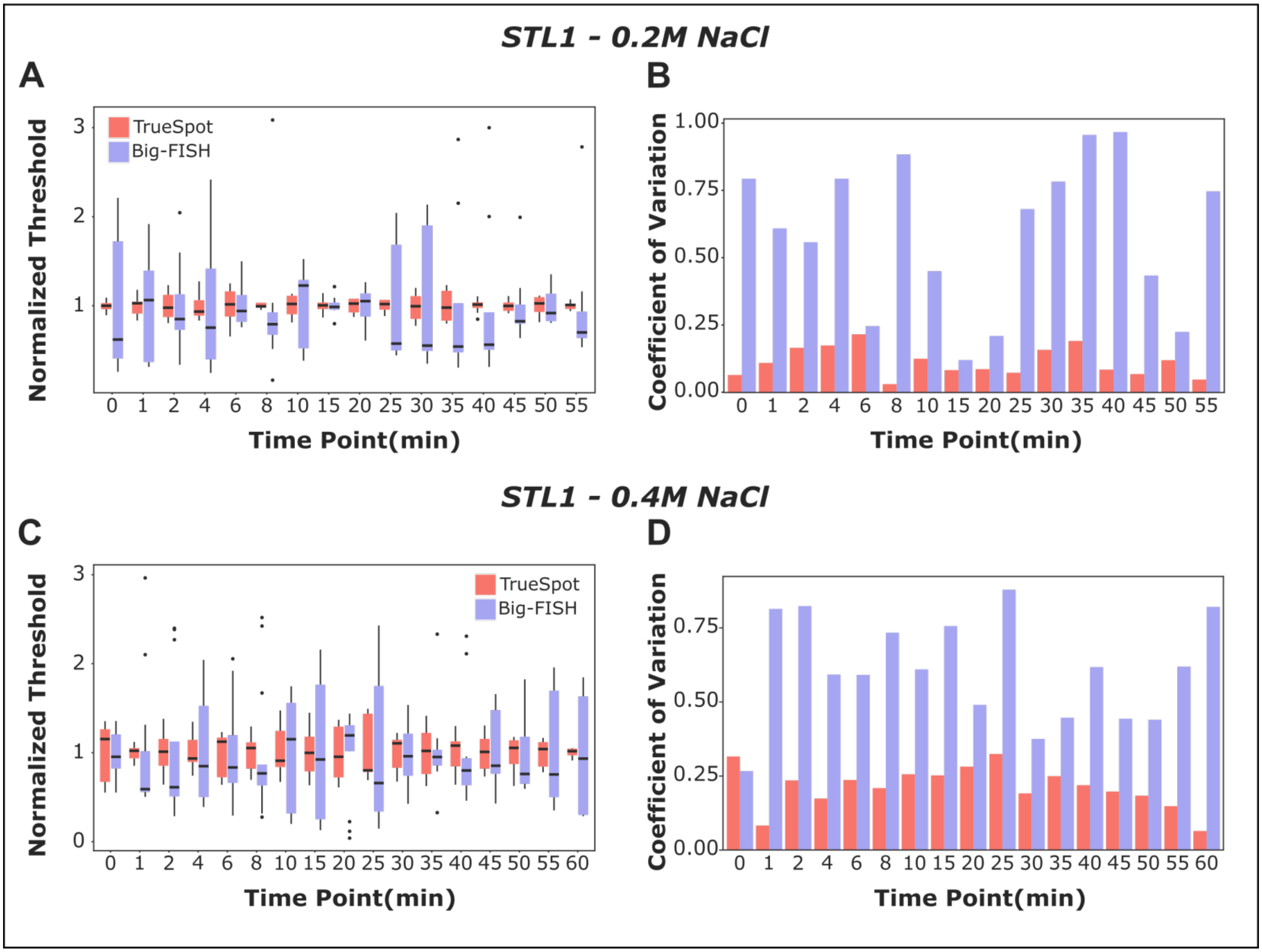


**Figure S29. TrueSpot tends to vary far less in threshold selection than Big-FISH within STL1 time points.** Variation in threshold selection across STL1-TMR osmotic stress time course images, grouped by time point and experiment by tools with automated threshold selection (TrueSpot (red), Big-FISH (blue)). (A-B) Experiment with maximum salt concentration of 0.2M, (C-D) experiment with maximum salt concentration of 0.4M. (A,C) Distribution of selected thresholds normalized to batch average (box bounds and midline represent 25$^{th}$, 50$^{th}$, and 75$^{th}$ percentiles, whiskers extend to further values within 1.5 * IQR from box edges, spots represent outliers). (B,D) coefficient of variation (standard deviation / mean) for each batch. Statistics are shown in Table S10.

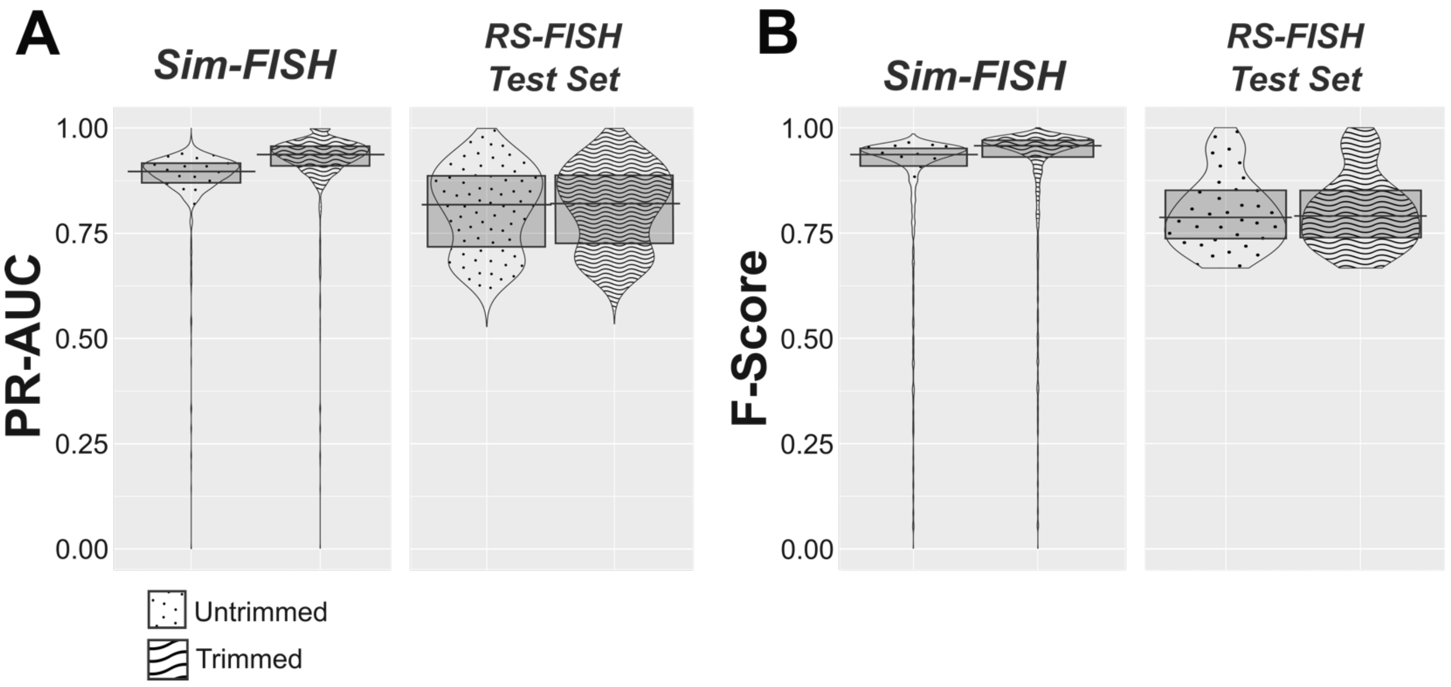


**Figure S30. Trimming out borders of a simulated image marginally improves performance score.** Distribution of TrueSpot (A) PR-AUC and (B) F-Scores for all simulated images (split by simulation tool) when edges of image are untrimmed versus trimmed (7 pixels – Gaussian filter radius - from each edge in x and y). Mann-Whitney test calculations are shown in Table S1. The same 7 pixel border trim was applied for all tools.

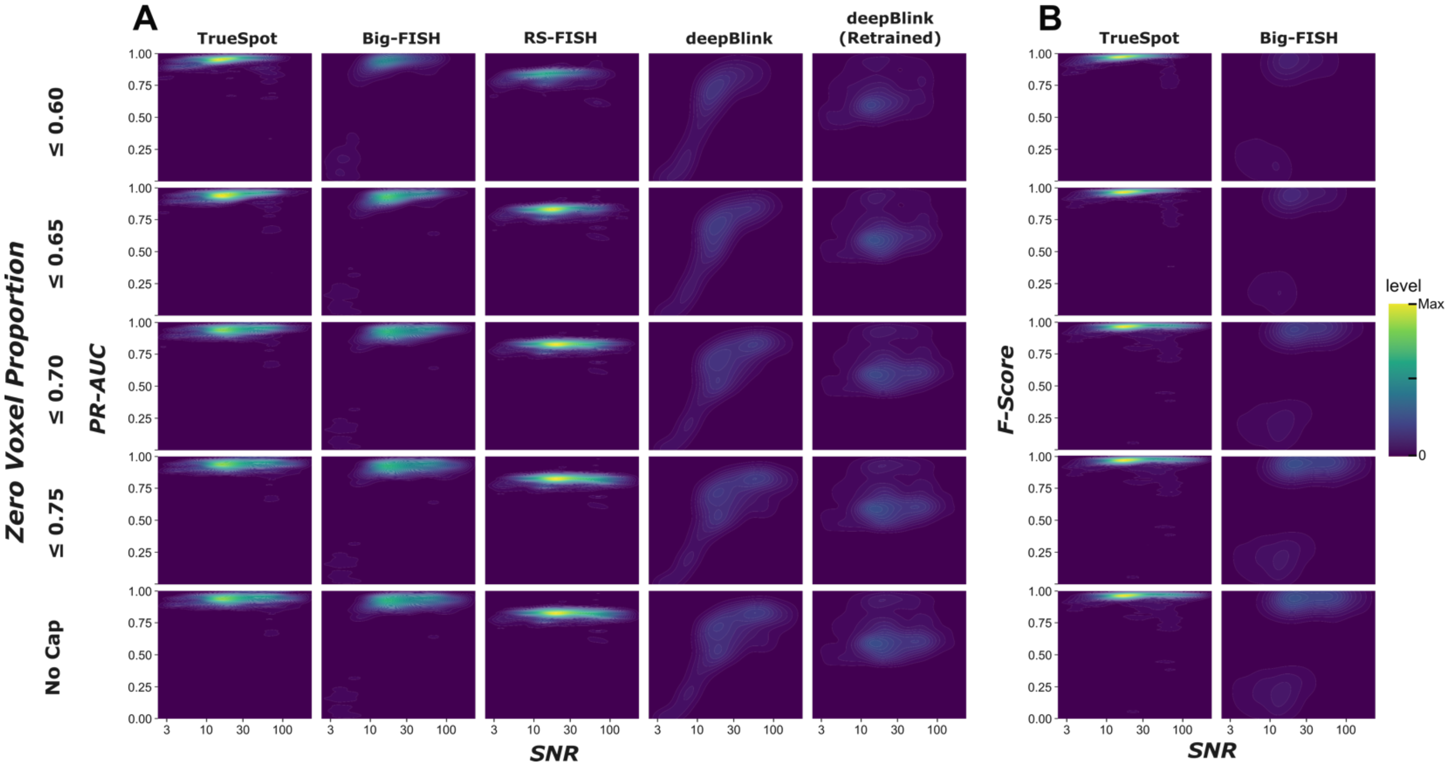


**Figure S31. ZVP filtering removes less realistic images without significantly altering overall results.** Density maps of (A) PR-AUC versus SNR (all four tools, two deepBlink models) and (B) F-Score (automatic thresholding tools only) are displayed for simulated image pools filtered using five ZVP caps (see Table S17 for n values and Spearman calculations). General score distributions remain roughly the same across ZVP caps for a given tool.

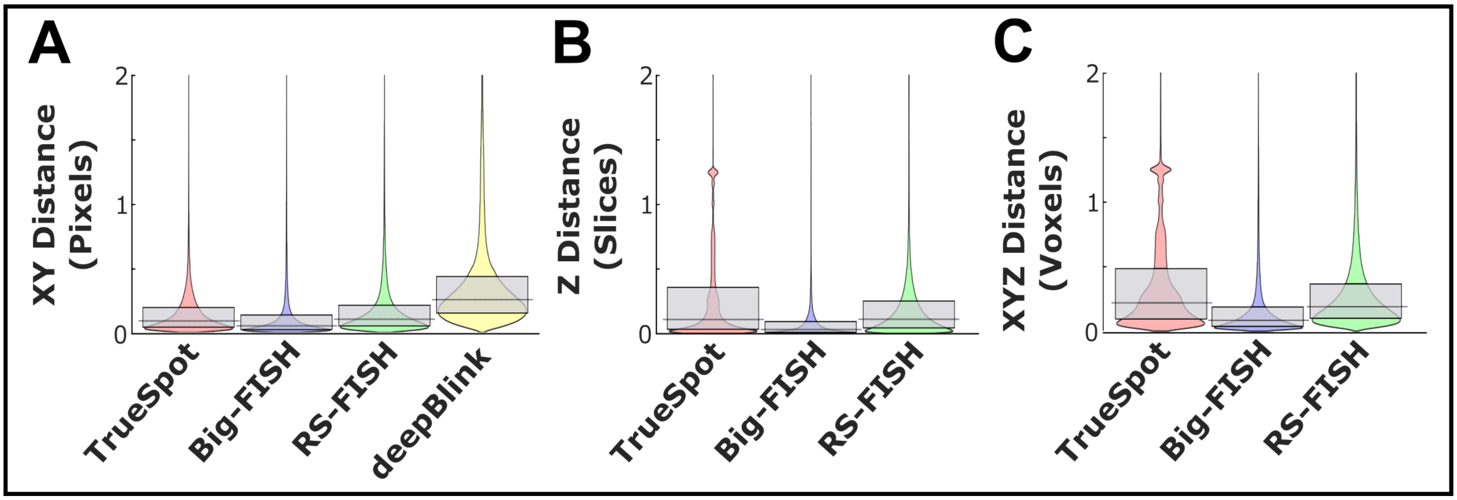


**Figure S32. Tools are able to localize simulated spots to within half a pixel in most cases.** Subpixel fit distance from ground truth spot coordinates (Sim-FISH generated images) in (A) the xy plane, (B) z only, and (C) xyz (3D). deepBlink (yellow) omitted from 3D evaluations (B-C) due to tool not operating across stack slices. Grey box marks 25$^{th}$ and 75$^{th}$ percentiles, black line marks median. Statistics can be found in Table S13.

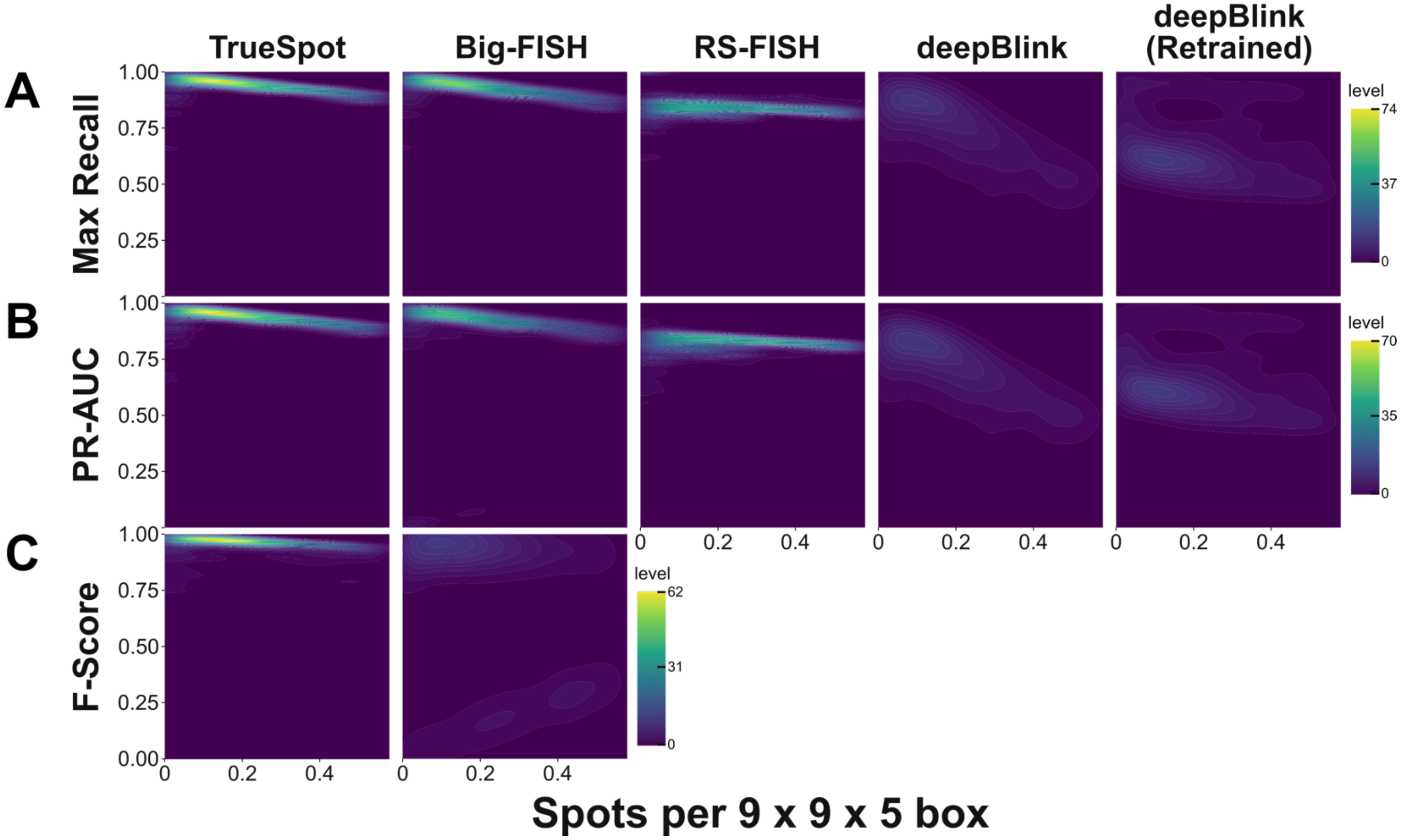


**Figure S33. Performance tends to decrease with spot density.** Density plots of (A) maximum recall, (B) PR-AUC, and (C) F-Score versus a measurement of spot density (ground truth spot count adjusted for image size). Spearman correlation values can be found in Table S15.

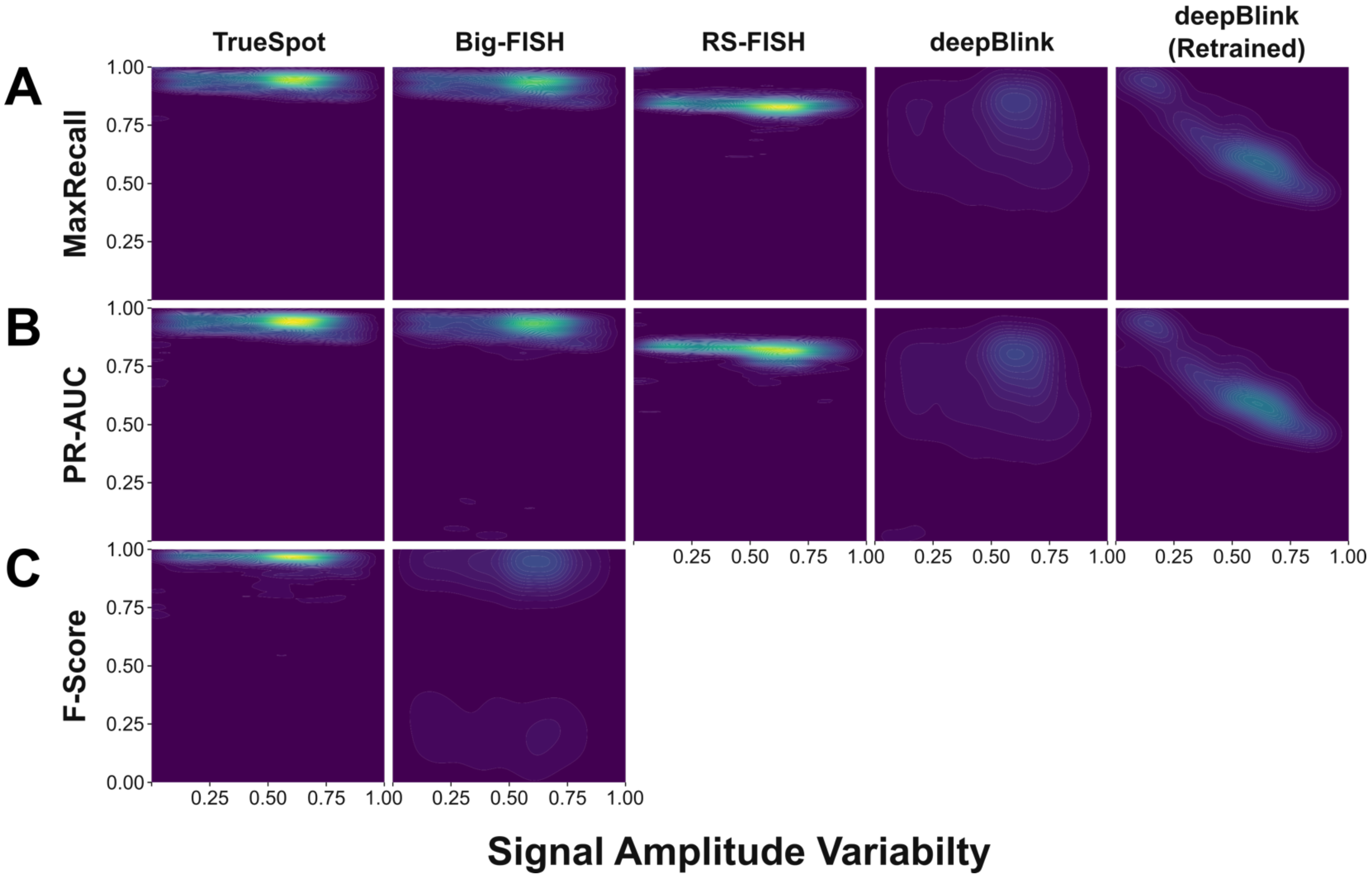


**Figure S34. Performance is usually unaffected by signal variability.** Density plots of (A) maximum recall, (B) PR-AUC, and (C) F-Score versus signal amplitude variability (simulation input parameter, proportion of mean signal amplitude). Correlation is strongest in the case where deepBlink was used provided a model trained on a subset of the simulated data (Spearman: Max Recall Estimate = -0.769, p = $2.19 \times 10^{-270}$; PR-AUC Estimate = -0.770, p = $8.27 \times 10^{-272}$). Spearman calculations are shown in Table S16.

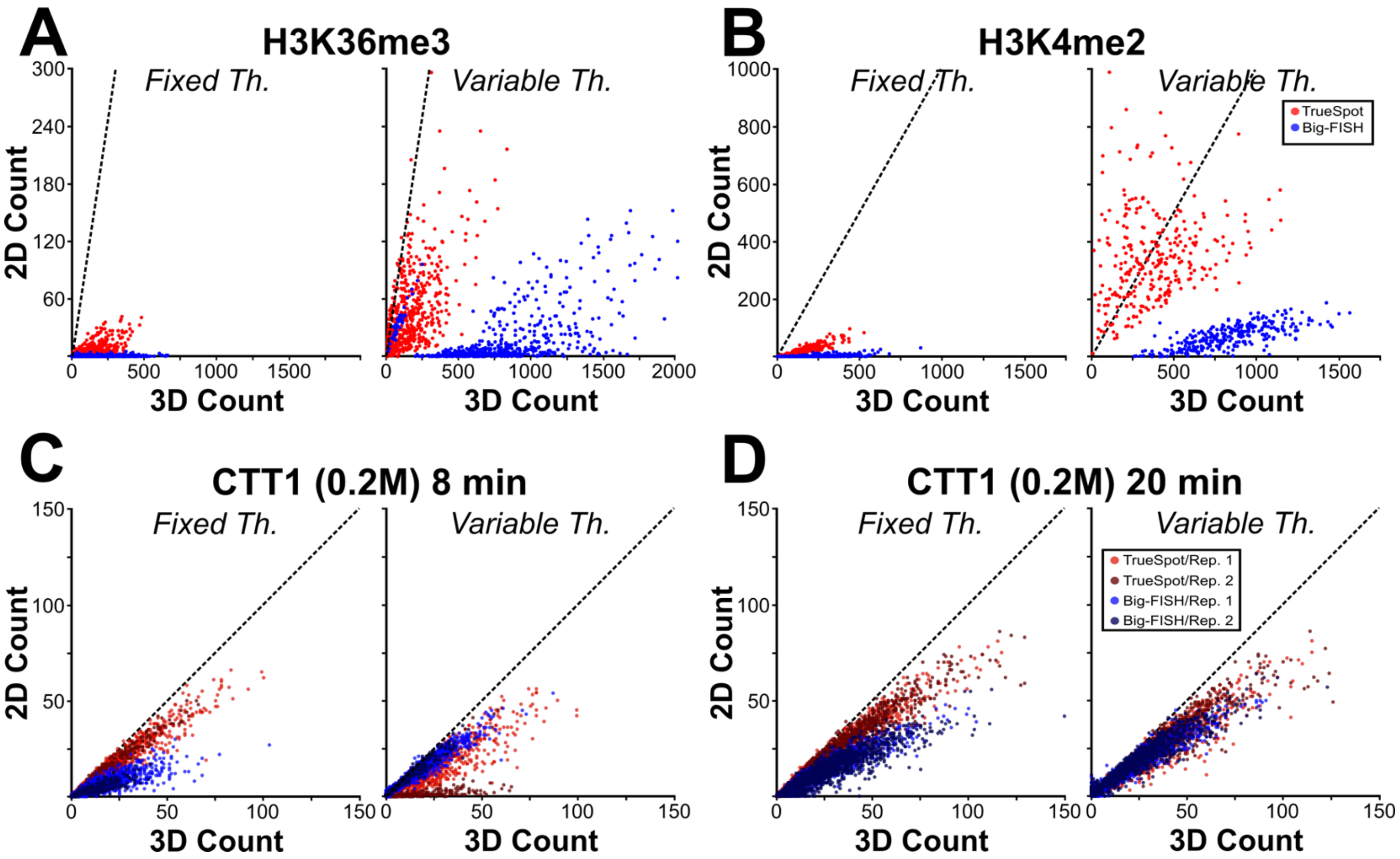


**Figure S35. Use of automated threshold selection alters per-cell count distribution relative to fixed thresholding.** Scatterplots of per-cell spot count in 2D (from full stack maximum intensity projections) versus 3D for histone groups (A-B) and two CTT1 0.2M time points (C-D). Distributions derived from using fixed threshold values for each tool are compared to distributions using tools' built-in automated threshold selection. Both biological replicates are shown for CTT1 0.2M. Each point represents a single cell. Fill color reflects tool and replicate.

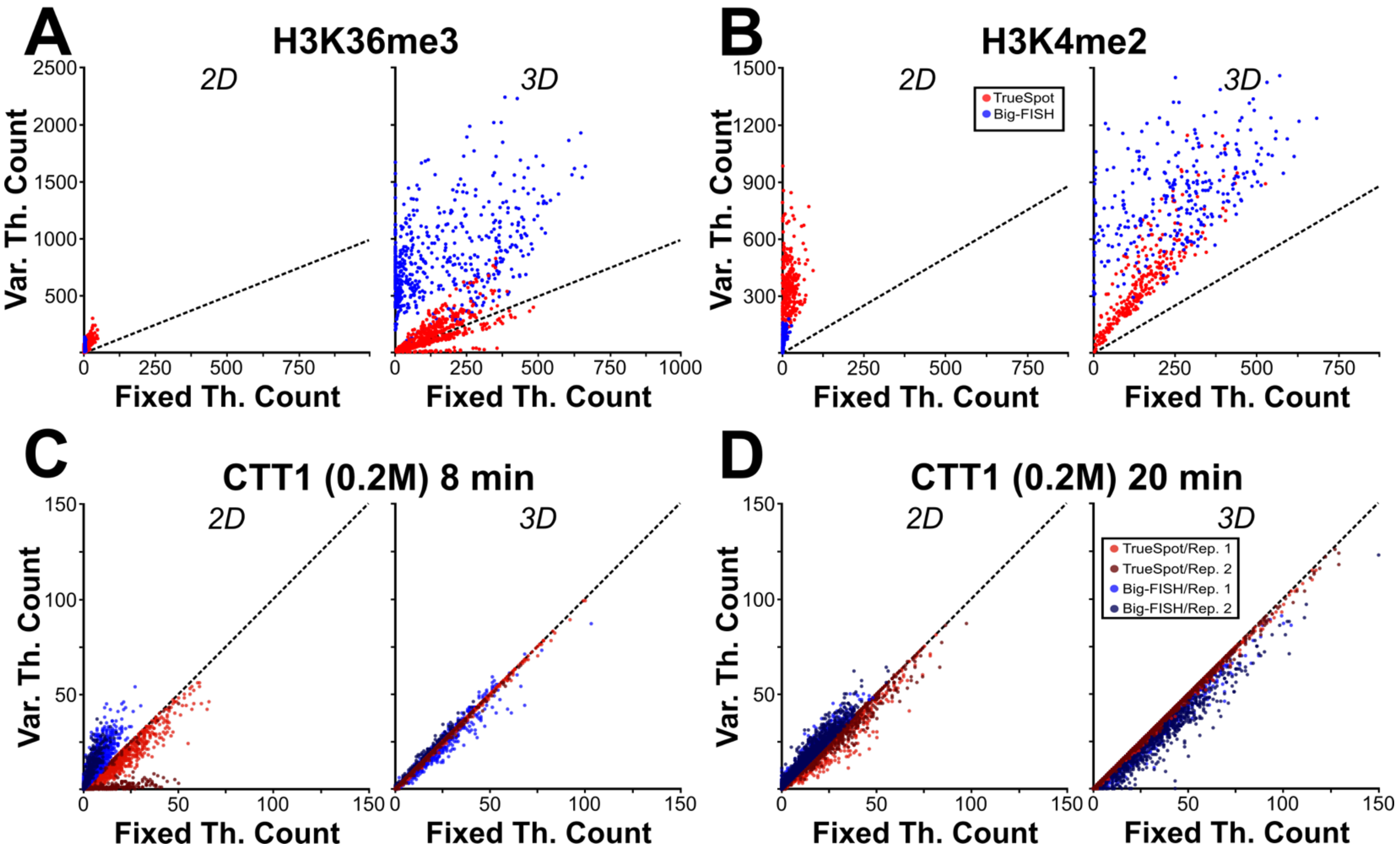


**Figure S36. Correlation between fixed threshold and variable threshold counts is higher when using full 3D stacks than when using maximum intensity projections.** Scatterplots of per-cell spot counts using fixed versus variable thresholds for both 2D maximum intensity projections and 3D full stacks. Results for both histone groups (A-B) and two CTT1 0.2M time points (C-D) are shown. Both biological replicates are shown for CTT1 0.2M. Each point represents a single cell. Fill color reflects tool and replicate.

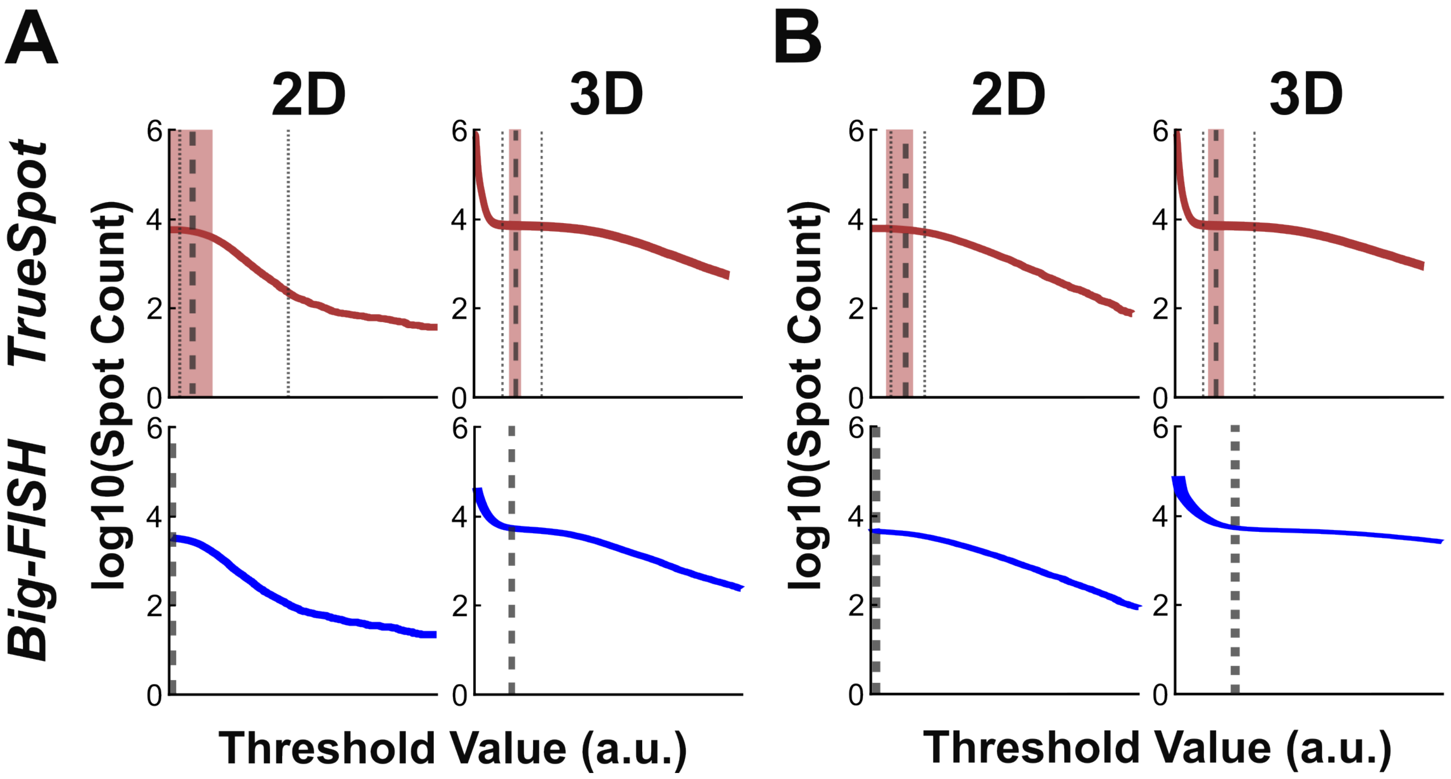


**Figure S37. Maximum intensity projections may produce spot count curves with no background noise dropoff.** Examples of log10 scaled spot count curves from two sample 0.2M STL1 time course images at 8 min (A) and 15 min (B) from both 2D maximum intensity projections (left) and 3D full stacks(right). Dashed lines denoted automatically selected thresholds. TrueSpot red range represents threshold suggestion range mean ± one standard deviation and dotted lines represent the extremes of suggestion range.

### Supplementary Tables

**Table S1. Tool performance on two sets of simulated images was assessed.** Mean, standard deviation, and quartiles of performance metric distributions (i) as well as Mann-Whitney comparisons between tools (ii) are shown for the metrics of (A) Maximum recall, (B) PR-AUC, and (C) F-Score at automatically selected threshold. Visual representations of results can be found in Figures 3A-B and S6A. Simulated image sets are denoted either “SF” (Sim-FISH) or “PL” (Preibisch Lab – RS-FISH).

**Table S2. Tool performance on two singletons and twelve sets of experimental images was assessed.** Mean, standard deviation, and quartiles of performance metric distributions (i) as well as Mann-Whitney comparisons between tools (ii) are shown for the metrics of (A) Maximum recall, (B) PR-AUC, and (C) F-Score at automatically selected threshold. Visual representations of results can be found in Figures 3C-D and S9.

**Table S3. TrueSpot has many parameter presets.** Parameter values provided to automatic thresholding module for 17 presets numbered from most precise to most sensitive. Values used for individual images can be found in Table S12. Window size options specify Fano factor scan window size. Values can be integers, which indicate a fixed size, or floats representing a proportion of the full threshold scan range. “Log Mode” specifies when log10 transformation is applied. If “all”, then it is applied immediately, before first derivative approximation is taken. If “fit”, the transformation is only applied for 2-piece linear fit. Weights specify the weighted average multipliers for each suggestion pool toward the final suggestion. MAD factors specify the range of factors to multiply the MAD by to search for median/MAD suggestions. “Add St. Devs” is the number of standard deviations of the suggestion pool to add to the pool weighted average to get the final suggestion. “Reweight Fit” and “Spline Iterations” were unused experimental parameters and were eventually deprecated.

**Table S4. Per-cell spot count correlation between TrueSpot and Big-FISH varies across image groups.** Pearson estimates, Spearman estimates, and linear regression parameters for each subgroup dataset (biological replicate for *S. cerevisiae* time courses) shown in Figures 4B, 4D, S15, and S17.

**Table S5. Per-cell spot counts from 2D maximum intensity projections and 3D stacks rarely agree.** Linear regression parameters with Pearson and Spearman estimates for all tested groups and time course time points. Values are shown for all tested maximum intensity generation and comparison parameters for both TrueSpot and Big-FISH.

**Table S6. Group fixed thresholds were derived from mean of group auto-selected thresholds.** Values used as fixed thresholds for time course and simulated time course groups. For experimental images, grouping was performed across biological replicate, experiment (salt concentration), and full channel. Simulated images were only grouped by batch. Fixed thresholds for mESC image channels were derived from the mean of the threshold values at peak F-Score for the subset of images with reference sets.

**Table S7. Automated tools can replicate RNA counts derived from manual threshold selection.** By time-point comparisons of ON proportion and average RNA per ON cell curves (paired t-test) of each biological replicate, for each tool to manually derived reference values. Curves derived from variable per-image thresholding and fixed threshold selections are shown. Visual representations of results can be found in Figures 5, S20, and S21.

**Table S8. Fixing thresholds for simulated image groups affects correlation between counts of ground truth and detected spots.** Spearman and Pearson correlation values for actual versus detected spots when thresholds are either per-image variable or fixed to batch mean. (A) CY5L simulated image set, (B) TMRL simulated image set. Visual representations of results can be found in Figures S22 and S23.

**Table S9. Fixing thresholds does not change final RNA counts much relative to using per-image thresholding.** Comparisons by time-point (paired t-test) between RNA count curves (ON proportion and RNA per ON cell) of the same tool and biological replicate using different thresholding strategies. Visual representations of results can be found in Figures S24 and S25.

**Table S10. Distributions of automatically selected thresholds show more consistency for TrueSpot than for Big-FISH.** Counts (n), with mean, standard deviation, and coefficients of variation (standard deviation/mean) for tool selected threshold values within image batches. Visual representations of results can be found in Figures 6, S28, and S29.

**Table S11. Mass and small set simulated images were generated using Sim-FISH.** Parameter values used to generate each batch of images in Sim-FISH. Ranges are shown for mass simulated batches. Exact parameters for each individual images within the mass batches can be found in the result MAT files and sim_results.csv.

**Table S12. The test set included nearly 2500 simulated and experimental images.** Table of information for each test image in the set including name, group classification, stack file name, dimensions, voxel dimensions and metadata regarding the source of experimental images.

**Table S13. Subpixel fits were performed for untrimmed spots in simulated images.** Counts (n), mean, standard deviation, and quartiles, and Mann-Whitney statistics for subpixel fits by tool for (A) XY only, (B) Z only, (C) XYZ. deepBlink was not included in any 3D analyses.

**Table S14. The performance of some tools is more sensitive to SNR than others.** Spearman correlation values for SNR versus (A) Maximum recall, (B) PR-AUC, and (C) F-Score at auto selected threshold for set of all simulated images for all tools tested. Visual representations of results can be found in Figures S6B, S8 and S31.

**Table S15. The performance of most tools is anticorrelated with spot density.** Spearman correlation values for spot density measured as the number of ground truth spots per 9 x 9 x 5 box versus (A) Maximum recall, (B) PR-AUC, and (C) F-Score at auto selected threshold for set of all simulated images for all tools tested. Visual representations of results can be found in Figure S33.

**Table S16. Correlation between performance and amplitude variation is weak for most tools except the retrained deepBlink model.** Spearman correlation values for amplitude variability as input during simulated image generation versus (A) Maximum recall, (B) PR-AUC, and (C) F-Score at auto selected threshold for set of all simulated images for all tools tested. Visual representations of results can be found in Figure S34.

**Table S17. There are some weak correlations between zero voxel proportion and performance metrics.** (A) Spearman test values comparing the correlation between ZVP and various image properties and performance metrics. Performance metrics are split by tool. (B) Total count (n) of images at or below different ZVP ceilings. See Figure S5 for visual representation of some of the results in table.
